# RNA sequencing demonstrates *ex vivo* neocortical transcriptomic changes induced by epileptiform activity in male and female mice

**DOI:** 10.1101/2023.11.20.567411

**Authors:** Alec J. Vaughan, Laura J. McMeekin, Kutter Hine, Isaac W. Stubbs, Neela K. Codadu, Simon Cockell, Jonathon T. Hill, Rita Cowell, Andrew J Trevelyan, R. Ryley Parrish

## Abstract

Seizures are generally associated with epilepsy but may also be a symptom of many other neurological conditions. A hallmark of a seizure is the intensity of the local neuronal activation, which can drive large-scale gene transcription changes. Such changes in the transcriptional profile are likely to alter neuronal function, thereby contributing to the pathological process. Therefore, there is a strong clinical imperative to characterie how gene expression is changed by seizure activity. To this end, we developed a simplified *ex vivo* technique for studying seizure-induced transcriptional changes. We compared the RNA sequencing profile in mouse neocortical tissue that had up to 3 hours of epileptiform activity induced by 4-aminopyridine (4AP), relative to control brain slices not exposed to the drug. We identified over 100 genes with significantly altered expression after 4AP treatment, including multiple genes involved in MAPK, TNF, and neuroinflammatory signalling pathways, all of which have been linked to epilepsy previously. Notably, the patterns in male and female brain slices were almost identical. Various immediate early genes were among those showing the largest upregulation. The set of down-regulated genes included ones that might be expected either to increase or to decrease neuronal excitability. In summary, we found the seizure-induced transcriptional profile to be complex, but the changes aligned well with an analysis of published epilepsy-associated genes. We discuss how simple models may provide new angles for investigating seizure-induced transcriptional changes.

**Significance Statement:** It is well-established that strong neuronal activation results in large-scale transcriptomic changes. Understanding this process is of particular importance in epilepsy, which is characterized by paroxysmal pathological discharges. The complexity of *in vivo* activity patterns, however, present many difficulties for interpretation of the transcriptional changes. In contrast, *ex vivo* seizure models provide better experimental control and quantification of activity patterns, with lower welfare impact. Importantly, we now show that these models also replicate the transcriptional patterns previously reported in chronic human and animal epilepsy, thus validating their use in these kinds of study.

## Introduction

It has long been established that neuronal activity has a powerful influence on gene expression (Lee & Fields, 2021; Sheng et al., 1993). In turn, altered gene expression in neurons leads to changes in neuronal function, which constitutes an important form of feedback control within the brain, albeit mediated on a time-scale orders of magnitude slower than synaptic interactions. These slow transcriptional feedback mechanisms are, however, likely to play an important role in epileptic pathophysiology, given that seizures represent the most intense form of neuronal activation. It is important, therefore, to characterise exactly how acute seizure activity alters the transcriptomic landscape (Morgan et al., 1987). Gene expression changes that reduce neuronal network excitability can be characterised as negative feedback mechanisms, serving a protective, homeostatic function. In contrast, changes that increase neuronal excitability, depending where in the network they occur, can act as positive feedback mechanisms, leading to an escalation of the pathophysiology. Both sets of changes, however, are likely to alter the interictal functionality of the network, and thus contribute to co-morbidity in epilepsy, including its effects upon memory, attention, and general mental health. In short, there remains a strong imperative to understand seizure-induced transcriptional changes, and to develop methodologies for studying these.

To address this shortfall in our understanding, many groups have undertaken genetic, epigenetic, and transcriptome studies to explore causal links that lead to epilepsy and also explore possible protective changes (Chen et al., 2017; Hansen et al., 2014; Hunsberger et al., 2005; Pitkanen et al., 2015; Ryley Parrish et al., 2013). These large-scale transcriptomic data may further provide indications for novel therapeutic targets or extend our understanding of the basic pathophysiology of seizures by identifying “molecular hubs” (Dixit et al., 2016; Dong et al., 2020; Gorter et al., 2006; Kane et al., 2022). Despite this breadth of research, we still have a limited handle on many of the causal effects of brain injury, genetic alterations, status epilepticus, or other recognised triggers for epileptogenesis (Pitkanen et al., 2015).

Many groups have demonstrated large-scale transcriptomic changes during seizure activity (Dixit et al., 2016; Guelfi et al., 2019; Hawkins & Kearney, 2012; Li et al., 2022; O’Leary et al., 2020; Okamoto et al., 2010), indicating what might be termed the “molecular hallmark” of epileptiform activity. These studies have mainly been conducted using *in vivo* animal models or postmortem human brain tissue, primarily from hippocampal or whole-brain tissue sections (Conte et al., 2020; Jamali et al., 2006; Pitkanen et al., 2015; Zhang et al., 2018). While this is a valuable option for observing the changes in the genome, there are significant costs to these methods. For instance, the postmortem studies are potentially confounded by gene transcription changes during the terminal phase of life and may lack relevance for epileptogenesis in early-onset (development) epilepsy. The *in vivo* models, on the other hand, can mitigate this cost by using multiple epilepsy-induced animals, but these models often involve significant animal welfare impact, and the experimental control for such studies can be difficult.

To address both these shortfalls, we developed an *ex vivo* technique to probe for transcriptional changes in mouse neocortical tissue, comparing brain slices with pharmacologically induced seizures to control slices from the same brain, which were not exposed to the ictogenic treatment. We identified over 100 genes showing marked seizure-induced changes from the control levels, and analysed these using Gene ontology (GO), KEGG pathway enrichment, and Qiagen Ingenuity Pathway Analyses. We found no sex differences, but the set of genes whose expression patterns changed showed over-representation of several seizure-associated pathways, including MAPK, TNF, and neuroinflammatory signaling. These results prompted us to examine possible disease associations with our gene expression changes and using ingenuity pathway analysis (IPA) we found that the predicted disease of our differential expression (DE) was epilepsy.

Taken together, our results suggest that our *ex vivo* brain slices replicated the large-scale transcriptional changes observed following epileptiform activity *in vivo*, while offering levels of accessibility for recording and experimental control not usually possible *in vivo* and certainly not in clinical tissue. It further provides a highly refined approach to studying epileptic pathology, yielding multiple experimental replicates within biological individuals, while minimising welfare issues and harm to the animals involved.

## Methods

### Ethical Approval

All experimental procedures were approved by the Institutional Animal Care and Use Committee of the University of Alabama at Birmingham and the UK Home Office and Animals (Scientific Procedures) Act 1986 and approved by the Newcastle University Animal Welfare and Ethical Review Body (AWERB # 545)

### Slice preparation

Male and female C57BL/6 mice (Jackson Laboratory stock number 000664) (age ∼12 weeks) were used in this study. Mice were housed in individually ventilated cages in a 12-hour light, 12-hour dark regime. Animals received food and water *ad libitum*. Mice were sacrificed by cervical dislocation, brains removed and stored in cold cutting solution (in mM): 3 MgCl2; 126 NaCl; 26 NaHCO3; 3.5 KCl; 1.26 NaH2PO4; 10 glucose. For local field potential (LFP) recordings, 450µm horizontal sections were made containing the neocortex, entorhinal cortex, and the hippocampus using a Leica VT1000 vibratome (Nussloch, Germany). Slices were then transferred to an interface holding chamber and incubated for 1-2 hours at room temperature in artificial cerebrospinal fluid (aCSF) containing (in mM): 2 CaCl2; 1 MgCl2; 126 NaCl; 26 NaHCO3; 3.5 KCl; 1.26 NaH2PO4; 10 glucose.

### Extracellular field recordings

were performed using interface recording chambers. Slices were placed in the recording chamber perfused with modified aCSF to induce epileptiform activity (100µM 4-aminopyrimidine (4AP)). Recordings were obtained using normal aCSF-filled 1-3MΩ borosilicate glass microelectrodes (GC120TF-10; Harvard Apparatus, Cambridge, UK) placed in deep layers of the temporal association area. Experiments were performed at 33-36°C. The solutions were perfused at the rate of 3.5mls/min. Waveform signals were acquired using BMA-931 biopotential amplifier (Dataq Instruments, Akron, OH, USA), Micro 1401-3 ADC board (Cambridge Electronic Design, Cambridge, UK) and Spike2 software (v7.10, Cambridge Electronic Design). Signals were sampled at 10kHz, amplified (gain: 500), and bandpass filtered (1-3000Hz). The CED4001-16 Mains Pulser (Cambridge Electronic Design) was connected to the events input of CED micro 1401-3 ADC board and used to remove 50Hz hum offline. Seizure-like events (SLEs) were visually identified with their start time as the time of occurrence of high-frequency rhythmic bursts (tonic-phase) associated with high-frequency signals, and the events were considered to end when the interval between two after-discharges (clonic-phase) was ≥ 2s.

### Tissue collection and RNA extraction

After 3 hours of recordings, the neocortex (containing the auditory and somatosensory cortex) was subdissected away from the entorhinal cortex and the hippocampus, and then flash frozen with dry ice and stored at −80 degrees Celsius. For the RNA extraction, tissue was homogenized in Trizol using an Omni bead ruptor homogenizer (Omni International, Kennesaw, GA, USA), and RNA was isolated using the Trizol/choloform-isopropanol method following the manufacturer’s instructions (Invitrogen, Carlsbad, CA, USA). RNA concentration and purity were determined using a Thermo Scientific NanoDropOne (Fisher Scientific, Pittsburg, PA, USA). 14 total samples were collected, 8 of which were exposed to 4AP, and 6 controls that were treated identically except without 4AP treatment. Of the 4AP-treated slices, 5 were from male and 3 were from female mice. Likewise, 6 total slices from male and female brains were sequenced from the aCSF control group.

### RNA Sequencing and Differential Expression Analysis

Both the library prep and the sequencing was conducted by the Genomics Core Facility at Newcastle University on a Illumina NextSeq 500 (Illumina Inc.). Library prep was performed using the SmartSeq and Nextera XT kits. The sequencing was across two NextSeq 500 High-Output (150 cycle flow cells) producing 2 x 75 bp reads. All parts of the RNA-sequencing analysis pipeline were conducted in R and can be found in their entirety in the .html file Supplementary 1. Each of the samples were designated with their treatment followed by an assigned number and their sex (e.g. ACSF-5F) for the analysis. After sequencing, quality control was assessed using FastQC (www.bioinformatics.babraham.ac.uk/projects/fastqc/). The sequencing reads were aligned using the Rsubread package (Liao, Smyth, and Shi 2019) to the GRCm39 assembly of the mouse genome accessed through the Ensembl genome annotation platform. A count matrix was created from this alignment, which was used to conduct the differential expression (DE) analysis through the DESeq2 package (Love et al., 2014). Data was analyzed using a factorial design with the factors 4AP treatment, sex, and a sex:treatment interacting factor to determine the cause of particular gene expression changes. Genes with a calculated adjusted p-value of < .05 and absolute log_2_ Fold Change (FC) values greater than .585 (a 50% change in expression) were considered significant. Once our analysis of sex-specific differences in response to treatment revealed no significant DE genes, we focused on the impact of the 4AP treatment on DE. Hierarchical clustering was performed for a comparison of the top 30 upregulated and 10 downregulated genes based on log_2_FC and adjusted p-value. The respective count values from the DE analysis were converted into z-scores and clustered and converted into a heatmap using the ComplexHeatmap package (Gu et al., 2016).

### GO and other Functional Enrichment Analyses

To further characterize the associations between the 4AP treatment DE genes, functional analysis was performed on the list of 110 DE genes, referred to hereafter as the gene set. Enrichment of Gene Ontology (GO) Biological Processes (BP) and Kyoto Encyclopedia of Genes and Genomes (KEGG) pathways analyses were assessed using the clusterProfiler package (Yu et al., 2012). For the GO:BP analysis, the simplify function was used to filter redundant parent-child associations into singular, informative BP terms. Full GO results can be found in Supplementary Data 3. The meshes package was used to access MeSH Disease annotations for the gene list to investigate the diseases that our gene set was associated with (Yu, 2018). Twenty-four brain-related diseases and disorders were selected to highlight, and the full list of 247 enriched disease categories, along with the full GO:BP and KEGG results, can be found in Supplementary Data 4. Unless otherwise specified, all plots are of the top enriched categories (adjusted p-values < 0.05) and were plotted using the enrichplot package (Yu et al., 2012). To analyze transcription factor binding sites, the Homer software package was used to do an overrepresentation analysis for our gene list, and both *de novo* and known motifs were selected (Heinz et al., 2010). The function was set to search 1000 bp upstream and 100 bp downstream of the transcription start of the specific gene. For a background dataset, we selected the whole genome to enable statistical analysis.

### IPA analysis

Quiagen Inuititve Pathway analysis was carried out on our DE gene set using the log_2_FC and adjusted p-value for each gene (Kramer et al., 2014). All DE genes mapped to the knowledge base and the z-score setting was used to quantify specific pathway activation status. Canonical pathway analysis was examined and extracted to identify pathways that were predicted by the algorithm to be activated or inhibited, the full list of which can be found in Supplementary File 4. Pathways with greater than log B-H adjusted p-values of 1.3 were considered significant. Likewise, IPA Diseases and Function analysis was used to predict disease states based on our DE genes. B-H adjusted p-values were used to test for significance.

### Statistics

Electrophysiology data were analyzed offline using Matlab R2018b (MathWorks, USA). Statistical tests shown in figure 1 were analyzed using a Student’s t-tests. R was used for the analysis of all the RNA sequencing data as described above.

**Figure 1.**
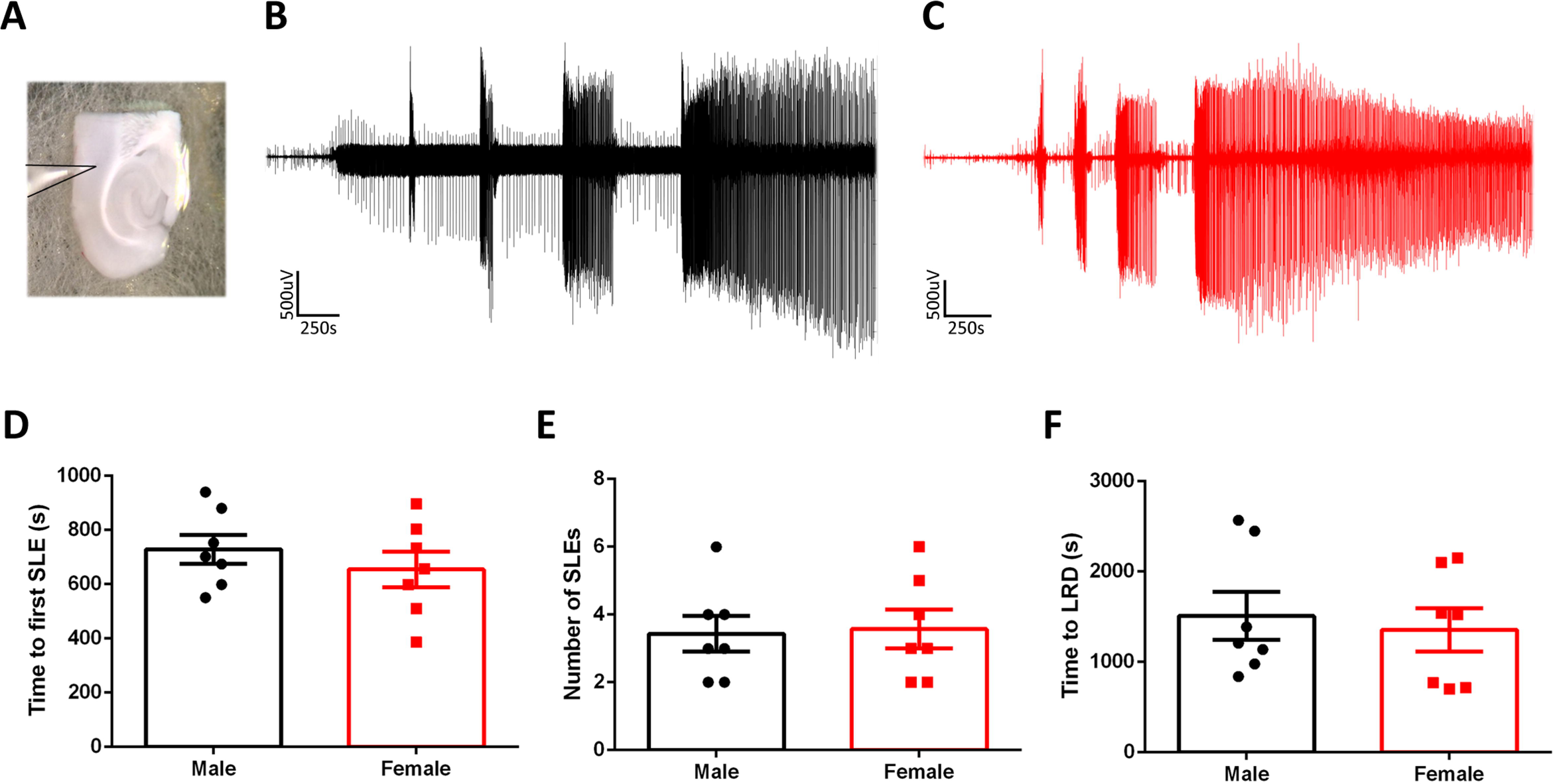
Induced seizure-like activity is similar from slices made from male and female mice. A) Example of typical deep-layer LFP recording. B) Example LFP trace from a slice made from a male mouse and exposed to 4AP. C) Equivalent recording from a female mouse. D) There is no difference in the time to the first seizure-like event (SLE) between slices made from male and female mice (unpaired t-test, p = 0.24). D) There is no difference in the number of SLE between slices made from male and female mice (unpaired t-test, p = 0.72). E) There is no difference in the time to late recurrent discharges between male and female mice (unpaired t-test, p = 0.68).

## Results

### *Ex vivo* epileptiform activity evolves in a similar pattern in both males and females

We sought to determine the utility of *ex vivo* seizure models for studying seizure-induced transcriptomic changes. Brain slices were prepared from male and female mice, and bathed in 100 µM 4AP while recording the evolving epileptiform activity patterns. 4-AP induced a characteristic build-up of seizure-like activity over time, starting with early interictal-like discharges, followed by intermittent seizure-like events (SLEs), and culminating in late recurrent discharges (Codadu et al., 2019). This final stage appears to represent a steady state of hyperexcitability, and has been suggested to be equivalent to *in vivo* status epilepticus (Figure 1B, C) (Dreier et al., 1998; Zhang et al., 1995).

### *Ex vivo* epileptiform activity shows minimal gene expression differences between the sexes

We found no difference in the evolving pattern of epileptiform activity between male and female mice (Figure 1D, E, F), suggesting that naïve susceptibility to pharmacologically-induced seizure activity is similar between the sexes. We then examined whether the equivalent epileptiform activity induced distinct transcriptomic expression patterns between sexes. To address this, we conducted RNA-sequencing on 14 different brain slices: 8 treated with 4AP (5 male and 3 female), all of which developed intense epileptiform discharges, and 6 control slices (3 male and 3 female), where no drug was added, and which did not display any pathological discharges. The duration of the treatment was 3 hours. After this period, the RNA was isolated from the tissue. Illumina sequencing was performed on these samples, and the reads were aligned to the mouse genome for subsequent analysis of differential expression (DE) using the DESeq2 package.

To determine the specific factors contributing to the gene expression changes, namely sex, seizure induction, or their combination, we employed the design feature of DESeq2 to account for these variables by segmenting the variance in the samples. Initially, we examined any sex-specific gene expression changes associated with epileptiform activity. Principal component analysis (PCA) revealed that both male and female samples exhibited the greatest variance between slices with induced seizure-like activity versus control samples, rather than being influenced by sex alone (Figure 2A). Consistent with this result, only 15 genes exhibited differential expression based on sex of the sample, with the most significant difference between the sexes being found in genes on the X or Y chromosomes (Figure 2B, Table 1). Further analysis of the combinatorial DE between the responses of each sex to seizures revealed no significant gene changes (Figure 2C). Consequently, we concluded that induced seizure-like activity elicits minimal differences between male and female mice when utilizing an *ex vivo* preparation.

**Figure 2.**
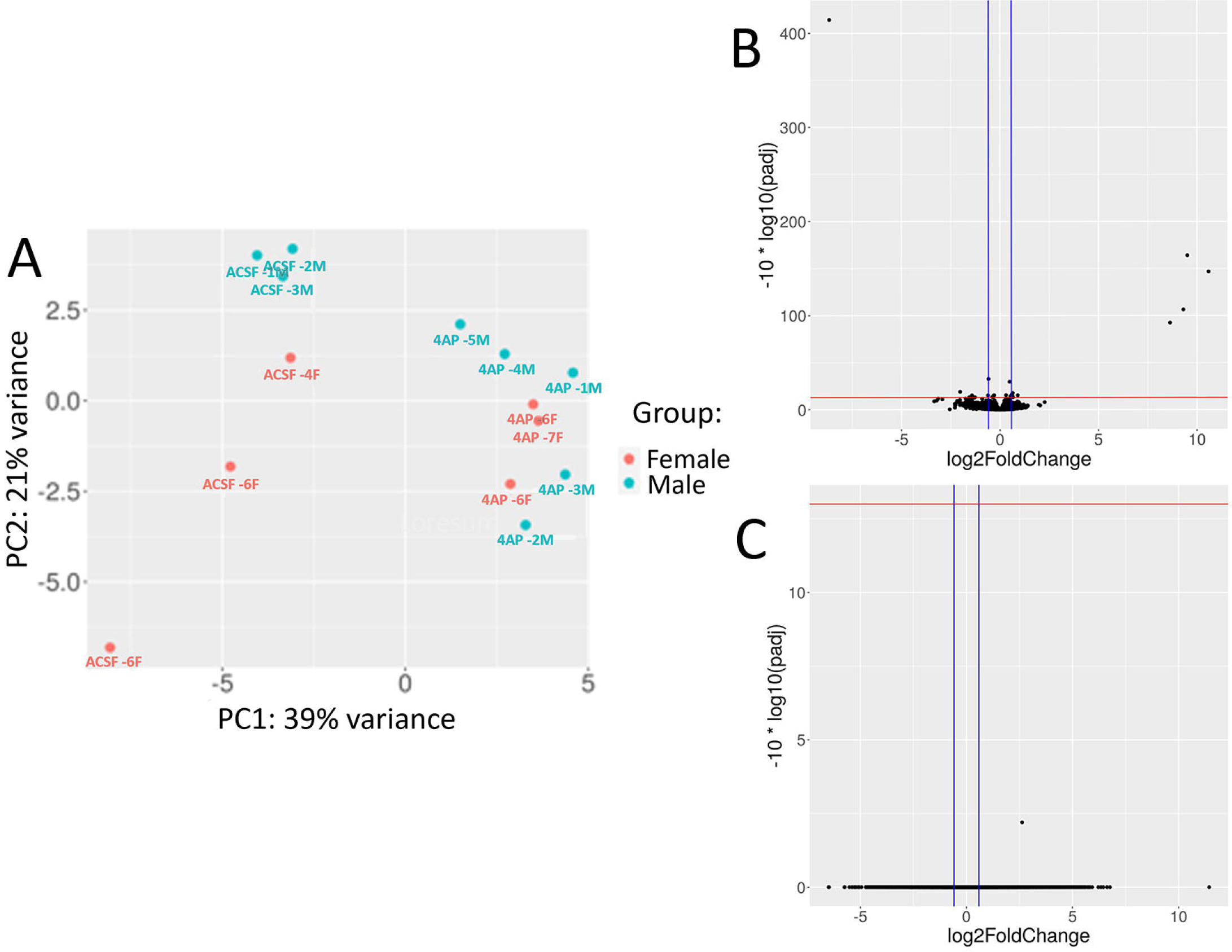
RNA-seq of male and female slices show no differences between sex in gene expression in response to epileptiform activity. A) PCA plot of the RNA-seq gene expression data from each of the 14 samples separated by sex as shown by the female samples in orange and the males in blue. B) Volcano plot showing DE of genes between male and female samples controlling for treatment condition. Upregulated genes are higher expressed in the male slices and down-regulated genes are present more in the female slices. Genes are plotted with their log2FC against the -log10(padj). The red horizontal line is at 13 which denotes a FDR=0.05 and the blue vertical lines are at −0.58 and 0.58 showing the log2FC cutoffs.

**Table 1.** List of differential expressed genes based on sex.

| Gene Name | Log2FC | padj | Chromosome | Description |
| --- | --- | --- | --- | --- |
| Eif2s3y | 10.59003969 | 2.12E-15 | Y | eukaryotic translation initiation factor 2, subunit 3, structural gene |
| Ddx3y | 9.512385256 | 3.98E-17 | Y | DEAD box helicase 3 |
| Kdm5d | 9.308209984 | 2.30E-11 | Y | lysine (K)-specific demethylase 5D |
| Xist | -8.653909241 | 3.78E-42 | X | inactive X specific transcripts |
| Uty | 8.635518223 | 5.76E-10 | Y | ubiquitously transcribed tetratricopeptide repeat containing |
| Depp1 | -2.022647721 | 0.0126 | 6 | DEPP1 autophagy regulator |
| Tubb4b-ps2 | -1.420474479 | 0.0298 | 1 | tubulin, beta 4B class IVB, pseudogene 2 |
| Etnppl | 0.904044518 | 0.0292 | 3 | ethanolamine phosphate phospholyase |
| Lrrc55 | 0.657426506 | 0.0473 | 2 | leucine rich repeat containing 55 |
| Cd34 | 0.641665353 | 0.0167 | 1 | CD34 antigen |
| Rskr | 0.629185198 | 0.0379 | 11 | ribosomal protein S6 kinase related |
| Rassf3 | 0.620279552 | 0.0281 | 10 | Ras association (RalGDS/AF-6) domain family member 3 |
| Crhbp | -0.60703997 | 0.03 | 13 | corticotropin releasing hormone binding protein |
| Eng | 0.587780636 | 0.0281 | 2 | endoglin |
| Eif2s3x | -0.580478737 | 0.0005 | X | eukaryotic translation initiation factor 2, subunit 3, structural gene |

### DE analysis of *ex vivo* epileptiform activity reveals significant gene expression changes

After finding no significant DE genes in our sex-specific analysis, we evaluated differences between the experimental groups, using both principal component analysis (PCA) and hierarchical clustering (Figure 3A, B). The volcano plot in Figure 3C visually represents the DE genes, showing upregulation of 82 genes and downregulation of 28 genes in response to seizure induction (refer to Supplementary file 2 for the full list). As expected, upon initial analysis of the DE genes, we observed that several of the most highly upregulated transcripts were immediate early genes, including c-Fos, early growth response protein 4 (Egr4), Activity-regulated cytoskeleton-associated protein (Arc), ADP-ribosylation factor-like protein 4D (Arl4d), and Neuronal PAS domain protein 4 (Npas4). Notably, Npas4 has also been described as a transcription factor with a potential antiseizure role (Wang et al., 2014). Among the downregulated genes, we found examples that might be predicted to either decrease neuronal excitability (e.g. the calcium voltage channel Cav3.2, CACNA1H; log_2_FC of −0.70) or increase it (e.g. Adora2a, which codes for the adenosine A2A receptor; log_2_FC of −2.61).

**Figure 3.**
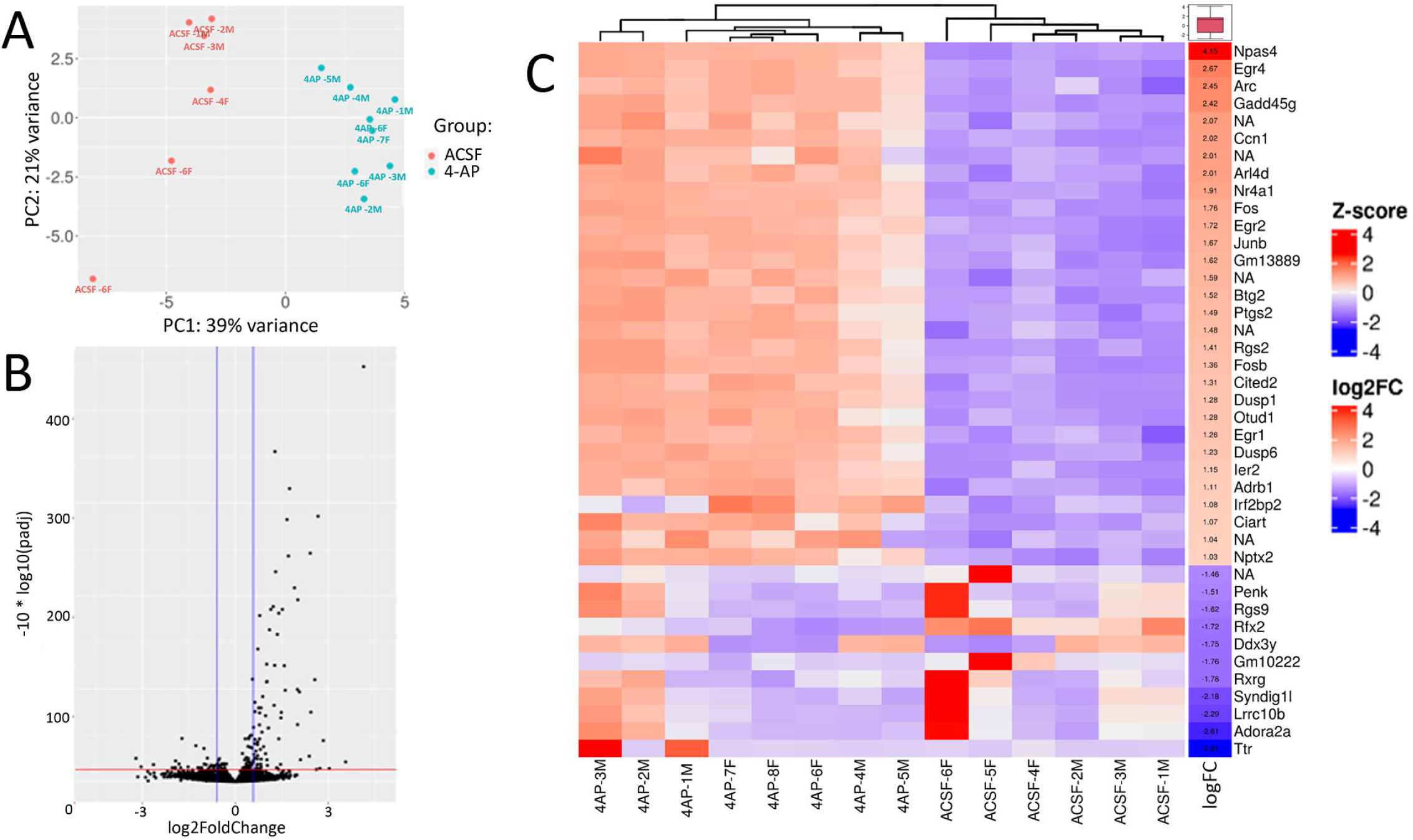
DE analysis of *ex vivo* slices show large scale gene expression changes in response to 4AP treatment. A) PCA plot of the samples as in Figure 2A, separated by aCSF and 4AP treatment shown in orange and blue respectively. B) Volcano plot as in 2B of the DE genes after seizure induction. C) Heat map and hierarchical clustering of the top 30 upregulated and 10 downregulated genes for each of the samples shown in a column. A gradient from red to blue represents the expression of the gene in each sample, the red box denotes a higher expression and a blue box denotes lower expression. The columns show the log2FC value and average expression values for the specific genes.

In order to explore these gene expression changes more deeply, we used the Gene Ontology (GO) and KEGG pathway resources (Ashburner et al., 2000; Gene Ontology et al., 2023; Kanehisa & Goto, 2000). GO:Biological Process analysis revealed a significant enrichment of over 300 categories (refer to Supplementary 3 for the full list). To address redundancy resulting from hierarchical ontology parent-child relationships, we utilized the clusterProfiler package to streamline the GO results, resulting in 159 simplified categories. Figure 4A presents a selection of 20 enriched categories, illustrating various cellular responses to seizure activity. Among the enriched categories, several signaling pathways stood out, including the p38 MAPK, ERK 1/2 cascade, response to TGFB signaling, neurotropin signaling, calcium ion response, and response to peptide hormone pathways. While the involvement of the MAPK pathway and calcium ion response in epileptogenesis have long been recognized, the roles of these other pathways in relation to seizure disorders are not yet fully established (Gautam et al., 2021; Steinlein, 2014). Notably, several categories related to neuron death, apoptosis, and transcription activation in response to stress were also enriched, which may be attributed to the extensive brain damage caused by seizure activity. We found several intriguing pathways, including those involving the positive regulation of miRNAs, catecholamine secretion, and categories associated with vasculogenesis, angiogenesis, as well as learning and memory. Additionally, when we performed KEGG pathway enrichment analysis, we observed overlaps within the enriched pathways identified through GO analysis, particularly in the MAPK signaling pathway, TNF signaling pathway, and apoptosis (Figure 4B). These findings provide valuable insights into the complex molecular processes underlying seizure-induced changes.

**Figure 4.**
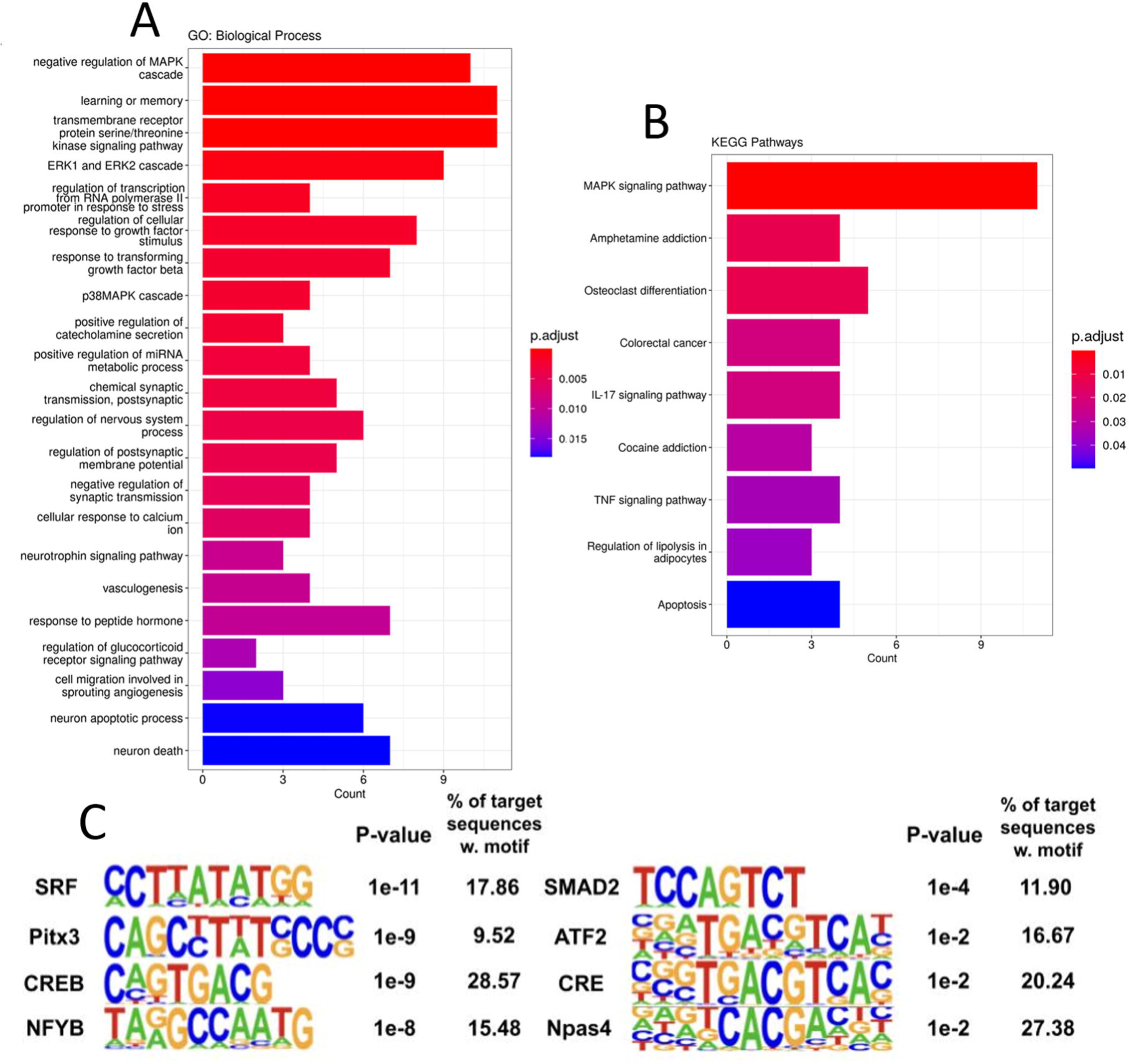
Gene ontology and pathway analysis reveal complex molecular regulatory networks post epileptiform activity. A) Bar plot of 20 significantly enriched GO: Biological Process categories, bar color shows the relative significance based on the adjusted p-value and the length of the bar shows the number of upregulated genes enriched in the respective categories. B) KEGG pathway enrichment analysis was performed on the gene set. Bar plot is arranged as in 4A and includes the 9 pathways that were significantly enriched. C) Homer motif software results from enrichment analysis of the DE genes, motifs were either created *de novo* or were from the known database. P-values were calculated against the full list of background genes in the software and % of genes with a respective transcription factor motif are shown.

In light of the broad activity observed across various signaling pathways, we aimed to identify critical transcription factors responsible for the observed gene expression changes. To accomplish this, we analyzed our DE genes for enrichment of specific transcription factor binding motifs using the HOMER software. Of particular interest was the cAMP response-element binding protein (CREB), known to be activated by phosphorylation downstream of the MAPK and PKA pathways, as well as the crucial role of CREB-mediated transcriptional regulation in response to epileptogenesis (Mertz et al., 2020; Sun et al., 2022; Wang et al., 2020). Further analysis using HOMER showed a CREB motif upstream of 29% of the genes (Figure 4C), indicative of the involvement of CREB in mediating a large proportion of our observed gene expression changes. Additionally, we identified highly enriched motifs for SRF, ATF2, and Npas4, which were upstream of over 15% of the DE genes. These transcription factors play significant roles in cellular stress response, and many have been implicated in mediating gene expression changes following seizures (Losing et al., 2017; Mielke et al., 1999; Shan et al., 2018). By identifying these enriched transcription factor binding motifs, we provide insights into the potential regulatory mechanisms driving the observed gene expression patterns. Further investigation into the specific interactions between these transcription factors and their target genes could shed light on the molecular processes underlying the cellular response to epileptiform activity.

To deepen our understanding of the specific cellular pathway responses to our observed gene expression changes, we leveraged Qiagen IPA analysis, which provides a machine learning algorithm and gene annotations to identify pathway enrichment and to predict pathway activation state (Kramer et al., 2014). Our results, shown in Figure 5A, illustrate the complex signalling environment triggered by epileptiform activity. Several of the seizure-induced upregulated pathways that we identified are related to inflammatory and immune signalling, including IL-6 and IL-8 cytokine signalling, HMGB1 production, neuroinflammation, and CXCR4 signalling. CXCR4, the receptor for the CXCL12 ligand, has been implicated in neuroinflammation and has an emerging role in the induction of pro-seizure activity (Zhou et al., 2017). Furthermore, our GO and KEGG analysis also showed an enrichment for the production of IL-1β and IL-17 signalling, providing increased evidence of the importance of immune signalling after seizure activity (Dong et al., 2020; Wolinski et al., 2022). IPA predicted a downregulation of cAMP signalling, which would lead to a decrease in MAPK and PKA pathways (Bozzi et al., 2011). Upon further investigation, we observed an upregulation of several phosphatases involved in dampening MAPK signalling (DUSP1/4/5/6/14), many of which have been implicated in seizure responses (An et al., 2021). This might be considered to be a compensatory mechanism opposing the dramatic activation of these signalling pathways following seizure induction, and could be potential therapeutic targets (Kirchner et al., 2020).

**Figure 5.**
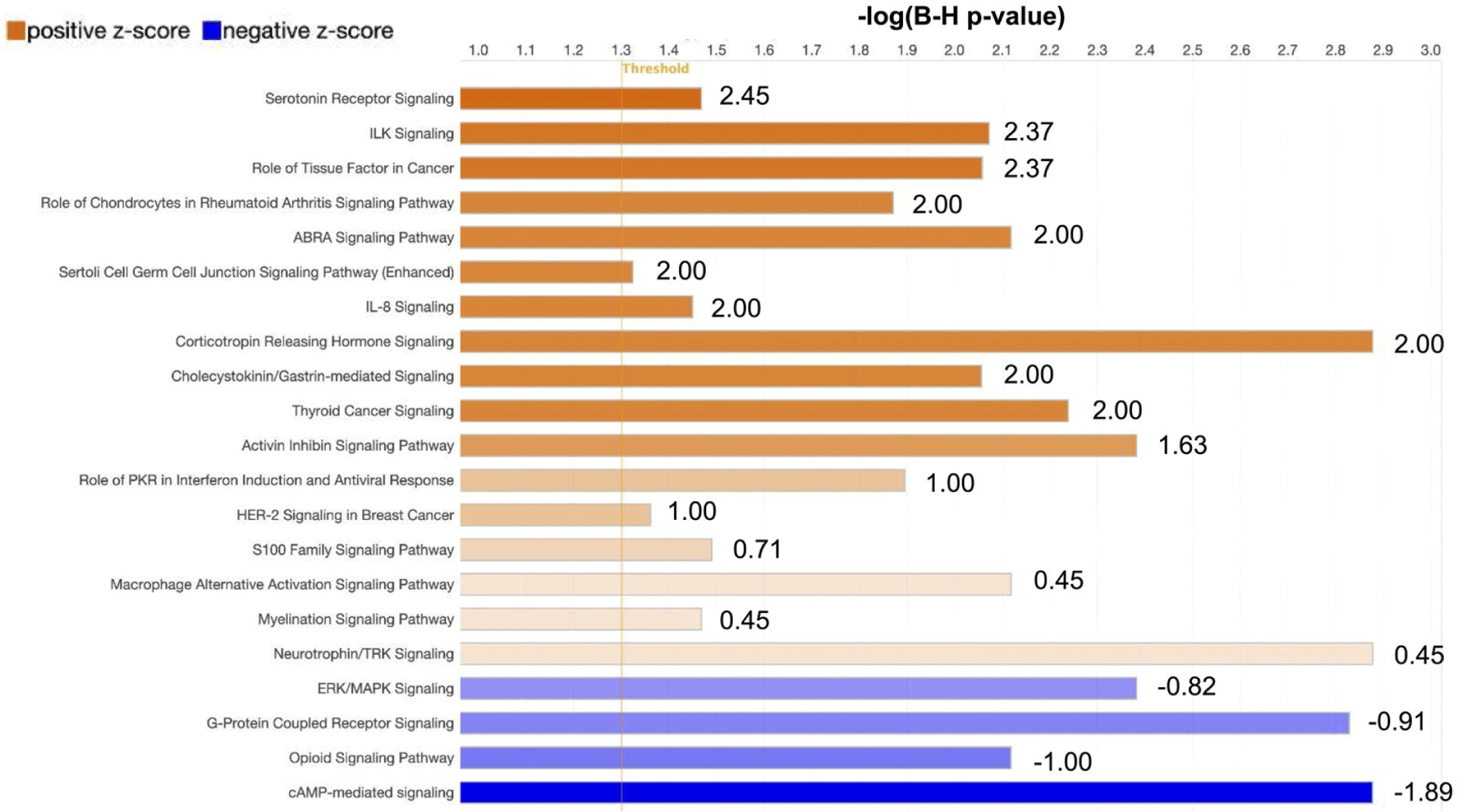
IPA predicts the activation of inflammatory regulatory networks and inhibition of cAMP signalling in response to epileptiform activity. Plot shows IPA canonical pathway prediction status based on the DE genes. Respective pathways are shown with the length of the bar representing the -log of the B-H adjusted p-value for multiple testing. The bar color represents the z-score value activation (orange) or inhibition (blue) predicted state of the pathway.

### Ingenuity Pathway Analysis predicted epilepsy as the causal disease

We were interested to compare the transcriptional changes following acute induction of seizure-like activity *ex vivo*, with the gene expression profiles in chronic epilepsy. To examine this, we utilized MeSH (Medical Subject Headings) analysis of epilepsy-associated genes in published articles on PubMed (http://pubmed.ncbi.nlm.nih.gov). This analysis links terms, such as a particular gene, to specified categories, such as neurological conditions (Yu, 2018). Using MeSH, we performed an enrichment analysis with our DE gene set for diseases associated with them. We identified 247 diseases that could potentially be associated with the observed gene expression changes, focusing particularly on neurological disorders and brain injuries (Figure 6A, complete list available in Supplemental 5). We found significant enrichment of status epilepticus, absence epilepsy, and tonic-clonic epilepsy. Additionally, we observed categories related to substance addiction (e.g., cocaine-related disorders, morphine dependence), brain development disorders, and brain injuries. To complement the enrichment analysis, we employed the predictive capabilities of IPA to determine the diseases most likely responsible for the gene expression changes based on the gene set. Epilepsy was the top disease state predicted based on the interaction of the 30 genes shown in Figure 6B. Taken together, these results suggest that our *ex vivo* approach can replicate on a small scale the gene expression phenotype of seizure activity.

**Figure 6.**
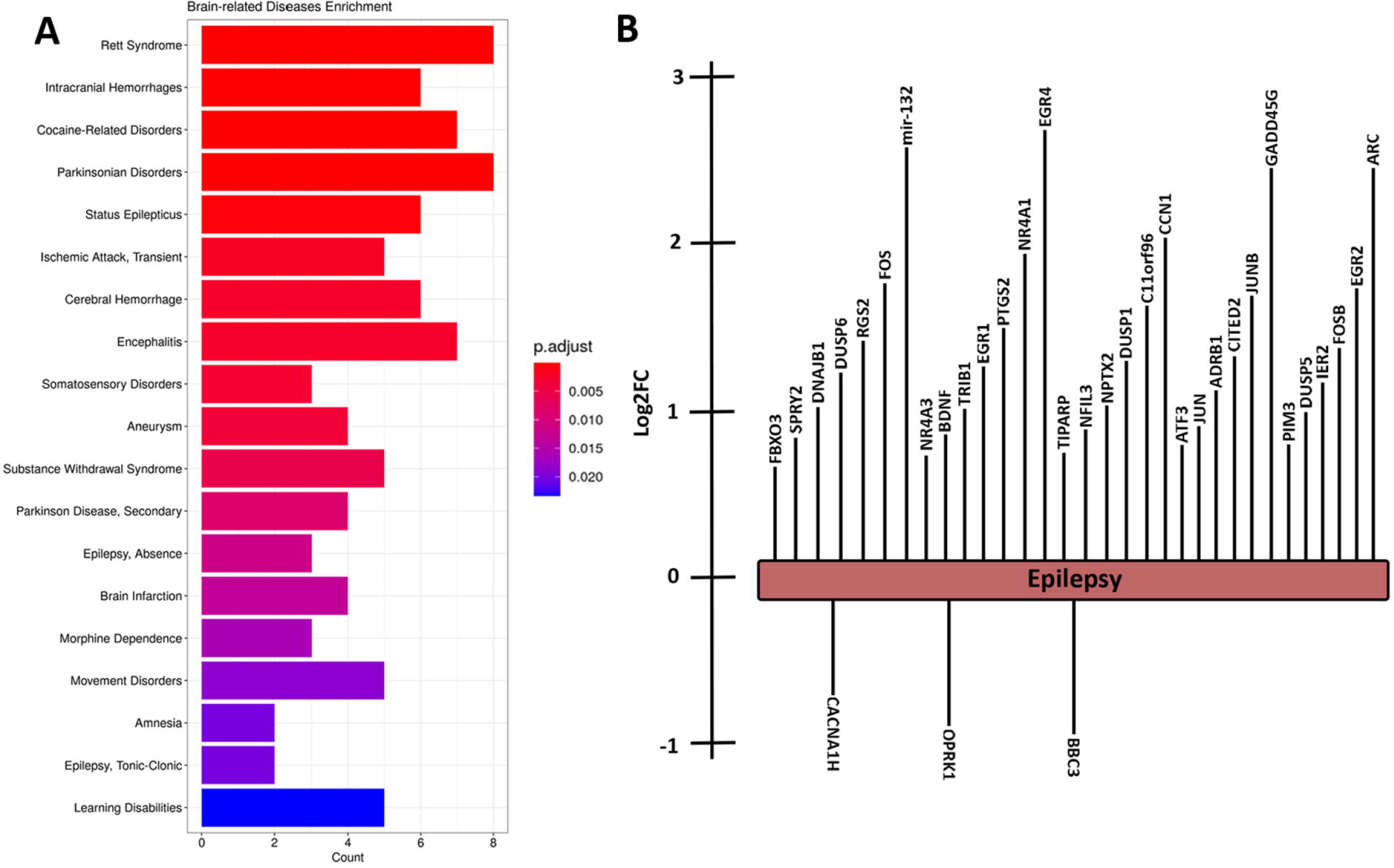
Disease Enrichment and IPA analysis of upregulated genes show significant links to epilepsy and other brain injuries. A) Enrichment analysis of MeSH disease annotations from the upregulated gene set. 247 disease terms were enriched and the bar plot is a selection of 20 enriched neuro-related diseases. Bar plot is as Figure 4. B) IPA analysis of gene set reveals epilepsy as the top disease state predicted from the DE genes with a p-value of 1×10^(−28). Image shows the specific genes associated with the disease in the IPA knowledge base on enriched in the analysis. Length of the bar represents the log2FC values for each gene.

## Discussion

*Ex vivo* studies of large network activity patterns are sometimes disregarded because they are considered unrepresentative of the *in vivo* condition. Yet such simplified models offer many experimental advantages, and so it is helpful to examine how they might be used to tease apart the nature of especially complex neurological phenomena such as epileptic pathophysiology. Here, we performed transcriptomic analyses of brain slices that have experienced periods of intense epileptiform activation, finding changes in many genes that have been highlighted in other studies of epilepsy, as well as predicting key upstream transcription factors and potential therapeutic targets. Our *ex vivo* data were further validated by finding that the most closely related disorder associated with the gene expression patterns is indeed epilepsy.

These simple *ex vivo* preparations offer various benefits. Firstly, the ease of access to the tissue allows for a far tighter correlation between the observed gene expression profiles and specific patterns of electrophysiology activity, mapped with precision using multielectrode arrays (Mahadevan et al., 2022; Thouta et al., 2022), patch clamp recordings, or imaging (Calin et al., 2021; Cammarota et al., 2013; Codadu et al., 2019). Experimental manipulations using pharmacology, optogenetic, chemogenetic, or electrical stimulation can be similarly targeted (Calin et al., 2018; Graham et al., 2023; Papasavvas et al., 2020). Additionally, one animal can provide many experimental replicates, reducing variability and cost of the study, while also increasing the power of the study, thereby increasing the likelihood of identifying potential therapeutic targets. *Ex vivo* preparations do have their limitations; our preparations were fixed just a few hours after the slices are prepared, thus providing information only on the earliest transcriptional changes. Slower processes, though, may yet be amenable to experimental study in organotypic cultures (Lillis et al., 2015). In general, though, our validation of the patterns of translational changes in these acute preparations, in conjunction with the advantages they offer, particularly with respect to animal welfare considerations (the “3Rs”, reduction, refinement, replacement) (Lidster et al., 2016), provide a strong incentive for their inclusion in the arsenal of experimental approaches for studying epilepsy.

Notably, we found that slices from male and female mice show similar evolution of seizure-like activity following application of the convulsant drug 4AP, as well as similar patterns of seizure-induced gene expression changes. The absence of obvious sex differences in acutely induced seizure-like activity, thus appears to be subtly different from situation in chronic epilepsy. For instance, seizure severity has previously been reported to be higher in male than in female rats following pilocarpine administration (Matovu & Cavalheiro, 2022). In clinical medicine, men also show higher epilepsy prevalence while also having more severe seizures (Christensen et al., 2007; Christian et al., 2020; McHugh & Delanty, 2008), although a factor here may be that men are more likely to sustain head injuries. Notably, though, sex differences were found in gene expression profiles of blood samples from patients with epilepsy (Zhu et al., 2021). This study incorporated a wide demographic range and multiple seizure types. Further work will be required to ascertain whether gender differences are restricted to specific types of epilepsy or experimental models or are specific to the tissue sampled (blood versus the cortical tissue itself).

Since our own data found no difference between sexes, we pooled the male and female datasets for subsequent analyses to increase their statistical power, finding 82 genes to be upregulated and 28 to be downregulated by the prior seizure-like activity. Many of these were immediate early genes, with the upregulation of Npas4 being particularly interesting given its suggested anti-seizure role (Wang et al., 2014). Similarly, one may impute homeostatic, anti-epileptic, negative feedback mechanisms (Cain et al., 2018; Chen et al., 2014; Trevelyan et al., 2023; Wang et al., 2014) from the known function of certain genes (e.g. upregulation of Npas4, or downregulation of CACNA1H). In contrast, other transcriptional changes we observed appear to enact positive feedback by promoting more seizure activity, such as the downregulation of Adora2a. Adenosine acts as an endogenous anticonvulsive, and the A2A receptor is one of its primary targets (Baltos et al., 2023). These results may be reconciled by the observation that long-term activation of the A2A receptor has been demonstrated to result in internalization (Palmer et al., 1994; Sheth et al., 2014), in which case downregulation of Adora2a may also be anti-epileptic by the same mechanism.

We also examined gene changes within the context of their involvement in various signalling and cellular pathways. Notable among these were changes in transcription factors and genes involving the MAPK pathway, neurotrophins, and calcium ion response. Previous work also implicated the MAPK pathway as a potential disease-modifying route in epilepsy (Parrish et al., 2018), while transcription factors, such as CREB, are increasingly considered as targets for therapy in seizure disorders (Sun et al., 2022).

A major goal for the treatment of complex conditions such as epilepsy is to find how to use big transcriptional data sets for precision medicine for individual patients, but this still lies in the future (Knowles et al., 2022; Orsini et al., 2018; Striano & Minassian, 2020). We present a case for progressing this field using simplified epilepsy models, such as those used here, paired with other new toolkits in neuroscience, such as optogenetics and chemogenetics, to advance our understanding of these conditions and create avenues for novel treatment options.

## Author Contributions

The project was conceived by R. Ryley Parrish, Rita Cowell, and Andrew J. Trevelyan. Experiments were performed by R. Ryley Parrish, Neela K. Codadu, and Laura J. McMeekin. Analysis was performed by Alec J. Vaughan, R. Ryley Parrish, Jonathon Hill, Simon Cockell, Kutter Hine, Isaac Stubbs, and Neela K. Codadu. Figures were created by Isaac Stubbs, Alec Vaughan, R. Ryley Parrish and Neela K. Codadu. Manuscript was written and edited by Alec J. Vaughan, R. Ryley Parrish, Andrew J. Trevelyan, Isaac Stubbs, Jonathan Hill, Kutter Hine. All authors approved the final version of this manuscript.

## Acknowledgments

We would like to thank Rachel Queen for her early assistance with the RNA sequencing analysis. We would also like to thank the Newcastle University sequencing core for their help with preforming the RNA sequencing. In addition, we thank Jing Wang of the UAB Neuroscience Core Center (Project number: 5P30NS047466-13).

## Availability of data

The data and code that support the findings of this study are available from the corresponding author upon request. The RNA-sequencing datasets generated and analysed during the current study will also be made public on the NCBI website following peer-reviewed publication.

## Acknowledgments

The authors declare no conflicts of interests.

## References

An, N., Bassil, K., Al Jowf, G. I., Steinbusch, H. W. M., Rothermel, M., de Nijs, L., & Rutten, B. P. F. (2021). Dual-specificity phosphatases in mental and neurological disorders. Prog Neurobiol, 198, 101906. 10.1016/j.pneurobio.2020.101906

Ashburner, M., Ball, C. A., Blake, J. A., Botstein, D., Butler, H., Cherry, J. M., Davis, A. P., Dolinski, K., Dwight, S. S., Eppig, J. T., Harris, M. A., Hill, D. P., Issel-Tarver, L., Kasarskis, A., Lewis, S., Matese, J. C., Richardson, J. E., Ringwald, M., Rubin, G. M., & Sherlock, G. (2000). Gene ontology: tool for the unification of biology. The Gene Ontology Consortium. Nat Genet, 25(1), 25–29. 10.1038/75556

Baltos, J. A., Casillas-Espinosa, P. M., Rollo, B., Gregory, K. J., White, P. J., Christopoulos, A., Kwan, P., O’Brien, T. J., & May, L. T. (2023). The role of the adenosine system in epilepsy and its comorbidities. Br J Pharmacol. 10.1111/bph.16094

Bozzi, Y., Dunleavy, M., & Henshall, D. C. (2011). Cell signaling underlying epileptic behavior. Front Behav Neurosci, 5, 45. 10.3389/fnbeh.2011.00045

Cain, S. M., Tyson, J. R., Choi, H. B., Ko, R., Lin, P. J. C., LeDue, J. M., Powell, K. L., Bernier, L. P., Rungta, R. L., Yang, Y., Cullis, P. R., O’Brien, T. J., MacVicar, B. A., & Snutch, T. P. (2018). Ca(V) 3.2 drives sustained burst-firing, which is critical for absence seizure propagation in reticular thalamic neurons. Epilepsia, 59(4), 778–791. 10.1111/epi.14018

Calin, A., Ilie, A. S., & Akerman, C. J. (2021). Disrupting Epileptiform Activity by Preventing Parvalbumin Interneuron Depolarization Block. J Neurosci, 41(45), 9452–9465. 10.1523/JNEUROSCI.1002-20.2021

Calin, A., Stancu, M., Zagrean, A. M., Jefferys, J. G. R., Ilie, A. S., & Akerman, C. J. (2018). Chemogenetic Recruitment of Specific Interneurons Suppresses Seizure Activity. Front Cell Neurosci, 12, 293. 10.3389/fncel.2018.00293

Cammarota, M., Losi, G., Chiavegato, A., Zonta, M., & Carmignoto, G. (2013). Fast spiking interneuron control of seizure propagation in a cortical slice model of focal epilepsy. J Physiol, 591(4), 807–822. 10.1113/jphysiol.2012.238154

Chen, T., Giri, M., Xia, Z., Subedi, Y. N., & Li, Y. (2017). Genetic and epigenetic mechanisms of epilepsy: a review. Neuropsychiatr Dis Treat, 13, 1841–1859. 10.2147/NDT.S142032

Chen, Y., Parker, W. D., & Wang, K. (2014). The role of T-type calcium channel genes in absence seizures. Front Neurol, 5, 45. 10.3389/fneur.2014.00045

Christensen, J., Vestergaard, M., Pedersen, M. G., Pedersen, C. B., Olsen, J., & Sidenius, P. (2007). Incidence and prevalence of epilepsy in Denmark. Epilepsy Res, 76(1), 60–65. 10.1016/j.eplepsyres.2007.06.012

Christian, C. A., Reddy, D. S., Maguire, J., & Forcelli, P. A. (2020). Sex Differences in the Epilepsies and Associated Comorbidities: Implications for Use and Development of Pharmacotherapies. Pharmacol Rev, 72(4), 767–800. 10.1124/pr.119.017392

Codadu, N. K., Graham, R. T., Burman, R. J., Jackson-Taylor, R. T., Raimondo, J. V., Trevelyan, A. J., & Parrish, R. R. (2019). Divergent paths to seizure-like events. Physiol Rep, 7(19), e14226. 10.14814/phy2.14226

Conte, G., Parras, A., Alves, M., Olla, I., De Diego-Garcia, L., Beamer, E., Alalqam, R., Ocampo, A., Mendez, R., Henshall, D. C., Lucas, J. J., & Engel, T. (2020). High concordance between hippocampal transcriptome of the mouse intra-amygdala kainic acid model and human temporal lobe epilepsy. Epilepsia, 61(12), 2795–2810. 10.1111/epi.16714

Dixit, A. B., Banerjee, J., Srivastava, A., Tripathi, M., Sarkar, C., Kakkar, A., Jain, M., & Chandra, P. S. (2016). RNA-seq analysis of hippocampal tissues reveals novel candidate genes for drug refractory epilepsy in patients with MTLE-HS. Genomics, 107(5), 178–188. 10.1016/j.ygeno.2016.04.001

Dong, X., Hao, X., Xu, P., Fan, M., Wang, X., Huang, X., Jiang, P., Zeng, L., & Xie, Y. (2020). RNA sequencing analysis of cortex and hippocampus in a kainic acid rat model of temporal lobe epilepsy to identify mechanisms and therapeutic targets related to inflammation, immunity and cognition. Int Immunopharmacol, 87, 106825. 10.1016/j.intimp.2020.106825

Dreier, J. P., Zhang, C. L., & Heinemann, U. (1998). Phenytoin, phenobarbital, and midazolam fail to stop status epilepticus-like activity induced by low magnesium in rat entorhinal slices, but can prevent its development. Acta Neurol Scand, 98(3), 154–160. 10.1111/j.1600-0404.1998.tb07286.x

Gautam, V., Rawat, K., Sandhu, A., Kumari, P., Singh, N., & Saha, L. (2021). An insight into crosstalk among multiple signaling pathways contributing to epileptogenesis. Eur J Pharmacol, 910, 174469. 10.1016/j.ejphar.2021.174469

Gene Ontology, C., Aleksander, S. A., Balhoff, J., Carbon, S., Cherry, J. M., Drabkin, H. J., Ebert, D., Feuermann, M., Gaudet, P., Harris, N. L., Hill, D. P., Lee, R., Mi, H., Moxon, S., Mungall, C. J., Muruganugan, A., Mushayahama, T., Sternberg, P. W., Thomas, P. D., … Westerfield, M. (2023). The Gene Ontology knowledgebase in 2023. Genetics, 224(1). 10.1093/genetics/iyad031

Gorter, J. A., van Vliet, E. A., Aronica, E., Breit, T., Rauwerda, H., Lopes da Silva, F. H., & Wadman, W. J. (2006). Potential new antiepileptogenic targets indicated by microarray analysis in a rat model for temporal lobe epilepsy. J Neurosci, 26(43), 11083–11110. 10.1523/JNEUROSCI.2766-06.2006

Graham, R. T., Parrish, R. R., Alberio, L., Johnson, E. L., Owens, L., & Trevelyan, A. J. (2023). Optogenetic stimulation reveals a latent tipping point in cortical networks during ictogenesis. Brain, 146(7), 2814–2827. 10.1093/brain/awac487

Gu, Z., Eils, R., & Schlesner, M. (2016). Complex heatmaps reveal patterns and correlations in multidimensional genomic data. Bioinformatics, 32(18), 2847–2849. 10.1093/bioinformatics/btw313

Guelfi, S., Botia, J. A., Thom, M., Ramasamy, A., Perona, M., Stanyer, L., Martinian, L., Trabzuni, D., Smith, C., Walker, R., Ryten, M., Reimers, M., Weale, M. E., Hardy, J., & Matarin, M. (2019). Transcriptomic and genetic analyses reveal potential causal drivers for intractable partial epilepsy. Brain, 142(6), 1616–1630. 10.1093/brain/awz074

Hansen, K. F., Sakamoto, K., Pelz, C., Impey, S., & Obrietan, K. (2014). Profiling status epilepticus-induced changes in hippocampal RNA expression using high-throughput RNA sequencing. Sci Rep, 4, 6930. 10.1038/srep06930

Hawkins, N. A., & Kearney, J. A. (2012). Confirmation of an epilepsy modifier locus on mouse chromosome 11 and candidate gene analysis by RNA-Seq. Genes Brain Behav, 11(4), 452–460. 10.1111/j.1601-183X.2012.00790.x

Heinz, S., Benner, C., Spann, N., Bertolino, E., Lin, Y. C., Laslo, P., Cheng, J. X., Murre, C., Singh, H., & Glass, C. K. (2010). Simple combinations of lineage-determining transcription factors prime cis-regulatory elements required for macrophage and B cell identities. Mol Cell, 38(4), 576–589. 10.1016/j.molcel.2010.05.004

Hunsberger, J. G., Bennett, A. H., Selvanayagam, E., Duman, R. S., & Newton, S. S. (2005). Gene profiling the response to kainic acid induced seizures. Brain Res Mol Brain Res, 141(1), 95–112. 10.1016/j.molbrainres.2005.08.005

Jamali, S., Bartolomei, F., Robaglia-Schlupp, A., Massacrier, A., Peragut, J. C., Regis, J., Dufour, H., Ravid, R., Roll, P., Pereira, S., Royer, B., Roeckel-Trevisiol, N., Fontaine, M., Guye, M., Boucraut, J., Chauvel, P., Cau, P., & Szepetowski, P. (2006). Large-scale expression study of human mesial temporal lobe epilepsy: evidence for dysregulation of the neurotransmission and complement systems in the entorhinal cortex. Brain, 129(Pt 3), 625–641. 10.1093/brain/awl001

Kane, O., McCoy, A., Jada, R., Borisov, V., Zag, L., Zag, A., Schragenheim-Rozales, K., Shalgi, R., Levy, N. S., Levy, A. P., & Marsh, E. D. (2022). Characterization of spontaneous seizures and EEG abnormalities in a mouse model of the human A350V IQSEC2 mutation and identification of a possible target for precision medicine based therapy. Epilepsy Res, 182, 106907. 10.1016/j.eplepsyres.2022.106907

Kanehisa, M., & Goto, S. (2000). KEGG: kyoto encyclopedia of genes and genomes. Nucleic Acids Res, 28(1), 27–30. 10.1093/nar/28.1.27

Kirchner, A., Bagla, S., Dachet, F., & Loeb, J. A. (2020). DUSP4 appears to be a highly localized endogenous inhibitor of epileptic signaling in human neocortex. Neurobiol Dis, 145, 105073. 10.1016/j.nbd.2020.105073

Knowles, J. K., Helbig, I., Metcalf, C. S., Lubbers, L. S., Isom, L. L., Demarest, S., Goldberg, E. M., George, A. L., Jr., Lerche, H., Weckhuysen, S., Whittemore, V., Berkovic, S. F., & Lowenstein, D. H. (2022). Precision medicine for genetic epilepsy on the horizon: Recent advances, present challenges, and suggestions for continued progress. Epilepsia, 63(10), 2461–2475. 10.1111/epi.17332

Kramer, A., Green, J., Pollard, J., Jr., & Tugendreich, S. (2014). Causal analysis approaches in Ingenuity Pathway Analysis. Bioinformatics, 30(4), 523–530. 10.1093/bioinformatics/btt703

Lee, P. R., & Fields, R. D. (2021). Activity-Dependent Gene Expression in Neurons. Neuroscientist, 27(4), 355–366. 10.1177/1073858420943515

Li, X., Wang, Q., Zhang, D. W., Wu, D., Zhang, S. W., Wei, Z. R., Chen, X., & Li, W. (2022). Hippocampus RNA Sequencing of Pentylenetetrazole-Kindled Rats and Upon Treatment of Novel Chemical Q808. Front Pharmacol, 13, 820508. 10.3389/fphar.2022.820508

Lidster, K., Jefferys, J. G., Blumcke, I., Crunelli, V., Flecknell, P., Frenguelli, B. G., Gray, W. P., Kaminski, R., Pitkanen, A., Ragan, I., Shah, M., Simonato, M., Trevelyan, A., Volk, H., Walker, M., Yates, N., & Prescott, M. J. (2016). Opportunities for improving animal welfare in rodent models of epilepsy and seizures. J Neurosci Methods, 260, 2–25. 10.1016/j.jneumeth.2015.09.007

Lillis, K. P., Wang, Z., Mail, M., Zhao, G. Q., Berdichevsky, Y., Bacskai, B., & Staley, K. J. (2015). Evolution of Network Synchronization during Early Epileptogenesis Parallels Synaptic Circuit Alterations. J Neurosci, 35(27), 9920–9934. 10.1523/JNEUROSCI.4007-14.2015

Losing, P., Niturad, C. E., Harrer, M., Reckendorf, C. M. Z., Schatz, T., Sinske, D., Lerche, H., Maljevic, S., & Knoll, B. (2017). SRF modulates seizure occurrence, activity induced gene transcription and hippocampal circuit reorganization in the mouse pilocarpine epilepsy model. Mol Brain, 10(1), 30. 10.1186/s13041-017-0310-2

Love, M. I., Huber, W., & Anders, S. (2014). Moderated estimation of fold change and dispersion for RNA-seq data with DESeq2. Genome Biol, 15(12), 550. 10.1186/s13059-014-0550-8

Mahadevan, A., Codadu, N. K., & Parrish, R. R. (2022). Xenon LFP Analysis Platform Is a Novel Graphical User Interface for Analysis of Local Field Potential From Large-Scale MEA Recordings. Front Neurosci, 16, 904931. 10.3389/fnins.2022.904931

Matovu, D., & Cavalheiro, E. A. (2022). Differences in Evolution of Epileptic Seizures and Topographical Distribution of Tissue Damage in Selected Limbic Structures Between Male and Female Rats Submitted to the Pilocarpine Model. Front Neurol, 13, 802587. 10.3389/fneur.2022.802587

McHugh, J. C., & Delanty, N. (2008). Epidemiology and classification of epilepsy: gender comparisons. Int Rev Neurobiol, 83, 11–26. 10.1016/S0074-7742(08)00002-0

Mertz, C., Krarup, S., Jensen, C. D., Lindholm, S. E. H., Kjaer, C., Pinborg, L. H., & Bak, L. K. (2020). Aspects of cAMP Signaling in Epileptogenesis and Seizures and Its Potential as Drug Target. Neurochem Res, 45(6), 1247–1255. 10.1007/s11064-019-02853-x

Mielke, K., Brecht, S., Dorst, A., & Herdegen, T. (1999). Activity and expression of JNK1, p38 and ERK kinases, c-Jun N-terminal phosphorylation, and c-jun promoter binding in the adult rat brain following kainate-induced seizures. Neuroscience, 91(2), 471–483. 10.1016/s0306-4522(98)00667-8

Morgan, J. I., Cohen, D. R., Hempstead, J. L., & Curran, T. (1987). Mapping patterns of c-fos expression in the central nervous system after seizure. Science, 237(4811), 192–197. 10.1126/science.3037702

O’Leary, H., Vanderlinden, L., Southard, L., Castano, A., Saba, L. M., & Benke, T. A. (2020). Transcriptome analysis of rat dorsal hippocampal CA1 after an early life seizure induced by kainic acid. Epilepsy Res, 161, 106283. 10.1016/j.eplepsyres.2020.106283

Okamoto, O. K., Janjoppi, L., Bonone, F. M., Pansani, A. P., da Silva, A. V., Scorza, F. A., & Cavalheiro, E. A. (2010). Whole transcriptome analysis of the hippocampus: toward a molecular portrait of epileptogenesis. BMC Genomics, 11, 230. 10.1186/1471-2164-11-230

Orsini, A., Zara, F., & Striano, P. (2018). Recent advances in epilepsy genetics. Neurosci Lett, 667, 4–9. 10.1016/j.neulet.2017.05.014

Palmer, T. M., Gettys, T. W., Jacobson, K. A., & Stiles, G. L. (1994). Desensitization of the canine A2a adenosine receptor: delineation of multiple processes. Mol Pharmacol, 45(6), 1082–1094. https://www.ncbi.nlm.nih.gov/pubmed/8022402

Papasavvas, C. A., Parrish, R. R., & Trevelyan, A. J. (2020). Propagating Activity in Neocortex, Mediated by Gap Junctions and Modulated by Extracellular Potassium. eNeuro, 7(2). 10.1523/ENEURO.0387-19.2020

Parrish, R. R., Codadu, N. K., Racca, C., & Trevelyan, A. J. (2018). Pyramidal cell activity levels affect the polarity of activity-induced gene transcription changes in interneurons. J Neurophysiol, 120(5), 2358–2367. 10.1152/jn.00287.2018

Pitkanen, A., Lukasiuk, K., Dudek, F. E., & Staley, K. J. (2015). Epileptogenesis. Cold Spring Harb Perspect Med, 5(10). 10.1101/cshperspect.a022822

Ryley Parrish, R., Albertson, A. J., Buckingham, S. C., Hablitz, J. J., Mascia, K. L., Davis Haselden, W., & Lubin, F. D. (2013). Status epilepticus triggers early and late alterations in brain-derived neurotrophic factor and NMDA glutamate receptor Grin2b DNA methylation levels in the hippocampus. Neuroscience, 248, 602–619. 10.1016/j.neuroscience.2013.06.029

Shan, W., Nagai, T., Tanaka, M., Itoh, N., Furukawa-Hibi, Y., Nabeshima, T., Sokabe, M., & Yamada, K. (2018). Neuronal PAS domain protein 4 (Npas4) controls neuronal homeostasis in pentylenetetrazole-induced epilepsy through the induction of Homer1a. J Neurochem, 145(1), 19–33. 10.1111/jnc.14274

Sheng, H. Z., Fields, R. D., & Nelson, P. G. (1993). Specific regulation of immediate early genes by patterned neuronal activity. J Neurosci Res, 35(5), 459–467. 10.1002/jnr.490350502

Sheth, S., Brito, R., Mukherjea, D., Rybak, L. P., & Ramkumar, V. (2014). Adenosine receptors: expression, function and regulation. Int J Mol Sci, 15(2), 2024–2052. 10.3390/ijms15022024

Steinlein, O. K. (2014). Calcium signaling and epilepsy. Cell Tissue Res, 357(2), 385–393. 10.1007/s00441-014-1849-1

Striano, P., & Minassian, B. A. (2020). From Genetic Testing to Precision Medicine in Epilepsy. Neurotherapeutics, 17(2), 609–615. 10.1007/s13311-020-00835-4

Sun, Q., Xu, W., Piao, J., Su, J., Ge, T., Cui, R., Yang, W., & Li, B. (2022). Transcription factors are potential therapeutic targets in epilepsy. J Cell Mol Med, 26(19), 4875–4885. 10.1111/jcmm.17518

Thouta, S., Waldbrook, M. G., Lin, S., Mahadevan, A., Mezeyova, J., Soriano, M., Versi, P., Goodchild, S. J., & Parrish, R. R. (2022). Pharmacological determination of the fractional block of Nav channels required to impair neuronal excitability and ex vivo seizures. Front Cell Neurosci, 16, 964691. 10.3389/fncel.2022.964691

Trevelyan, A. J., Graham, R. T., Parrish, R. R., & Codadu, N. K. (2023). Synergistic Positive Feedback Mechanisms Underlying Seizure Initiation. Epilepsy Curr, 23(1), 38–43. 10.1177/15357597221127163

Wang, D., Ren, M., Guo, J., Yang, G., Long, X., Hu, R., Shen, W., Wang, X., & Zeng, K. (2014). The inhibitory effects of Npas4 on seizures in pilocarpine-induced epileptic rats. PLoS One, 9(12), e115801. 10.1371/journal.pone.0115801

Wang, G., Zhu, Z., Xu, D., & Sun, L. (2020). Advances in Understanding CREB Signaling-Mediated Regulation of the Pathogenesis and Progression of Epilepsy. Clin Neurol Neurosurg, 196, 106018. 10.1016/j.clineuro.2020.106018

Wolinski, P., Ksiazek-Winiarek, D., & Glabinski, A. (2022). Cytokines and Neurodegeneration in Epileptogenesis. Brain Sci, 12(3). 10.3390/brainsci12030380

Yu, G. (2018). Using meshes for MeSH term enrichment and semantic analyses. Bioinformatics, 34(21), 3766–3767. 10.1093/bioinformatics/bty410

Yu, G., Wang, L. G., Han, Y., & He, Q. Y. (2012). clusterProfiler: an R package for comparing biological themes among gene clusters. OMICS, 16(5), 284–287. 10.1089/omi.2011.0118

Zhang, C. L., Dreier, J. P., & Heinemann, U. (1995). Paroxysmal epileptiform discharges in temporal lobe slices after prolonged exposure to low magnesium are resistant to clinically used anticonvulsants. Epilepsy Res, 20(2), 105–111. 10.1016/0920-1211(94)00067-7

Zhang, L., Li, Y., Ye, X., & Bian, L. (2018). Bioinformatics Analysis of Microarray Profiling Identifies That the miR-203-3p Target Ppp2ca Aggravates Seizure Activity in Mice. J Mol Neurosci, 66(1), 146–154. 10.1007/s12031-018-1145-8

Zhou, Z., Liu, T., Sun, X., Mu, X., Zhu, G., Xiao, T., Zhao, M., & Zhao, C. (2017). CXCR4 antagonist AMD3100 reverses the neurogenesis promoted by enriched environment and suppresses long-term seizure activity in adult rats of temporal lobe epilepsy. Behav Brain Res, 322(Pt A), 83–91. 10.1016/j.bbr.2017.01.014

Zhu, Y., Huang, D., Zhao, Z., & Lu, C. (2021). Bioinformatic analysis identifies potential key genes of epilepsy. PLoS One, 16(9), e0254326. 10.1371/journal.pone.0254326

