## Supplementary File 1 for "RNA sequencing demonstrates *ex vivo* neocortical transcriptomic changes induced by epileptiform activity in male and female mice": Supplementary File 1.html

Analysis\_SeizureInduction


### Analysis\_SeizureInduction

```
dir.create("reference")
```

```
Warning in dir.create("reference"): 'reference' already exists
```

```
dir.create("fastQC_files")
```

```
Warning in dir.create("fastQC_files"): 'fastQC_files' already exists
```

```
dir.create('bam_files')
```

```
Warning in dir.create("bam_files"): 'bam_files' already exists
```

```
genome_url <- "https://ftp.ensembl.org/pub/release-108/fasta/mus_musculus/dna/Mus_musculus.GRCm39.dna_sm.toplevel.fa.gz" 
genome_file <- "Mus_musculus.GRCm39.dna_sm.toplevel.fa.gz" 
annot_url <- "https://ftp.ensembl.org/pub/release-108/gtf/mus_musculus/Mus_musculus.GRCm39.108.gtf.gz"
annot_file <- "Mus_musculus.GRCm39.108.gtf.gz"

options(timeout = max(8000, getOption("timeout")))

#download.file(genome_url, genome_file)
#download.file(annot_url, annot_file)

library(Rsubread)

#buildindex("genomefile", 
           #genome_file, 
           #indexSplit = TRUE, 
           #gappedIndex = TRUE, 
           #memory = 2000)
```

FASTQC

```
fastq_dir <- "Seq_Files"  # The folder with the sequencing data
qc_dir <- "fastQC_files"  # The results folder

# Load the library
library(fastqcr)
#fastqc_install()
# Run the QC
#fastqc(fastq_dir, qc_dir)
```

Alignment

```
# Set variables
annot <- "Mus_musculus.GRCm39.108.gtf.gz"  # Filename for gtf file
fastq_dir <- "Seq_Files"  # Folder where fastq files are saved
bname <- "genomefile"  # The name you gave your genome index
library(Rsubread)
library(Rsamtools)
```

```
Loading required package: GenomeInfoDb
```

```
Loading required package: BiocGenerics
```

```
Attaching package: 'BiocGenerics'
```

```
The following objects are masked from 'package:stats':

    IQR, mad, sd, var, xtabs
```

```
The following objects are masked from 'package:base':

    anyDuplicated, aperm, append, as.data.frame, basename, cbind,
    colnames, dirname, do.call, duplicated, eval, evalq, Filter, Find,
    get, grep, grepl, intersect, is.unsorted, lapply, Map, mapply,
    match, mget, order, paste, pmax, pmax.int, pmin, pmin.int,
    Position, rank, rbind, Reduce, rownames, sapply, setdiff, sort,
    table, tapply, union, unique, unsplit, which.max, which.min
```

```
Loading required package: S4Vectors
```

```
Loading required package: stats4
```

```
Attaching package: 'S4Vectors'
```

```
The following object is masked from 'package:utils':

    findMatches
```

```
The following objects are masked from 'package:base':

    expand.grid, I, unname
```

```
Loading required package: IRanges
```

```
Loading required package: GenomicRanges
```

```
Loading required package: Biostrings
```

```
Loading required package: XVector
```

```
Attaching package: 'Biostrings'
```

```
The following object is masked from 'package:base':

    strsplit
```

```
# Get file names
fastq_files <- list.files(
  path = fastq_dir,
  pattern = ".fastq.gz",
  full.names = TRUE)

files2 <- list.files(
  path = fastq_dir,
  pattern = "28|29|30",
  full.names = TRUE
)

mat2 <- matrix(data= files2, ncol =2, byrow= TRUE)
mat2
```

```
     [,1]                                   
[1,] "Seq_Files/RP-28-UAB7_R1.001.fastq.gz" 
[2,] "Seq_Files/RP-29-UAB10_R1_001.fastq.gz"
[3,] "Seq_Files/RP-30-UAB22_R1_001.fastq.gz"
     [,2]                                   
[1,] "Seq_Files/RP-28-UAB7_R2_001.fastq.gz" 
[2,] "Seq_Files/RP-29-UAB10_R2_001.fastq.gz"
[3,] "Seq_Files/RP-30-UAB22_R2_001.fastq.gz"
```

```
# Make matrix of file names
fastq_files_mat <- matrix(data = fastq_files, ncol = 2, byrow = TRUE)
fastq_files_mat
```

```
      [,1]                                   
 [1,] "Seq_Files/RP-17-UAB17_R1_001.fastq.gz"
 [2,] "Seq_Files/RP-18-UAB20_R1_001.fastq.gz"
 [3,] "Seq_Files/RP-19-UAB26_R1_001.fastq.gz"
 [4,] "Seq_Files/RP-20-UAB8_R1_001.fastq.gz" 
 [5,] "Seq_Files/RP-21-UAB11_R1_001.fastq.gz"
 [6,] "Seq_Files/RP-22-UAB23_R1_001.fastq.gz"
 [7,] "Seq_Files/RP-23-UAB3_R1_001.fastq.gz" 
 [8,] "Seq_Files/RP-24-UAB13_R1_001.fastq.gz"
 [9,] "Seq_Files/RP-25-UAB16_R1_001.fastq.gz"
[10,] "Seq_Files/RP-26-UAB19_R1_001.fastq.gz"
[11,] "Seq_Files/RP-27-UAB25_R1_001.fastq.gz"
[12,] "Seq_Files/RP-28-UAB7_R1.001.fastq.gz" 
[13,] "Seq_Files/RP-29-UAB10_R1_001.fastq.gz"
[14,] "Seq_Files/RP-30-UAB22_R1_001.fastq.gz"
      [,2]                                   
 [1,] "Seq_Files/RP-17-UAB17_R2_001.fastq.gz"
 [2,] "Seq_Files/RP-18-UAB20_R2_001.fastq.gz"
 [3,] "Seq_Files/RP-19-UAB26_R2_001.fastq.gz"
 [4,] "Seq_Files/RP-20-UAB8_R2_001.fastq.gz" 
 [5,] "Seq_Files/RP-21-UAB11_R2_001.fastq.gz"
 [6,] "Seq_Files/RP-22-UAB23_R2_001.fastq.gz"
 [7,] "Seq_Files/RP-23-UAB3_R2_001.fastq.gz" 
 [8,] "Seq_Files/RP-24-UAB13_R2_001.fastq.gz"
 [9,] "Seq_Files/RP-25-UAB16_R2_001.fastq.gz"
[10,] "Seq_Files/RP-26-UAB19_R2_001.fastq.gz"
[11,] "Seq_Files/RP-27-UAB25_R2_001.fastq.gz"
[12,] "Seq_Files/RP-28-UAB7_R2_001.fastq.gz" 
[13,] "Seq_Files/RP-29-UAB10_R2_001.fastq.gz"
[14,] "Seq_Files/RP-30-UAB22_R2_001.fastq.gz"
```

```
# Create alignment function
# This function will help you create output file names.
#RS_align <- function(read_file) {
  #basename <- gsub(".*/(.*?)_R1.001.fastq.gz", "\\1", 
                #read_file[1])

  # Run alignment
  #subjunc(
    #bname,
    #readfile1 = read_file[1],
    #readfile2 = read_file[2],
    #output_file = paste("~/RileyProj/bam_files/", basename, ".bam", sep = ""),
    #annot.ext = annot_file,
    #useAnnotation = TRUE,
    #isGTF = TRUE,
    #nthreads = 2)
  
  # Sort files
  #sorted <- sortBam(paste("bam_files/", basename, ".bam", sep = ""), 
                    #paste("bam_files/", basename, ".sorted", sep = ""))
  # Index files
  #indexBam(sorted)


# Run apply to use the alignment function on rows of file names
#apply(fastq_files_mat, 1, RS_align)
```

Count Table

```
library(Rsubread)
annot_file <- 'Mus_musculus.GRCm39.108.gtf.gz'
# Create vector of bam files
filepath1<- "bam_files/non_sorted"
bam_files <- list.files(filepath1, full.names= TRUE)
bam_files
```

```
 [1] "bam_files/non_sorted/RP-17-UAB17.bam"
 [2] "bam_files/non_sorted/RP-18-UAB20.bam"
 [3] "bam_files/non_sorted/RP-19-UAB26.bam"
 [4] "bam_files/non_sorted/RP-20-UAB8.bam" 
 [5] "bam_files/non_sorted/RP-21-UAB11.bam"
 [6] "bam_files/non_sorted/RP-22-UAB23.bam"
 [7] "bam_files/non_sorted/RP-23-UAB3.bam" 
 [8] "bam_files/non_sorted/RP-24-UAB13.bam"
 [9] "bam_files/non_sorted/RP-25-UAB16.bam"
[10] "bam_files/non_sorted/RP-26-UAB19.bam"
[11] "bam_files/non_sorted/RP-27-UAB25.bam"
[12] "bam_files/non_sorted/RP-28-UAB7.bam" 
[13] "bam_files/non_sorted/RP-29-UAB10.bam"
[14] "bam_files/non_sorted/RP-30-UAB22.bam"
```

```
# Set sample names
samples <- c("ACSF-1M", "ACSF-2M", "ACSF-3M","ACSF-4F", "ACSF-5F","ACSF-6F","4AP-1M","4AP-2M","4AP-3M","4AP-4M", "4AP-5M", "4AP-6F", "4AP-7F", "4AP-8F")
# Assign features
genecounts <- featureCounts(files = bam_files,
                            annot.ext = annot_file,
                          isGTFAnnotationFile = TRUE,
                            isPairedEnd= TRUE)
```

```
        ==========     _____ _    _ ____  _____  ______          _____  
        =====         / ____| |  | |  _ \|  __ \|  ____|   /\   |  __ \ 
          =====      | (___ | |  | | |_) | |__) | |__     /  \  | |  | |
            ====      \___ \| |  | |  _ <|  _  /|  __|   / /\ \ | |  | |
              ====    ____) | |__| | |_) | | \ \| |____ / ____ \| |__| |
        ==========   |_____/ \____/|____/|_|  \_\______/_/    \_\_____/
       Rsubread 2.14.2

//========================== featureCounts setting ===========================\\
||                                                                            ||
||             Input files : 14 BAM files                                     ||
||                                                                            ||
||                           RP-17-UAB17.bam                                  ||
||                           RP-18-UAB20.bam                                  ||
||                           RP-19-UAB26.bam                                  ||
||                           RP-20-UAB8.bam                                   ||
||                           RP-21-UAB11.bam                                  ||
||                           RP-22-UAB23.bam                                  ||
||                           RP-23-UAB3.bam                                   ||
||                           RP-24-UAB13.bam                                  ||
||                           RP-25-UAB16.bam                                  ||
||                           RP-26-UAB19.bam                                  ||
||                           RP-27-UAB25.bam                                  ||
||                           RP-28-UAB7.bam                                   ||
||                           RP-29-UAB10.bam                                  ||
||                           RP-30-UAB22.bam                                  ||
||                                                                            ||
||              Paired-end : yes                                              ||
||        Count read pairs : yes                                              ||
||              Annotation : Mus_musculus.GRCm39.108.gtf.gz (GTF)             ||
||      Dir for temp files : .                                                ||
||                 Threads : 1                                                ||
||                   Level : meta-feature level                               ||
||      Multimapping reads : counted                                          ||
|| Multi-overlapping reads : not counted                                      ||
||   Min overlapping bases : 1                                                ||
||                                                                            ||
\\============================================================================//

//================================= Running ==================================\\
||                                                                            ||
|| Load annotation file Mus_musculus.GRCm39.108.gtf.gz ...                    ||
||    Features : 868961                                                       ||
||    Meta-features : 56980                                                   ||
||    Chromosomes/contigs : 39                                                ||
||                                                                            ||
|| Process BAM file RP-17-UAB17.bam...                                        ||
||    Paired-end reads are included.                                          ||
||    Total alignments : 12807868                                             ||
||    Successfully assigned alignments : 10767396 (84.1%)                     ||
||    Running time : 1.00 minutes                                             ||
||                                                                            ||
|| Process BAM file RP-18-UAB20.bam...                                        ||
||    Paired-end reads are included.                                          ||
||    Total alignments : 14026817                                             ||
||    Successfully assigned alignments : 11781789 (84.0%)                     ||
||    Running time : 1.07 minutes                                             ||
||                                                                            ||
|| Process BAM file RP-19-UAB26.bam...                                        ||
||    Paired-end reads are included.                                          ||
||    Total alignments : 13235987                                             ||
||    Successfully assigned alignments : 11136904 (84.1%)                     ||
||    Running time : 1.01 minutes                                             ||
||                                                                            ||
|| Process BAM file RP-20-UAB8.bam...                                         ||
||    Paired-end reads are included.                                          ||
||    Total alignments : 6931871                                              ||
||    Successfully assigned alignments : 5746554 (82.9%)                      ||
||    Running time : 0.54 minutes                                             ||
||                                                                            ||
|| Process BAM file RP-21-UAB11.bam...                                        ||
||    Paired-end reads are included.                                          ||
||    Total alignments : 15923567                                             ||
||    Successfully assigned alignments : 12915270 (81.1%)                     ||
||    Running time : 1.87 minutes                                             ||
||                                                                            ||
|| Process BAM file RP-22-UAB23.bam...                                        ||
||    Paired-end reads are included.                                          ||
||    Total alignments : 9639553                                              ||
||    Successfully assigned alignments : 8145758 (84.5%)                      ||
||    Running time : 0.74 minutes                                             ||
||                                                                            ||
|| Process BAM file RP-23-UAB3.bam...                                         ||
||    Paired-end reads are included.                                          ||
||    Total alignments : 10348296                                             ||
||    Successfully assigned alignments : 8609919 (83.2%)                      ||
||    Running time : 0.79 minutes                                             ||
||                                                                            ||
|| Process BAM file RP-24-UAB13.bam...                                        ||
||    Paired-end reads are included.                                          ||
||    Total alignments : 12203872                                             ||
||    Successfully assigned alignments : 10302900 (84.4%)                     ||
||    Running time : 0.95 minutes                                             ||
||                                                                            ||
|| Process BAM file RP-25-UAB16.bam...                                        ||
||    Paired-end reads are included.                                          ||
||    Total alignments : 12491116                                             ||
||    Successfully assigned alignments : 10430318 (83.5%)                     ||
||    Running time : 0.99 minutes                                             ||
||                                                                            ||
|| Process BAM file RP-26-UAB19.bam...                                        ||
||    Paired-end reads are included.                                          ||
||    Total alignments : 11183149                                             ||
||    Successfully assigned alignments : 9320555 (83.3%)                      ||
||    Running time : 0.89 minutes                                             ||
||                                                                            ||
|| Process BAM file RP-27-UAB25.bam...                                        ||
||    Paired-end reads are included.                                          ||
||    Total alignments : 10678762                                             ||
||    Successfully assigned alignments : 9079534 (85.0%)                      ||
||    Running time : 0.85 minutes                                             ||
||                                                                            ||
|| Process BAM file RP-28-UAB7.bam...                                         ||
||    Paired-end reads are included.                                          ||
||    Total alignments : 10806041                                             ||
||    Successfully assigned alignments : 9120556 (84.4%)                      ||
||    Running time : 0.86 minutes                                             ||
||                                                                            ||
|| Process BAM file RP-29-UAB10.bam...                                        ||
||    Paired-end reads are included.                                          ||
||    Total alignments : 10415361                                             ||
||    Successfully assigned alignments : 8832251 (84.8%)                      ||
||    Running time : 0.82 minutes                                             ||
||                                                                            ||
|| Process BAM file RP-30-UAB22.bam...                                        ||
||    Paired-end reads are included.                                          ||
||    Total alignments : 8963579                                              ||
||    Successfully assigned alignments : 7579449 (84.6%)                      ||
||    Running time : 0.70 minutes                                             ||
||                                                                            ||
|| Write the final count table.                                               ||
|| Write the read assignment summary.                                         ||
||                                                                            ||
\\============================================================================//
```

```
# Extract count table
counts<- genecounts$counts
colnames(counts)<- samples
# Set column name
head(counts)
```

```
                   ACSF-1M ACSF-2M ACSF-3M ACSF-4F ACSF-5F ACSF-6F 4AP-1M
ENSMUSG00000102628       4       2       1       1       3       0      0
ENSMUSG00000100595       0       0       0       0       0       0      0
ENSMUSG00000097426       0       0       0       0       4       0      0
ENSMUSG00000104478       0       0       0       0       0       0      0
ENSMUSG00000104385       0       0       0       0       0       0      0
ENSMUSG00000086053       7       4       2       5       4       2      2
                   4AP-2M 4AP-3M 4AP-4M 4AP-5M 4AP-6F 4AP-7F 4AP-8F
ENSMUSG00000102628      0      0      2      0      2      1      2
ENSMUSG00000100595      0      0      0      0      0      0      0
ENSMUSG00000097426      0      0      0      0      0      0      0
ENSMUSG00000104478      0      0      0      0      0      0      0
ENSMUSG00000104385      0      0      0      0      0      0      0
ENSMUSG00000086053      2      2      1      2      2      1      2
```

```
library(DESeq2)
```

```
Loading required package: SummarizedExperiment
```

```
Loading required package: MatrixGenerics
```

```
Loading required package: matrixStats
```

```
Attaching package: 'MatrixGenerics'
```

```
The following objects are masked from 'package:matrixStats':

    colAlls, colAnyNAs, colAnys, colAvgsPerRowSet, colCollapse,
    colCounts, colCummaxs, colCummins, colCumprods, colCumsums,
    colDiffs, colIQRDiffs, colIQRs, colLogSumExps, colMadDiffs,
    colMads, colMaxs, colMeans2, colMedians, colMins, colOrderStats,
    colProds, colQuantiles, colRanges, colRanks, colSdDiffs, colSds,
    colSums2, colTabulates, colVarDiffs, colVars, colWeightedMads,
    colWeightedMeans, colWeightedMedians, colWeightedSds,
    colWeightedVars, rowAlls, rowAnyNAs, rowAnys, rowAvgsPerColSet,
    rowCollapse, rowCounts, rowCummaxs, rowCummins, rowCumprods,
    rowCumsums, rowDiffs, rowIQRDiffs, rowIQRs, rowLogSumExps,
    rowMadDiffs, rowMads, rowMaxs, rowMeans2, rowMedians, rowMins,
    rowOrderStats, rowProds, rowQuantiles, rowRanges, rowRanks,
    rowSdDiffs, rowSds, rowSums2, rowTabulates, rowVarDiffs, rowVars,
    rowWeightedMads, rowWeightedMeans, rowWeightedMedians,
    rowWeightedSds, rowWeightedVars
```

```
Loading required package: Biobase
```

```
Welcome to Bioconductor

    Vignettes contain introductory material; view with
    'browseVignettes()'. To cite Bioconductor, see
    'citation("Biobase")', and for packages 'citation("pkgname")'.
```

```
Attaching package: 'Biobase'
```

```
The following object is masked from 'package:MatrixGenerics':

    rowMedians
```

```
The following objects are masked from 'package:matrixStats':

    anyMissing, rowMedians
```

```
# Make data frame 
colData <- data.frame(Treatment = factor(c("ACSF", "ACSF", "ACSF", "ACSF", "ACSF","ACSF","4-AP","4-AP","4-AP","4-AP","4-AP","4-AP","4-AP","4-AP"), levels = c("ACSF", "4-AP")), 
    Gender = c("Male", "Male", "Male", "Female","Female", "Female","Male","Male","Male", "Male", "Male", "Female", "Female", "Female"), row.names=samples)
colData
```

```
        Treatment Gender
ACSF-1M      ACSF   Male
ACSF-2M      ACSF   Male
ACSF-3M      ACSF   Male
ACSF-4F      ACSF Female
ACSF-5F      ACSF Female
ACSF-6F      ACSF Female
4AP-1M       4-AP   Male
4AP-2M       4-AP   Male
4AP-3M       4-AP   Male
4AP-4M       4-AP   Male
4AP-5M       4-AP   Male
4AP-6F       4-AP Female
4AP-7F       4-AP Female
4AP-8F       4-AP Female
```

```
head(counts)
```

```
                   ACSF-1M ACSF-2M ACSF-3M ACSF-4F ACSF-5F ACSF-6F 4AP-1M
ENSMUSG00000102628       4       2       1       1       3       0      0
ENSMUSG00000100595       0       0       0       0       0       0      0
ENSMUSG00000097426       0       0       0       0       4       0      0
ENSMUSG00000104478       0       0       0       0       0       0      0
ENSMUSG00000104385       0       0       0       0       0       0      0
ENSMUSG00000086053       7       4       2       5       4       2      2
                   4AP-2M 4AP-3M 4AP-4M 4AP-5M 4AP-6F 4AP-7F 4AP-8F
ENSMUSG00000102628      0      0      2      0      2      1      2
ENSMUSG00000100595      0      0      0      0      0      0      0
ENSMUSG00000097426      0      0      0      0      0      0      0
ENSMUSG00000104478      0      0      0      0      0      0      0
ENSMUSG00000104385      0      0      0      0      0      0      0
ENSMUSG00000086053      2      2      1      2      2      1      2
```

DESeq

```
library(DESeq2)
# Create DESeq dataset from the count matrix
dds <- DESeqDataSetFromMatrix(countData = counts,
                              colData = colData,
                              design = ~Treatment+ Gender+ Treatment:Gender)
```

```
Warning in DESeqDataSet(se, design = design, ignoreRank): some variables in
design formula are characters, converting to factors
```

```
  Note: levels of factors in the design contain characters other than
  letters, numbers, '_' and '.'. It is recommended (but not required) to use
  only letters, numbers, and delimiters '_' or '.', as these are safe characters
  for column names in R. [This is a message, not a warning or an error]
```

```
# Run DESeq
dds <- DESeq(dds)
```

```
estimating size factors
  Note: levels of factors in the design contain characters other than
  letters, numbers, '_' and '.'. It is recommended (but not required) to use
  only letters, numbers, and delimiters '_' or '.', as these are safe characters
  for column names in R. [This is a message, not a warning or an error]
```

```
estimating dispersions
```

```
gene-wise dispersion estimates
```

```
mean-dispersion relationship
```

```
  Note: levels of factors in the design contain characters other than
  letters, numbers, '_' and '.'. It is recommended (but not required) to use
  only letters, numbers, and delimiters '_' or '.', as these are safe characters
  for column names in R. [This is a message, not a warning or an error]
```

```
final dispersion estimates
```

```
  Note: levels of factors in the design contain characters other than
  letters, numbers, '_' and '.'. It is recommended (but not required) to use
  only letters, numbers, and delimiters '_' or '.', as these are safe characters
  for column names in R. [This is a message, not a warning or an error]
```

```
fitting model and testing
```

```
resultsNames(dds)
```

```
[1] "Intercept"                "Treatment_4.AP_vs_ACSF"  
[3] "Gender_Male_vs_Female"    "Treatment4.AP.GenderMale"
```

```
head(dds)
```

```
class: DESeqDataSet 
dim: 6 14 
metadata(1): version
assays(4): counts mu H cooks
rownames(6): ENSMUSG00000102628 ENSMUSG00000100595 ...
  ENSMUSG00000104385 ENSMUSG00000086053
rowData names(30): baseMean baseVar ... deviance maxCooks
colnames(14): ACSF-1M ACSF-2M ... 4AP-7F 4AP-8F
colData names(3): Treatment Gender sizeFactor
```

```
# Get results and order the tables
#Treatment
dds_results_treatment <- results(object = dds,
                       contrast = c("Treatment", "4-AP", "ACSF" ))

dds_results_treatment <- dds_results_treatment[order(abs(dds_results_treatment$log2FoldChange), decreasing = TRUE, na.last = TRUE), ]

#Gender
dds_results_gender <- results(object = dds,
                       contrast = c("Gender", "Male", "Female" ))

dds_results_gender <- dds_results_gender[order(abs(dds_results_gender$log2FoldChange), decreasing = TRUE, na.last = TRUE), ]

#Interacting Factor
dds_results_interaction <- results(object = dds,
                       name = c("Treatment4.AP.GenderMale" ))

dds_results_interaction <- dds_results_interaction[order(abs(dds_results_interaction$log2FoldChange), decreasing = TRUE, na.last = TRUE), ]

# Print the first 20 rows
head(dds_results_treatment)
```

```
log2 fold change (MLE): Treatment 4-AP vs ACSF 
Wald test p-value: Treatment 4.AP vs ACSF 
DataFrame with 6 rows and 6 columns
                    baseMean log2FoldChange     lfcSE      stat     pvalue
                   <numeric>      <numeric> <numeric> <numeric>  <numeric>
ENSMUSG00000111530   2.32357       -5.57128   3.11017  -1.79131 0.07324377
ENSMUSG00000097833   1.69143       -5.10144   2.89215  -1.76389 0.07774979
ENSMUSG00000097146   2.56931        4.68838   1.51150   3.10180 0.00192348
ENSMUSG00000108694   1.82536       -4.60475   1.68188  -2.73786 0.00618397
ENSMUSG00000100794  11.32651       -4.57835   1.37650  -3.32609         NA
ENSMUSG00000034224   3.24258       -4.54855   1.59749  -2.84730 0.00440915
                        padj
                   <numeric>
ENSMUSG00000111530        NA
ENSMUSG00000097833        NA
ENSMUSG00000097146        NA
ENSMUSG00000108694        NA
ENSMUSG00000100794        NA
ENSMUSG00000034224        NA
```

```
head(dds_results_gender, 20)
```

```
log2 fold change (MLE): Gender Male vs Female 
Wald test p-value: Gender Male vs Female 
DataFrame with 20 rows and 6 columns
                    baseMean log2FoldChange     lfcSE      stat      pvalue
                   <numeric>      <numeric> <numeric> <numeric>   <numeric>
ENSMUSG00000069049  176.2719       10.59004  1.184013   8.94420 3.74668e-19
ENSMUSG00000069045  296.4364        9.51239  1.010268   9.41570 4.69924e-21
ENSMUSG00000056673   73.5126        9.30821  1.190813   7.81668 5.42328e-15
ENSMUSG00000086503  282.0885       -8.65391  0.605227 -14.29862 2.23185e-46
ENSMUSG00000068457   82.9278        8.63552  1.171654   7.37037 1.70157e-13
...                      ...            ...       ...       ...         ...
ENSMUSG00000114087  1.360500       -4.43129   1.76035 -2.517280  0.01182648
ENSMUSG00000081772  1.980015       -4.43050   1.58796 -2.790059  0.00526985
ENSMUSG00000117856  2.229023       -4.41095   1.55854 -2.830173  0.00465229
ENSMUSG00000098102  1.704215       -4.39898   1.72335 -2.552582  0.01069277
ENSMUSG00000097920  0.729733       -4.38020   4.58968 -0.954358  0.33990231
                          padj
                     <numeric>
ENSMUSG00000069049 2.11500e-15
ENSMUSG00000069045 3.97908e-17
ENSMUSG00000056673 2.29608e-11
ENSMUSG00000086503 3.77963e-42
ENSMUSG00000068457 5.76323e-10
...                        ...
ENSMUSG00000114087          NA
ENSMUSG00000081772          NA
ENSMUSG00000117856          NA
ENSMUSG00000098102          NA
ENSMUSG00000097920          NA
```

```
head(dds_results_interaction, 20)
```

```
log2 fold change (MLE): Treatment4.AP.GenderMale 
Wald test p-value: Treatment4.AP.GenderMale 
DataFrame with 20 rows and 6 columns
                     baseMean log2FoldChange     lfcSE      stat     pvalue
                    <numeric>      <numeric> <numeric> <numeric>  <numeric>
ENSMUSG00000061808 121.815740       11.44496   4.87267   2.34881 0.01883362
ENSMUSG00000054932   1.032980        6.76773   2.51300   2.69308 0.00707946
ENSMUSG00000083043   1.409940        6.63199   2.51385   2.63819 0.00833508
ENSMUSG00000118311   1.044146       -6.50263   2.42811  -2.67806 0.00740493
ENSMUSG00000102732   0.883679       -6.47866   2.79001  -2.32209 0.02022814
...                       ...            ...       ...       ...        ...
ENSMUSG00000120620   0.688733        5.63092   3.18424   1.76837 0.07699867
ENSMUSG00000118144   1.804887        5.62682   2.18514   2.57504 0.01002276
ENSMUSG00000097833   1.691431        5.56426   4.00946   1.38778 0.16520319
ENSMUSG00000108860   3.160107        5.55464   2.02031   2.74939 0.00597055
ENSMUSG00000096177   1.412906        5.53104   2.39631   2.30815 0.02099093
                        padj
                   <numeric>
ENSMUSG00000061808  0.999996
ENSMUSG00000054932  0.999996
ENSMUSG00000083043  0.999996
ENSMUSG00000118311  0.999996
ENSMUSG00000102732  0.999996
...                      ...
ENSMUSG00000120620  0.999996
ENSMUSG00000118144  0.999996
ENSMUSG00000097833  0.999996
ENSMUSG00000108860  0.999996
ENSMUSG00000096177  0.999996
```

```
#CSV Files 
write.csv(dds_results_treatment, "dds.results.treatment.csv")
write.csv(dds_results_gender, "dds.results.gender.csv")
write.csv(dds_results_interaction, "dds.results.interaction.csv")
```

Volcano Plots

```
library(ggplot2)
library(DESeq2)

# Make PCA Plot
t <- plotPCA(rlog(dds), "Treatment") + geom_text(aes(label=samples),vjust=2,check_overlap = FALSE,size = 4, )
t <- t+theme(text = element_text(size = 24))
t
```

```
png('pca.treat.png', res= 250, height = 2400, width=2000)
print(t)

gen <- plotPCA(rlog(dds), "Gender") + geom_text(aes(label=samples),vjust=2,check_overlap = FALSE,size = 4)
gen <- gen+ theme(text = element_text(size = 24))
png('pca.gender.png', res= 300, height = 2400, width=2400)
print(gen)

# Make MA Plot
#plotMA(dds_results)
# Make Volcano Plot Treatment FIX SAVING THING 
gtreat<- ggplot(data.frame(dds_results_treatment), aes(x= log2FoldChange, y= -10*log10(padj))) + 
  geom_point() + 
  geom_vline(xintercept= c(-0.585, .585), color= 'blue') + 
  geom_hline(yintercept= 13, color= 'red')+
  theme(text = element_text(size = 24))
gtreat
```

```
Warning: Removed 39306 rows containing missing values (`geom_point()`).
```

```
#ggsave("volcano.treatment.png", gtreat)
png('volcano.treat.png', res= 250, height = 2500, width=2000)
print(gtreat)
```

```
Warning: Removed 39306 rows containing missing values (`geom_point()`).
```

```
# Make Volcano Plot Sex
ggender<- ggplot(data.frame(dds_results_gender), aes(x= log2FoldChange, y= -10*log10(padj))) + 
  geom_point() + 
  geom_vline(xintercept= c(-0.585, .585), color= 'blue') + 
  geom_hline(yintercept= 13, color= 'red')+
  theme(text = element_text(size = 24))
ggender
```

```
Warning: Removed 40045 rows containing missing values (`geom_point()`).
```

```
png("volcano.gender.png", res= 250, height = 2000, width=2000 )
print(ggender)
```

```
Warning: Removed 40045 rows containing missing values (`geom_point()`).
```

```
#Volcano Interacting Factor
gint<- ggplot(data.frame(dds_results_interaction), aes(x= log2FoldChange, y= -10*log10(padj))) + 
  geom_point() + 
  geom_vline(xintercept= c(-0.585, .585), color= 'blue') + 
  geom_hline(yintercept= 13, color= 'red')+
  theme(text = element_text(size = 24))
gint
```

```
Warning: Removed 17803 rows containing missing values (`geom_point()`).
```

```
png("volcano.interaction.png", res= 250, height = 2000, width=2000 )
print(gint)
```

```
Warning: Removed 17803 rows containing missing values (`geom_point()`).
```

Treatment Heatmap

```
treatment <- as.data.frame(dds_results_treatment)
treatment <- na.omit(treatment)

genderlist <- as.data.frame(dds_results_gender)
library(org.Mm.eg.db)
```

```
Loading required package: AnnotationDbi
```

```
library(DESeq2)
library(ggplot2)
library(Rsubread)

#Getting GeneID's 
treatment$symbol <- mapIds(org.Mm.eg.db, keys = rownames(treatment), keytype = "ENSEMBL", column = "SYMBOL")
```

```
'select()' returned 1:many mapping between keys and columns
```

```
head(treatment)
```

```
                      baseMean log2FoldChange     lfcSE      stat       pvalue
ENSMUSG00000045903 6665.662737       4.145262 0.2797654 14.816922 1.138841e-49
ENSMUSG00000114708    9.122834       3.563186 0.8669118  4.110206 3.953066e-05
ENSMUSG00000091123   16.252922      -3.200839 0.7363108 -4.347131 1.379301e-05
ENSMUSG00000066197   15.275700      -3.140181 1.3137453 -2.390251 1.683687e-02
ENSMUSG00000114218    9.925000       3.037285 0.8485024  3.579583 3.441423e-04
ENSMUSG00000094955   11.083658      -3.036563 0.8095220 -3.751057 1.760904e-04
                           padj symbol
ENSMUSG00000045903 2.012788e-45  Npas4
ENSMUSG00000114708 7.762943e-03   <NA>
ENSMUSG00000091123 3.216254e-03   <NA>
ENSMUSG00000066197 3.165689e-01 Gpr139
ENSMUSG00000114218 4.028061e-02   <NA>
ENSMUSG00000094955 2.551002e-02   <NA>
```

```
#Gender list and csv 
genderlist$symbol <- mapIds(org.Mm.eg.db, keys = rownames(genderlist), keytype = "ENSEMBL", column = "SYMBOL")
```

```
'select()' returned 1:many mapping between keys and columns
```

```
head(genderlist)
```

```
                     baseMean log2FoldChange     lfcSE       stat       pvalue
ENSMUSG00000069049 176.271920      10.590040 1.1840125   8.944196 3.746685e-19
ENSMUSG00000069045 296.436444       9.512385 1.0102681   9.415704 4.699235e-21
ENSMUSG00000056673  73.512647       9.308210 1.1908131   7.816684 5.423284e-15
ENSMUSG00000086503 282.088543      -8.653909 0.6052268 -14.298622 2.231846e-46
ENSMUSG00000068457  82.927849       8.635518 1.1716536   7.370368 1.701575e-13
ENSMUSG00000099835   3.875256      -5.540706 1.5962238  -3.471134 5.182658e-04
                           padj  symbol
ENSMUSG00000069049 2.115004e-15 Eif2s3y
ENSMUSG00000069045 3.979077e-17   Ddx3y
ENSMUSG00000056673 2.296083e-11   Kdm5d
ENSMUSG00000086503 3.779631e-42    Xist
ENSMUSG00000068457 5.763234e-10     Uty
ENSMUSG00000099835           NA    <NA>
```

```
genderlist <- na.omit(genderlist)
genderlist1 <- genderlist[abs(genderlist$log2FoldChange) >0.58 & genderlist$padj < 0.05,]
genderlist1
```

```
                    baseMean log2FoldChange     lfcSE       stat       pvalue
ENSMUSG00000069049 176.27192     10.5900397 1.1840125   8.944196 3.746685e-19
ENSMUSG00000069045 296.43644      9.5123853 1.0102681   9.415704 4.699235e-21
ENSMUSG00000056673  73.51265      9.3082100 1.1908131   7.816684 5.423284e-15
ENSMUSG00000086503 282.08854     -8.6539092 0.6052268 -14.298622 2.231846e-46
ENSMUSG00000068457  82.92785      8.6355182 1.1716536   7.370368 1.701575e-13
ENSMUSG00000048489  18.79638     -2.0226477 0.4467110  -4.527867 5.958211e-06
ENSMUSG00000099997  44.67098     -1.4204745 0.3383549  -4.198179 2.690702e-05
ENSMUSG00000019232 133.31254      0.9040445 0.2140959   4.222616 2.414834e-05
ENSMUSG00000075224 271.34958      0.6574265 0.1644941   3.996657 6.424317e-05
ENSMUSG00000016494 937.20590      0.6416654 0.1444511   4.442095 8.908740e-06
ENSMUSG00000037593 248.25238      0.6291852 0.1533518   4.102889 4.080232e-05
ENSMUSG00000025795 499.20728      0.6202796 0.1448904   4.281025 1.860347e-05
ENSMUSG00000021680 252.00939     -0.6070400 0.1454817  -4.172621 3.011153e-05
ENSMUSG00000026814 236.20348      0.5877806 0.1365191   4.305482 1.666225e-05
ENSMUSG00000035150 398.18559     -0.5804787 0.1114456  -5.208626 1.902436e-07
                           padj     symbol
ENSMUSG00000069049 2.115004e-15    Eif2s3y
ENSMUSG00000069045 3.979077e-17      Ddx3y
ENSMUSG00000056673 2.296083e-11      Kdm5d
ENSMUSG00000086503 3.779631e-42       Xist
ENSMUSG00000068457 5.763234e-10        Uty
ENSMUSG00000048489 1.261279e-02      Depp1
ENSMUSG00000099997 2.983321e-02 Tubb4b-ps2
ENSMUSG00000019232 2.921086e-02     Etnppl
ENSMUSG00000075224 4.730253e-02     Lrrc55
ENSMUSG00000016494 1.676328e-02       Cd34
ENSMUSG00000037593 3.788881e-02       Rskr
ENSMUSG00000025795 2.814637e-02     Rassf3
ENSMUSG00000021680 2.999639e-02      Crhbp
ENSMUSG00000026814 2.814637e-02        Eng
ENSMUSG00000035150 5.369627e-04    Eif2s3x
```

```
#write.csv(genderlist1, 'genderlist.csv')


#Selecting genes that are significant 
top.treatment <- treatment[ treatment$baseMean > 75 & abs(treatment$log2FoldChange) > .8,]
top.treatment <- top.treatment[order(top.treatment$log2FoldChange, decreasing= TRUE),]


#Getting Z scores for heatmap
rlog_out <- rlog(dds, blind= FALSE)# getting counts
countmat <- assay(rlog_out)[rownames(top.treatment), rownames(colData)]
base_mean <- rowMeans(countmat)
scaled_mat <- t(apply(countmat, 1, scale))
colnames(scaled_mat) <- rownames(colData)
scaled_mat
```

```
                        ACSF-1M      ACSF-2M      ACSF-3M      ACSF-4F
ENSMUSG00000045903 -1.091421339 -1.066158271 -0.939466767 -0.953191200
ENSMUSG00000071341 -1.381336772 -1.186527381 -1.132150477 -0.855980942
ENSMUSG00000022602 -1.725511291 -0.343692188 -1.199432055 -0.855967229
ENSMUSG00000021453 -1.089501432 -1.220423084 -1.051068319 -0.923719201
ENSMUSG00000085609 -1.297859422 -0.856126763 -1.086095899 -0.634706053
ENSMUSG00000028195 -1.246375724 -1.225682264 -1.285972745 -0.879241598
ENSMUSG00000114448 -1.044235954 -0.824511100 -1.152717440 -0.688970490
ENSMUSG00000034936 -1.399590664 -0.959792706 -0.976160542 -0.728248850
ENSMUSG00000023034 -1.441511678 -0.839500132 -1.153437909 -0.971664322
ENSMUSG00000021250 -1.243859323 -1.216019321 -1.168200123 -0.715600781
ENSMUSG00000037868 -1.414509435 -1.082336031 -1.316617449 -0.962929208
ENSMUSG00000052837 -1.427612716 -1.153103875 -1.350587537 -0.630127760
ENSMUSG00000087006 -1.333541813 -1.320311778 -1.166371619 -0.531603504
ENSMUSG00000106375 -0.768597135 -1.065161567 -1.137143764 -0.822803846
ENSMUSG00000020423 -1.210988734 -1.372434575 -1.187649620 -0.645723464
ENSMUSG00000032487 -1.009264889 -1.099582826 -1.474146148 -0.784497709
ENSMUSG00000105434 -1.177379828 -0.842104607 -1.286720237 -0.488948622
ENSMUSG00000026360 -1.233042481 -1.223378011 -0.848868632 -0.950677615
ENSMUSG00000003545 -1.267796672 -1.245797471 -1.289260846 -0.663336481
ENSMUSG00000039910 -1.148739914 -1.139917302 -1.163900972 -0.903268227
ENSMUSG00000024190 -1.194665432 -1.144960233 -1.188906659 -0.861365010
ENSMUSG00000043415 -1.262842246 -0.992159083 -1.185492446 -0.916809884
ENSMUSG00000038418 -1.778098715 -0.583359959 -0.993856948 -0.776743543
ENSMUSG00000019960 -1.378858556 -0.837659222 -0.828006156 -1.050327382
ENSMUSG00000053560 -1.218817357 -1.098096568 -1.232403795 -0.541354509
ENSMUSG00000035283 -1.360119695 -0.857396772 -1.121090402 -0.898981077
ENSMUSG00000051495 -0.953960158 -0.459030453 -0.550671124 -1.054526145
ENSMUSG00000038550 -0.641607338 -0.962778279 -1.127249434 -1.187289609
ENSMUSG00000113326 -0.834952166 -0.834743540 -0.666646004 -0.543030033
ENSMUSG00000059991 -1.345322563 -1.389976810 -0.734064481 -1.214124107
ENSMUSG00000032501 -1.326530211 -0.970662696 -1.500243306 -0.628711545
ENSMUSG00000005483 -1.386865837 -0.712114576 -1.193031375 -0.778661297
ENSMUSG00000034765 -1.437969320 -0.702376209 -1.326420401 -0.728791838
ENSMUSG00000004661 -0.876712002  0.384451039 -0.890741722 -0.715302526
ENSMUSG00000029641 -1.442359886 -1.155873095 -0.682105859 -1.449730815
ENSMUSG00000019997 -1.041288693 -1.483433057 -0.400578045 -0.767859573
ENSMUSG00000052684 -1.065845004 -1.562555860 -1.318891427 -0.527626946
ENSMUSG00000070564  1.004755991  0.570956201 -0.408475134 -0.321466618
ENSMUSG00000056749 -1.004100572 -0.464891307 -0.770615665 -1.369002366
ENSMUSG00000044471 -0.912720842 -0.528053386 -0.999543009 -1.052852766
ENSMUSG00000116358 -0.698672290  0.090976987 -0.281286228 -2.059366349
ENSMUSG00000048482 -1.644102502 -0.737049815 -0.717904529 -1.164854803
ENSMUSG00000107272 -2.003499726 -0.381086380 -0.518443310 -0.522899291
ENSMUSG00000044991 -0.490314313 -0.756751344 -0.689505208 -1.836781347
ENSMUSG00000037169 -1.201050799 -0.008809348 -0.577008519 -0.628932731
ENSMUSG00000103220 -0.961389592 -1.086662759 -0.765607427 -0.801381764
ENSMUSG00000022114 -1.461390651 -1.188481973 -1.045748503 -0.541682629
ENSMUSG00000041309 -0.739276975 -0.624032014 -0.020322346 -0.467304812
ENSMUSG00000021647 -0.147102417 -0.279951332  0.221886119 -0.350887548
ENSMUSG00000087528 -0.320821591 -0.472184437 -1.070804437 -0.926617823
ENSMUSG00000052374 -0.150612474 -0.695722462  0.181731244 -0.850353996
ENSMUSG00000019122  0.919699814 -0.114103036  0.583882861  0.730334567
ENSMUSG00000068696  0.308840969 -0.971487752  0.210788515 -0.315464340
ENSMUSG00000118252 -0.455636826 -0.257922411 -0.479383872 -0.931888820
ENSMUSG00000048240  0.261950001 -0.896610681 -0.100742060 -0.979573180
ENSMUSG00000100862 -0.502057395 -0.494632510 -0.275200899  0.767270902
ENSMUSG00000108772 -0.388328046 -0.552437624 -0.239302000  0.347594555
ENSMUSG00000084284 -0.157108259 -0.067394224  0.178791891  0.385728335
ENSMUSG00000082062 -0.576316714 -0.333220237 -0.428035345 -0.287342588
ENSMUSG00000061762 -0.325862570 -1.423571461 -0.271285744 -0.729177284
ENSMUSG00000025905  0.387576286  0.076513453 -0.502333548 -0.156531582
ENSMUSG00000058427  0.031765687  0.248172846 -1.516222912  0.814429376
ENSMUSG00000020491  0.911385385  1.017840222  0.924767059  0.046248694
ENSMUSG00000015981  0.390637717  1.059109517 -0.629686201  0.780794272
ENSMUSG00000082185 -0.031614567 -0.211966344 -0.242807813 -0.147929158
ENSMUSG00000068457  0.949339820  0.855161802  1.025170408 -0.941397528
ENSMUSG00000022840 -0.134798746 -0.663899341 -0.070266575 -0.846436230
ENSMUSG00000021367  1.165952927  0.043484806 -0.219854261 -0.006096291
ENSMUSG00000021478 -0.463930255 -0.689568055  0.362968218 -0.223664282
ENSMUSG00000081227  0.267128276 -0.140157086 -0.365737393  0.972352724
ENSMUSG00000103906  1.107338956  0.006911797  0.852250087  0.683940176
ENSMUSG00000082329 -0.026620760 -0.347868662 -0.587835066  0.090785990
ENSMUSG00000062038 -0.234992502 -0.310800467  0.029444890  0.434228857
ENSMUSG00000101249 -0.535636012 -0.391135228 -0.280938481  0.605576975
ENSMUSG00000034173  0.045644929 -0.059892514 -0.061241790  0.414771224
ENSMUSG00000106755 -0.254615044 -0.195225923 -0.211429021 -0.138914138
ENSMUSG00000041380 -0.829574770 -0.958321810 -0.058133478 -0.772370070
ENSMUSG00000058050 -0.219500851 -0.204051014 -0.074776051  0.526622972
ENSMUSG00000024401  0.945402025  0.647582554 -1.168028686  0.865888438
ENSMUSG00000023868  0.339809125 -0.365390714  0.846182836 -0.156232810
ENSMUSG00000112926 -0.495552960 -0.481862209 -0.459153038 -0.303462061
ENSMUSG00000083863 -0.570685682 -0.347000640 -0.429403075  0.449178127
ENSMUSG00000100131 -0.537112491 -0.478416541 -0.151800397  0.388055950
ENSMUSG00000047507 -0.445221767 -0.891851434 -0.303648735 -0.192851995
ENSMUSG00000100033 -0.673343566 -0.376071416 -0.089133671 -0.226212352
ENSMUSG00000084289 -0.191118542 -0.279252370 -0.358980866  0.389391507
ENSMUSG00000113275 -0.682835072 -0.446360004 -0.177690343 -0.267383946
ENSMUSG00000045573  0.330960752 -0.667554013  0.210850149 -0.774955922
ENSMUSG00000020599  0.284386026 -0.837812492  0.253387112 -0.650863128
ENSMUSG00000024206  1.510450774  0.226356024  0.510215699  0.265119564
ENSMUSG00000069045  0.836147269  0.900609931  0.698905711 -0.946777703
ENSMUSG00000067736 -0.257447660 -0.262201070 -0.134425663  0.589294186
ENSMUSG00000015843 -0.380756494 -0.801703020  0.026200729 -0.839964159
ENSMUSG00000071234  0.280587568 -0.955673936  0.308530105 -0.797481299
ENSMUSG00000090291  0.123583145 -0.836486313  0.139650252 -0.673703659
ENSMUSG00000020178  0.009933572 -0.985975878  0.001295962 -0.636008398
ENSMUSG00000061808 -0.404443807 -0.464356129 -0.450883777 -0.179321860
                       ACSF-5F     ACSF-6F      4AP-1M     4AP-2M       4AP-3M
ENSMUSG00000045903 -1.30015881 -1.22779163  0.82414398  0.9889899  1.035529905
ENSMUSG00000071341 -1.04425093 -0.91440205  0.70350055  1.0096082  1.056459578
ENSMUSG00000022602 -1.21681109 -0.97559815  0.64239212  0.8929890  0.748113352
ENSMUSG00000021453 -1.16789809 -1.07907785  0.69103662  1.0981981  1.078455382
ENSMUSG00000085609 -1.49046277 -0.84930565  0.56710908  1.3588936  1.084184372
ENSMUSG00000028195 -0.78466298 -1.05287342  0.96222373  1.0531017  0.919675023
ENSMUSG00000114448 -1.05858398 -1.10582665  0.69940172  1.1126659  1.633816430
ENSMUSG00000034936 -1.42372993 -0.89963761  0.64509653  1.0962820  1.051917324
ENSMUSG00000023034 -0.94314521 -1.18625261  0.91880350  1.0429161  1.012924643
ENSMUSG00000021250 -1.11186879 -1.06443474  0.87156426  1.0672146  1.105245993
ENSMUSG00000037868 -0.98065884 -0.72628319  0.97273271  1.0382877  0.947721255
ENSMUSG00000052837 -0.87516008 -0.98400424  0.76101208  0.9770188  1.053246268
ENSMUSG00000087006 -1.08862378 -0.92501533  0.73093348  1.1951069  1.167021225
ENSMUSG00000106375 -1.51105226 -1.04141030  1.17603847  0.9460585  1.056694633
ENSMUSG00000020423 -1.00455583 -0.99231146  0.86646663  1.1905241  1.091205531
ENSMUSG00000032487 -0.90188950 -1.02137753  0.85052207  1.2092041  1.183489708
ENSMUSG00000105434 -0.92345883 -1.56152351  0.71284297  0.9145496  1.063613105
ENSMUSG00000026360 -1.09090683 -1.08383282  0.79434035  1.1612193  1.174750210
ENSMUSG00000003545 -0.92177076 -0.98919747  0.88045613  1.1809314  1.144473541
ENSMUSG00000039910 -0.78662875 -1.33706497  0.65560612  0.9274519  1.003802564
ENSMUSG00000024190 -1.02363726 -1.13041241  0.88434567  0.9416628  1.003920324
ENSMUSG00000043415 -0.68727032 -1.21401416  0.87583128  1.1323067  1.068618915
ENSMUSG00000038418 -1.27944643 -0.94131790  0.83756891  1.0436455  0.830220918
ENSMUSG00000019960 -1.13512691 -1.19121134  1.09979838  0.9436034  1.023052774
ENSMUSG00000053560 -1.18135333 -1.18526872  0.91522898  1.0324934  0.942528608
ENSMUSG00000035283 -0.75922789 -1.44996901  1.01783711  0.5672572  0.941948720
ENSMUSG00000051495 -1.34262511 -0.59937488 -0.18314296 -0.7604416 -0.060228431
ENSMUSG00000038550 -1.18904401 -0.85445354  0.91738905  0.8365336  1.514231901
ENSMUSG00000113326 -1.46329252 -0.99892692  1.42896412  0.3487038  1.060904645
ENSMUSG00000059991 -0.87901696 -0.75739996  0.92332297  0.9108709  1.247190097
ENSMUSG00000032501 -0.95062215 -0.90659910  0.71203665  1.1290368  0.969581641
ENSMUSG00000005483 -1.17906211 -0.96512799  0.22344012  1.3314432  0.898277499
ENSMUSG00000034765 -1.19914174 -0.79859065  0.79558137  1.3919847  0.884920219
ENSMUSG00000004661 -2.24894062 -1.03659069  1.25471027  0.7321736  0.710151053
ENSMUSG00000029641 -0.74587396 -0.90666471  0.58885748  1.2404107  0.539371109
ENSMUSG00000019997 -1.29882555 -0.17315676  1.71112123  1.2201174  0.711177771
ENSMUSG00000052684 -0.83291190 -0.96262118  0.79249806  1.2280124  1.197938465
ENSMUSG00000070564 -1.38950753 -1.43285676 -0.35808576 -1.7566442 -0.048060208
ENSMUSG00000056749 -1.42116727 -0.95531225  1.06290287  1.5637820  1.277248956
ENSMUSG00000044471 -0.82611629 -1.19603212  1.45845310  1.0644974  1.180591057
ENSMUSG00000116358 -1.90287398 -0.33579889  0.28516235  0.1780213  0.975934495
ENSMUSG00000048482 -0.96435930 -1.11432701  1.29237578  0.7923996  1.058293880
ENSMUSG00000107272 -1.50043532 -0.34185457  1.15558496  0.4915216  0.458078683
ENSMUSG00000044991 -1.28322841 -0.46242513 -0.33167275  0.3396862  0.913943328
ENSMUSG00000037169 -1.50237932 -1.44561903  0.39424794  0.1060298  0.004441632
ENSMUSG00000103220 -1.15524061 -0.83491277  1.03869162  0.8412602  2.178461191
ENSMUSG00000022114 -1.25645750 -0.69225893  0.77899068  0.4433737  1.291968057
ENSMUSG00000041309 -1.19461790 -0.87950818  2.05099691 -0.9085853  0.075440166
ENSMUSG00000021647 -1.18347072 -2.22406468  0.46777915 -0.7507198  1.536735487
ENSMUSG00000087528 -2.22835378  0.22239794  0.39492877  1.4529136  1.078681067
ENSMUSG00000052374  0.57931473  2.55414291 -0.89424898  0.9267237  1.214348876
ENSMUSG00000019122  1.53062284  1.40752968 -0.79976787 -0.4498754 -0.466535966
ENSMUSG00000068696 -0.61483850  2.76736661  0.05827276  0.3843475  1.126824627
ENSMUSG00000118252  3.10318942 -0.04037663  0.37146052  0.2496080 -0.024669203
ENSMUSG00000048240  0.28574997  2.63451368 -0.11577972  1.0614121  0.934438669
ENSMUSG00000100862  3.09537396 -0.03574554 -0.34243322 -0.8437891 -0.633062881
ENSMUSG00000108772  3.25454806 -0.81899642 -0.65738451 -0.1154423 -0.495128588
ENSMUSG00000084284  3.18632875 -0.94717897 -0.59084981 -0.4459482  0.125570843
ENSMUSG00000082062  3.40649854  0.02159184 -0.45034399 -0.2114187 -0.169057250
ENSMUSG00000061762  0.25350869  2.24257902 -0.18984432  1.3437308  1.290310872
ENSMUSG00000025905  1.50337671  1.90089106  0.55921959  0.2317963  0.914665237
ENSMUSG00000058427  1.06124305  1.08891685 -0.95794295  1.2665773  1.034759279
ENSMUSG00000020491  0.28873196  1.96821103  0.19371281 -0.3874061 -0.243403914
ENSMUSG00000015981 -0.35601725  1.36369355  0.62040573 -0.8811110 -0.509383622
ENSMUSG00000082185  3.42144543 -0.21948867 -0.63085430 -0.2637276 -0.168814406
ENSMUSG00000068457 -1.27963329 -1.09048286  0.86918690  0.8548350  0.735358852
ENSMUSG00000022840  0.16450227  2.81390915 -0.07050058  0.7296144  1.174129173
ENSMUSG00000021367  1.22613997  1.42446361 -0.78421636  0.7962174  0.115156039
ENSMUSG00000021478  0.17275911  2.56581605 -0.02696606  1.0425035  0.939779124
ENSMUSG00000081227  3.03780071 -0.21178389 -0.92051502  0.1555880 -0.089703996
ENSMUSG00000103906  1.96461763  0.99723843 -0.56377344 -0.8098248 -0.701179689
ENSMUSG00000082329  3.24935179  0.18961787 -0.22427917  0.4485669 -0.357687205
ENSMUSG00000062038  3.33411195 -0.38670409 -0.56611927 -0.6136810 -0.279014094
ENSMUSG00000101249  3.28027036 -0.43425553 -0.51003557 -0.6074394 -0.284786798
ENSMUSG00000034173  3.23944829 -0.12025855 -0.58846129 -0.4528622 -0.356425987
ENSMUSG00000106755  3.40035337 -0.57045047 -0.47152088 -0.1614624 -0.403833196
ENSMUSG00000041380  1.51369573  2.14432802 -0.81180091  1.3170512  0.275868441
ENSMUSG00000058050  3.30538693 -0.29055177 -0.38091236 -0.2002623  0.091556269
ENSMUSG00000024401  1.20901031  0.27615251 -0.60710770  1.0114323  0.393921759
ENSMUSG00000023868 -0.36441293  2.66182048 -0.46610185  0.4639652  0.872996257
ENSMUSG00000112926  3.38297302 -0.11693997 -0.23803874  0.2094454  0.032300581
ENSMUSG00000083863  3.32314927  0.01378320 -0.44609261 -0.5114873 -0.448744705
ENSMUSG00000100131  3.29660120  0.02889268 -0.38018793 -0.5372468 -0.439660002
ENSMUSG00000047507  1.86157781  1.46936632 -0.60150381  1.5820724  0.853458257
ENSMUSG00000100033  3.33946524  0.39499147 -0.25515690 -0.4589630 -0.167337351
ENSMUSG00000084289  3.38213981 -0.62109732 -0.33569910 -0.1888734 -0.258041249
ENSMUSG00000113275  3.39321960  0.02449259 -0.21154906  0.1262308 -0.252136456
ENSMUSG00000045573 -0.49314977  2.46373462 -0.15578079  0.8219327  1.554880457
ENSMUSG00000020599 -0.06290074  2.46494177 -0.20906993  0.9431849  1.435485321
ENSMUSG00000024206  1.70842505  1.40098864 -0.53749994 -0.2091506 -0.010409673
ENSMUSG00000069045 -1.28545940 -1.09581642  0.93927235  0.6786907  0.823221999
ENSMUSG00000067736  3.30940195 -0.07130901 -0.35418521 -0.2409037 -0.373997763
ENSMUSG00000015843  0.50430849  2.70403248 -0.69481195  1.0681667  0.759197050
ENSMUSG00000071234  0.03106902  2.64649530 -0.30075444  0.8244767  1.121901676
ENSMUSG00000090291 -0.08348796  2.75200685 -0.19200877  0.6777253  1.260274114
ENSMUSG00000020178 -0.04364984  2.58280211 -0.10168076  0.9250542  1.426464602
ENSMUSG00000061808 -0.48548885 -0.34655571  2.02586283 -0.4260944  2.647275280
                         4AP-4M       4AP-5M      4AP-6F      4AP-7F
ENSMUSG00000045903  0.761034472  0.385232486  0.83199112  0.88456927
ENSMUSG00000071341  0.556892317  0.371011182  0.83254743  0.95779326
ENSMUSG00000022602  0.828517388  0.367544075  0.82513982  0.96208993
ENSMUSG00000021453  0.822887118  0.250265424  0.87043530  0.94138919
ENSMUSG00000085609  0.310802234  0.239971743  0.98690505  1.14687157
ENSMUSG00000028195  0.657398466  0.181021585  0.91408464  0.97967871
ENSMUSG00000114448  0.781800562 -0.369351642  1.20024272  0.69532340
ENSMUSG00000034936  0.865857971  0.140831983  0.67237703  0.85497715
ENSMUSG00000023034  0.762892103  0.365352448  0.81417097  0.80330983
ENSMUSG00000021250  0.582218391  0.364668810  0.88774291  0.84179970
ENSMUSG00000037868  0.372298221  0.503107443  0.80534588  1.00913565
ENSMUSG00000052837  0.546365402  0.255682029  0.88312048  0.98372256
ENSMUSG00000087006  0.442746230  0.269520234  0.91681644  0.89482804
ENSMUSG00000106375  0.526369758  0.086601818  0.78266546  0.69168696
ENSMUSG00000020423  0.446070404  0.333689907  1.05319967  0.82626986
ENSMUSG00000032487  0.189075566  0.154037697  0.75615312  0.89733833
ENSMUSG00000105434  0.347515830  0.202376594  0.97637578  1.02062356
ENSMUSG00000026360  0.673742161  0.091811069  0.88974802  0.93914915
ENSMUSG00000003545  0.461870748  0.144554975  0.89182905  0.80860076
ENSMUSG00000039910  0.688123790  0.282162404  0.70951385  1.13268975
ENSMUSG00000024190  0.560237272  0.318330644  0.86833955  0.99056158
ENSMUSG00000043415  0.185498952  0.018180712  0.96573269  1.15671861
ENSMUSG00000038418  0.773398918  0.296814237  0.92964825  0.76539401
ENSMUSG00000019960  0.616488866  0.076303828  0.82254172  0.91409003
ENSMUSG00000053560  0.450776856  0.359742167  0.76730975  1.04158685
ENSMUSG00000035283  0.556003685  0.451573955  0.73190804  1.07354586
ENSMUSG00000051495  0.913842239  1.178509772  0.75080304  1.68108132
ENSMUSG00000038550  0.707495171 -0.285382300  0.07929544  1.02484789
ENSMUSG00000113326  1.295257708 -0.936485865  1.15863689  0.65980342
ENSMUSG00000059991  0.098275599  0.444782372  0.79052380  1.03699760
ENSMUSG00000032501  0.306164594  0.125273665  0.98506316  0.95419573
ENSMUSG00000005483  0.656815642  0.045707068  1.17371913  0.93842680
ENSMUSG00000034765  0.298283969  0.067479728  0.82395802  0.76297306
ENSMUSG00000004661  0.982257055 -0.018258933  0.46906089  0.45304007
ENSMUSG00000029641  0.853726383  0.777480230  1.13140543  0.44463159
ENSMUSG00000019997  0.545989303 -0.813632078  0.15680175  0.87504221
ENSMUSG00000052684  0.349538078  0.251690732  0.91892799  0.83589520
ENSMUSG00000070564  0.962198184  1.344166973  0.29911772  0.45419659
ENSMUSG00000056749  0.552589164  0.096955234  0.77999002  0.45488072
ENSMUSG00000044471  0.633199203 -0.996403069  0.56304689  0.84822399
ENSMUSG00000116358  0.561618498  0.405588161  1.35001115  0.88628720
ENSMUSG00000048482  0.504520085  0.269514390  0.85468511  0.79523692
ENSMUSG00000107272  1.461855484 -0.239175669  0.35746315  0.25815325
ENSMUSG00000044991  1.650900850  0.427252520  0.33483570  1.01671098
ENSMUSG00000037169  0.862965194  1.731675128  0.10335069  0.74438245
ENSMUSG00000103220  0.775494240 -0.309704646  0.43866381  0.37837929
ENSMUSG00000022114  0.004200030  0.850213889  1.05748191  1.14893018
ENSMUSG00000041309 -0.098292000  0.377409841 -0.47461104  1.54249448
ENSMUSG00000021647  0.584360733 -0.196922917 -0.06553303  1.21686019
ENSMUSG00000087528 -0.405775384  0.007260860  1.18841994  0.36225351
ENSMUSG00000052374 -0.824343264 -0.430566169 -0.62025264 -0.44551866
ENSMUSG00000019122 -1.785149139  0.378388027 -1.54888597 -0.39265943
ENSMUSG00000068696 -1.177974318 -0.615187788 -0.69402943 -0.19721097
ENSMUSG00000118252  0.663519867 -0.448488544 -0.53691938 -0.86176851
ENSMUSG00000048240 -0.798393058 -0.679346958 -0.32471225 -0.42355305
ENSMUSG00000100862 -0.357602281 -0.286513672 -0.44295541 -0.35267636
ENSMUSG00000108772  0.419907696 -0.120029681 -0.14416982 -0.44601787
ENSMUSG00000084284  0.258346141 -0.139804303 -0.50186064 -0.56118254
ENSMUSG00000082062 -0.072100534 -0.006761904 -0.22848864 -0.58910513
ENSMUSG00000061762  0.101728957 -0.684254393 -0.09276133 -0.87386129
ENSMUSG00000025905 -0.793551849 -1.633022807 -0.94385737 -1.00691755
ENSMUSG00000058427 -1.420399644  0.597161528 -0.81779670 -0.76153811
ENSMUSG00000020491  0.024137140 -0.648441129 -1.55442509 -1.21860091
ENSMUSG00000015981  1.596158044 -0.001988420 -0.74619885 -0.79573116
ENSMUSG00000082185 -0.007343407 -0.377967321 -0.24087046 -0.57377609
ENSMUSG00000068457  0.482051488  0.841297255 -1.13592432 -1.11881730
ENSMUSG00000022840 -0.629074418 -0.590847649 -0.61558103 -0.48873928
ENSMUSG00000021367 -1.383378150  0.588400042 -1.42290205  0.04070889
ENSMUSG00000021478 -0.113703482 -1.291943054 -1.03862538 -0.51875904
ENSMUSG00000081227 -0.138221032 -0.588380340 -0.65141383 -0.72277819
ENSMUSG00000103906 -1.413946315  0.363826658 -0.92754206 -0.61929587
ENSMUSG00000082329 -0.798826678 -0.514380999 -0.56470704 -0.57348368
ENSMUSG00000062038 -0.240077169 -0.150278807 -0.41726079 -0.60301618
ENSMUSG00000101249  0.014675998 -0.130838561 -0.49610609 -0.40253611
ENSMUSG00000034173  0.137154149 -0.045274745 -0.61621006 -0.92760205
ENSMUSG00000106755  0.243019197 -0.443362702 -0.06593291 -0.43327737
ENSMUSG00000041380 -0.065576148 -0.511747687  0.23467139 -0.84076301
ENSMUSG00000058050 -0.313246723 -0.711170674 -0.39213293 -0.46928830
ENSMUSG00000024401 -1.249071499  0.968894115 -1.51640896 -1.12292820
ENSMUSG00000023868 -0.690450566 -0.338199344 -0.42859425 -1.15746996
ENSMUSG00000112926 -0.322180037 -0.673888750 -0.14769496 -0.14123509
ENSMUSG00000083863 -0.196118677 -0.344261085 -0.11597883 -0.52059582
ENSMUSG00000100131 -0.462315388 -0.207980533 -0.54167943 -0.35150473
ENSMUSG00000047507 -0.891101459 -0.518068008 -0.17957452 -0.71933839
ENSMUSG00000100033 -0.350342256  0.115552679 -0.63801605 -0.33709561
ENSMUSG00000084289 -0.098319893 -0.424298871 -0.46527684 -0.21335605
ENSMUSG00000113275 -0.275540699 -0.193816144 -0.06195124 -0.47497242
ENSMUSG00000045573 -0.302243845 -0.782598164 -0.79587708 -0.56802150
ENSMUSG00000020599 -0.761475618 -0.611060986 -0.85202823 -0.72726640
ENSMUSG00000024206 -1.078387811 -1.045566177 -1.26319450 -0.43918836
ENSMUSG00000069045  0.846999475  0.919628676 -1.13861361 -1.12246344
ENSMUSG00000067736 -0.370486094 -0.455595077 -0.42829347 -0.82110492
ENSMUSG00000015843 -0.039009648 -0.355564681 -0.22879946 -0.66303623
ENSMUSG00000071234 -0.379309061 -0.904890801 -0.79274849 -0.35906290
ENSMUSG00000090291 -0.700280830 -0.761822942 -0.65950678 -0.30624221
ENSMUSG00000020178 -0.735858351 -0.564691492 -0.68149360 -0.52964297
ENSMUSG00000061808 -0.406149689 -0.395915323 -0.38434618 -0.37549238
                         4AP-8F
ENSMUSG00000045903  0.866696879
ENSMUSG00000071341  1.026836008
ENSMUSG00000022602  1.050226277
ENSMUSG00000021453  0.779020802
ENSMUSG00000085609  0.519818869
ENSMUSG00000028195  0.807624890
ENSMUSG00000114448  0.120946561
ENSMUSG00000034936  1.059820336
ENSMUSG00000023034  0.815142276
ENSMUSG00000021250  0.799528391
ENSMUSG00000037868  0.834705257
ENSMUSG00000052837  0.960428564
ENSMUSG00000087006  0.748495311
ENSMUSG00000106375  1.080053273
ENSMUSG00000020423  0.606237597
ENSMUSG00000032487  1.050938002
ENSMUSG00000105434  1.042238161
ENSMUSG00000026360  0.705946115
ENSMUSG00000003545  0.864443133
ENSMUSG00000039910  1.080169732
ENSMUSG00000024190  0.976549183
ENSMUSG00000043415  0.855700252
ENSMUSG00000038418  0.876132755
ENSMUSG00000019960  0.925310621
ENSMUSG00000053560  0.947627651
ENSMUSG00000035283  1.106710320
ENSMUSG00000051495  1.439764495
ENSMUSG00000038550  1.168011470
ENSMUSG00000113326  0.325806491
ENSMUSG00000059991  0.867941547
ENSMUSG00000032501  1.102016741
ENSMUSG00000005483  0.947033707
ENSMUSG00000034765  1.168109043
ENSMUSG00000004661  0.800702492
ENSMUSG00000029641  0.806725391
ENSMUSG00000019997  0.758524138
ENSMUSG00000052684  0.695951416
ENSMUSG00000070564  1.079704541
ENSMUSG00000056749  0.196740517
ENSMUSG00000044471  0.763709878
ENSMUSG00000116358  0.544397628
ENSMUSG00000048482  0.775572151
ENSMUSG00000107272  1.324737173
ENSMUSG00000044991  1.167348939
ENSMUSG00000037169  1.416706883
ENSMUSG00000103220  0.263949233
ENSMUSG00000022114  0.610861747
ENSMUSG00000041309  1.360209186
ENSMUSG00000021647  1.171030772
ENSMUSG00000087528  0.717701820
ENSMUSG00000052374 -0.544642862
ENSMUSG00000019122  0.006518969
ENSMUSG00000068696 -0.270247889
ENSMUSG00000118252 -0.350723581
ENSMUSG00000048240 -0.859353454
ENSMUSG00000100862  0.704024440
ENSMUSG00000108772 -0.044813428
ENSMUSG00000084284 -0.723438985
ENSMUSG00000082062 -0.075899360
ENSMUSG00000061762 -0.641239915
ENSMUSG00000025905 -0.537823899
ENSMUSG00000058427 -0.669125607
ENSMUSG00000020491 -1.322757139
ENSMUSG00000015981 -1.890682307
ENSMUSG00000082185 -0.304285300
ENSMUSG00000068457 -1.046146250
ENSMUSG00000022840 -0.772011100
ENSMUSG00000021367 -1.584076543
ENSMUSG00000021478 -0.716666405
ENSMUSG00000081227 -0.604178906
ENSMUSG00000103906 -0.940561565
ENSMUSG00000082329  0.017366682
ENSMUSG00000062038  0.004158653
ENSMUSG00000101249  0.173184421
ENSMUSG00000034173 -0.608789413
ENSMUSG00000106755 -0.293348528
ENSMUSG00000041380 -0.637326890
ENSMUSG00000058050 -0.667673184
ENSMUSG00000024401 -0.654738996
ENSMUSG00000023868 -1.217921450
ENSMUSG00000112926 -0.244711216
ENSMUSG00000083863  0.144257870
ENSMUSG00000100131  0.374354420
ENSMUSG00000047507 -1.023314664
ENSMUSG00000100033 -0.278337202
ENSMUSG00000084289 -0.337216871
ENSMUSG00000113275 -0.499707581
ENSMUSG00000045573 -0.842177554
ENSMUSG00000020599 -0.668907564
ENSMUSG00000024206 -1.038158675
ENSMUSG00000069045 -1.054345553
ENSMUSG00000067736 -0.128746487
ENSMUSG00000015843 -1.058259768
ENSMUSG00000071234 -0.723139407
ENSMUSG00000090291 -0.739700193
ENSMUSG00000020178 -0.666549161
ENSMUSG00000061808 -0.354090045
```

```
#Selecting gene list
num_keep <- 30
num <- 10
rows_keep <- c(seq(1:num_keep), seq((nrow(scaled_mat)- num), nrow(scaled_mat) ) )
rows_keep
```

```
 [1]  1  2  3  4  5  6  7  8  9 10 11 12 13 14 15 16 17 18 19 20 21 22 23 24 25
[26] 26 27 28 29 30 87 88 89 90 91 92 93 94 95 96 97
```

```
#Getting l2 fold changes for map
l2_val <- as.matrix(top.treatment[rows_keep,] $log2FoldChange)
#rownames(l2_val) <- rownames(top.treatment1[rows_keep,])
l2_val
```

```
           [,1]
 [1,]  4.145262
 [2,]  2.673402
 [3,]  2.445002
 [4,]  2.424258
 [5,]  2.071698
 [6,]  2.021751
 [7,]  2.013174
 [8,]  2.009553
 [9,]  1.914216
[10,]  1.755309
[11,]  1.717922
[12,]  1.671464
[13,]  1.616261
[14,]  1.586009
[15,]  1.524861
[16,]  1.490393
[17,]  1.475102
[18,]  1.408863
[19,]  1.360141
[20,]  1.308182
[21,]  1.284420
[22,]  1.275364
[23,]  1.255876
[24,]  1.227866
[25,]  1.153399
[26,]  1.107827
[27,]  1.080424
[28,]  1.066785
[29,]  1.042042
[30,]  1.030389
[31,] -1.462734
[32,] -1.512748
[33,] -1.623274
[34,] -1.715373
[35,] -1.751013
[36,] -1.756945
[37,] -1.779825
[38,] -2.179828
[39,] -2.288731
[40,] -2.606568
[41,] -2.808529
```

```
colnames(l2_val) <- "logFC"
#Same as above but means 
means <- as.matrix(top.treatment[rows_keep,] $baseMean)
colnames(means) <- "AveExpr"

library(ComplexHeatmap)
```

```
Loading required package: grid
```

```
Attaching package: 'grid'
```

```
The following object is masked from 'package:Biostrings':

    pattern
```

```
========================================
ComplexHeatmap version 2.16.0
Bioconductor page: http://bioconductor.org/packages/ComplexHeatmap/
Github page: https://github.com/jokergoo/ComplexHeatmap
Documentation: http://jokergoo.github.io/ComplexHeatmap-reference

If you use it in published research, please cite either one:
- Gu, Z. Complex Heatmap Visualization. iMeta 2022.
- Gu, Z. Complex heatmaps reveal patterns and correlations in multidimensional 
    genomic data. Bioinformatics 2016.


The new InteractiveComplexHeatmap package can directly export static 
complex heatmaps into an interactive Shiny app with zero effort. Have a try!

This message can be suppressed by:
  suppressPackageStartupMessages(library(ComplexHeatmap))
========================================
```

```
library(RColorBrewer)
library(circlize)
```

```
========================================
circlize version 0.4.15
CRAN page: https://cran.r-project.org/package=circlize
Github page: https://github.com/jokergoo/circlize
Documentation: https://jokergoo.github.io/circlize_book/book/

If you use it in published research, please cite:
Gu, Z. circlize implements and enhances circular visualization
  in R. Bioinformatics 2014.

This message can be suppressed by:
  suppressPackageStartupMessages(library(circlize))
========================================
```

```
#color selection for fold changes and means
col_logFC <- colorRamp2(c(min(l2_val), 0, max(l2_val)), c("blue", "white", "red"))
col_averages <- colorRamp2(c(quantile(means)[1], quantile(means)[4]), c("white", "red"))

#Building Heatmap
ha <- HeatmapAnnotation(summary= anno_summary(gp= gpar(fill= 2), height= unit(2, 'cm')))

h1 <- Heatmap(scaled_mat[rows_keep,], cluster_rows = F, 
              row_names_gp = gpar(fontsize = 24),
              column_labels= colnames(scaled_mat), 
              column_names_gp = gpar(fontsize= 24),
              name = "Z-score", 
              cluster_columns= T,
              heatmap_legend_param= list(title_gp = gpar(fontsize = 8), 
              labels_gp = gpar(fontsize = 14)))

#h2 <- Heatmap(l2_val, cluster_rows = F, row_names_gp = rownames(l2_val))


h <- h1
h

png("./treatmentheatmap3.png", res= 300, width = 5000, height = 5500)
print(h)
```

GO ANALYSIS

```
library(clusterProfiler)
```

```
clusterProfiler v4.8.1  For help: https://yulab-smu.top/biomedical-knowledge-mining-book/

If you use clusterProfiler in published research, please cite:
T Wu, E Hu, S Xu, M Chen, P Guo, Z Dai, T Feng, L Zhou, W Tang, L Zhan, X Fu, S Liu, X Bo, and G Yu. clusterProfiler 4.0: A universal enrichment tool for interpreting omics data. The Innovation. 2021, 2(3):100141
```

```
Attaching package: 'clusterProfiler'
```

```
The following object is masked from 'package:AnnotationDbi':

    select
```

```
The following object is masked from 'package:XVector':

    slice
```

```
The following object is masked from 'package:IRanges':

    slice
```

```
The following object is masked from 'package:S4Vectors':

    rename
```

```
The following object is masked from 'package:stats':

    filter
```

```
library(AnnotationDbi)
library(org.Mm.eg.db)
library(enrichplot)


treatment4 <- treatment[abs(treatment$log2FoldChange) > .58 & treatment$padj< 0.05,]
treatment4
```

```
                       baseMean log2FoldChange      lfcSE      stat
ENSMUSG00000045903  6665.662737      4.1452625 0.27976542 14.816922
ENSMUSG00000114708     9.122834      3.5631858 0.86691178  4.110206
ENSMUSG00000091123    16.252922     -3.2008390 0.73631076 -4.347131
ENSMUSG00000114218     9.925000      3.0372850 0.84850237  3.579583
ENSMUSG00000094955    11.083658     -3.0365634 0.80952198 -3.751057
ENSMUSG00002075338    18.969287      2.8456441 0.53752909  5.293935
ENSMUSG00000090941     9.683991      2.7333472 0.75969229  3.597966
ENSMUSG00000071341  4230.195185      2.6734022 0.22357906 11.957301
ENSMUSG00000117518    15.301298     -2.6337621 0.67429304 -3.905961
ENSMUSG00000065537     9.971100      2.6214524 0.74944850  3.497842
ENSMUSG00000113076    62.297914      2.5689201 0.33376096  7.696886
ENSMUSG00000022602  8172.399871      2.4450017 0.37462886  6.526464
ENSMUSG00000021453  1860.279738      2.4242584 0.21774154 11.133651
ENSMUSG00000113183    22.250091      2.3910393 0.53374676  4.479726
ENSMUSG00000105528    16.877918      2.1069994 0.49432255  4.262398
ENSMUSG00000085609    97.527042      2.0716977 0.28490376  7.271570
ENSMUSG00000083270    21.397472     -2.0299973 0.55407700 -3.663746
ENSMUSG00000028195  2964.327673      2.0217506 0.20215450 10.001017
ENSMUSG00000114448    93.785310      2.0131742 0.33406590  6.026279
ENSMUSG00000034936   442.220180      2.0095535 0.27279271  7.366595
ENSMUSG00000086894    11.335571     -1.9342583 0.54789001 -3.530377
ENSMUSG00000081758    25.339692     -1.9315014 0.52381562 -3.687369
ENSMUSG00000108761    38.934995     -1.9216194 0.45991074 -4.178244
ENSMUSG00000023034  4266.695513      1.9142160 0.18582816 10.301001
ENSMUSG00000031103    14.631826     -1.8099771 0.45639596 -3.965804
ENSMUSG00000067736    89.454888     -1.7569453 0.47838263 -3.672678
ENSMUSG00000021250 10963.375922      1.7553094 0.14001130 12.536913
ENSMUSG00000037868   420.633278      1.7179216 0.15536860 11.057071
ENSMUSG00000024206    85.442389     -1.7153728 0.31743568 -5.403844
ENSMUSG00000052837  6077.207306      1.6714639 0.14080789 11.870527
ENSMUSG00000087006  1142.592593      1.6162607 0.22040682  7.333079
ENSMUSG00000106375   156.428926      1.5860091 0.19486107  8.139179
ENSMUSG00000020423  1189.847188      1.5248610 0.15661888  9.736125
ENSMUSG00000080904    45.651450     -1.5172686 0.40562009 -3.740615
ENSMUSG00000032487   963.803496      1.4903930 0.22878702  6.514325
ENSMUSG00000105434    78.206821      1.4751017 0.23432441  6.295126
ENSMUSG00000097028    25.151703      1.4638636 0.37257642  3.929029
ENSMUSG00000121250    51.614589      1.4099226 0.26191055  5.383222
ENSMUSG00000026360  1118.310912      1.4088627 0.14628248  9.631110
ENSMUSG00000003545   998.080227      1.3601408 0.15021483  9.054637
ENSMUSG00000104585    27.183147      1.3430160 0.36628460  3.666592
ENSMUSG00000045991    29.218375      1.3412769 0.36691152  3.655587
ENSMUSG00000039910   744.630721      1.3081818 0.12238296 10.689248
ENSMUSG00000024190  4509.976244      1.2844203 0.09674751 13.276003
ENSMUSG00000043415   859.551765      1.2753643 0.15638946  8.155053
ENSMUSG00000051499    55.948072     -1.2653101 0.25099067 -5.041263
ENSMUSG00000038418 10218.949166      1.2558763 0.18482415  6.794980
ENSMUSG00000019960  2461.587729      1.2278664 0.12496207  9.825912
ENSMUSG00000028004    40.641618     -1.1995500 0.32599844 -3.679619
ENSMUSG00000053560   949.511918      1.1533986 0.11843914  9.738323
ENSMUSG00000028702    30.578605     -1.1533544 0.32455306 -3.553670
ENSMUSG00000107128    32.825489     -1.1406396 0.31863012 -3.579824
ENSMUSG00000046761    34.107682     -1.1322743 0.31116037 -3.638877
ENSMUSG00000035283   627.213251      1.1078268 0.12060489  9.185588
ENSMUSG00000051495   384.621061      1.0804237 0.18378754  5.878656
ENSMUSG00000105733    49.684498     -1.0737449 0.29385704 -3.653970
ENSMUSG00000038550   452.941774      1.0667854 0.18173579  5.869980
ENSMUSG00000113326    76.409513      1.0420423 0.27065940  3.850013
ENSMUSG00000059991  1341.711480      1.0303893 0.13498491  7.633367
ENSMUSG00000021765    72.858677      1.0264997 0.24130189  4.254006
ENSMUSG00000032501   874.384402      1.0168269 0.12404084  8.197517
ENSMUSG00000005483  1146.368693      1.0140892 0.13316858  7.615079
ENSMUSG00000103906   141.401587     -1.0055735 0.20551987 -4.892828
ENSMUSG00000081227    99.688343     -0.9981897 0.25233596 -3.955796
ENSMUSG00000034765   400.447282      0.9814953 0.15379408  6.381880
ENSMUSG00000004661   133.387184      0.9813079 0.21253837  4.617086
ENSMUSG00000002083    60.700122     -0.9318011 0.23942674 -3.891801
ENSMUSG00000029641   111.989475      0.9159481 0.18361289  4.988474
ENSMUSG00000020491    87.092689     -0.8967023 0.20542364 -4.365137
ENSMUSG00000052684  4526.330284      0.8903146 0.12521982  7.110013
ENSMUSG00000025905   138.700546     -0.8781166 0.25097345 -3.498843
ENSMUSG00000056749   342.903795      0.8642388 0.15174224  5.695440
ENSMUSG00000044471    99.505750      0.8392426 0.22579348  3.716859
ENSMUSG00000116358    94.981151      0.8391183 0.19527775  4.297050
ENSMUSG00000048482   531.607150      0.8337126 0.12444504  6.699445
ENSMUSG00000056724    73.459280     -0.8284175 0.22446073 -3.690701
ENSMUSG00000044991    76.471606      0.8271356 0.22385471  3.694966
ENSMUSG00000037169   105.155215      0.8245357 0.20905147  3.944176
ENSMUSG00000022114   132.752677      0.8117090 0.16769720  4.840325
ENSMUSG00000021647   118.227508      0.8061487 0.22150644  3.639392
ENSMUSG00000038530  9266.200868      0.7946938 0.08308958  9.564301
ENSMUSG00000035828  1468.046737      0.7893853 0.12153705  6.495018
ENSMUSG00000029135   367.449176      0.7871599 0.14646887  5.374247
ENSMUSG00000026628  1048.176732      0.7832334 0.13026635  6.012553
ENSMUSG00000051243   187.373763     -0.7829910 0.20288089 -3.859363
ENSMUSG00000037573   538.330581      0.7583254 0.11328097  6.694200
ENSMUSG00000031530   114.357147      0.7417933 0.21093404  3.516707
ENSMUSG00000072620    98.337931     -0.7341606 0.19079279 -3.847947
ENSMUSG00000018648  1730.610238      0.7324927 0.08473705  8.644303
ENSMUSG00000034640   776.278952      0.7252324 0.13874976  5.226910
ENSMUSG00000017144   251.734732      0.7210799 0.15982258  4.511752
ENSMUSG00000028341   229.179004      0.7206668 0.18357749  3.925682
ENSMUSG00000019970  2070.238310      0.7203648 0.14140023  5.094509
ENSMUSG00000075224   271.349578      0.7169452 0.16630306  4.311076
ENSMUSG00000049649   137.163745      0.7060638 0.18138908  3.892538
ENSMUSG00000024112   187.136807     -0.6980849 0.19186982 -3.638326
ENSMUSG00000051652   118.419237     -0.6957626 0.19572615 -3.554776
ENSMUSG00000057706   135.752195      0.6893778 0.18862636  3.654727
ENSMUSG00000037211   314.689143      0.6721476 0.14132052  4.756193
ENSMUSG00000050890   139.273772     -0.6468584 0.16864165 -3.835698
ENSMUSG00000035329   959.821608      0.6410477 0.09784123  6.551917
ENSMUSG00000003032   330.405074      0.6376091 0.11922532  5.347934
ENSMUSG00000076431   607.867500      0.6375422 0.09230281  6.907072
ENSMUSG00000024042   277.290730      0.6311412 0.14421280  4.376457
ENSMUSG00000020893   793.008145      0.6211581 0.12062730  5.149399
ENSMUSG00000072893   292.405041      0.6014995 0.13155112  4.572364
ENSMUSG00000038894  2427.114279      0.5915334 0.10012513  5.907941
ENSMUSG00000090877   561.741022      0.5894376 0.14103919  4.179247
ENSMUSG00000026814   236.203481      0.5869095 0.13943546  4.209184
ENSMUSG00000051022   242.965864      0.5809589 0.14270033  4.071181
                         pvalue         padj        symbol
ENSMUSG00000045903 1.138841e-49 2.012788e-45         Npas4
ENSMUSG00000114708 3.953066e-05 7.762943e-03          <NA>
ENSMUSG00000091123 1.379301e-05 3.216254e-03          <NA>
ENSMUSG00000114218 3.441423e-04 4.028061e-02          <NA>
ENSMUSG00000094955 1.760904e-04 2.551002e-02          <NA>
ENSMUSG00002075338 1.197119e-07 3.846888e-05          <NA>
ENSMUSG00000090941 3.207152e-04 3.882411e-02          <NA>
ENSMUSG00000071341 5.946308e-33 2.627376e-29          Egr4
ENSMUSG00000117518 9.385176e-05 1.521776e-02          <NA>
ENSMUSG00000065537 4.690389e-04 4.963948e-02        Mir132
ENSMUSG00000113076 1.394221e-14 1.071368e-11          <NA>
ENSMUSG00000022602 6.734062e-11 3.400509e-08           Arc
ENSMUSG00000021453 8.603731e-29 2.534372e-25       Gadd45g
ENSMUSG00000113183 7.473887e-06 1.860471e-03          <NA>
ENSMUSG00000105528 2.022448e-05 4.468094e-03          <NA>
ENSMUSG00000085609 3.553325e-13 2.242910e-10          <NA>
ENSMUSG00000083270 2.485535e-04 3.278309e-02       Gm13498
ENSMUSG00000028195 1.508395e-23 2.665937e-20          Ccn1
ENSMUSG00000114448 1.677779e-09 7.413265e-07          <NA>
ENSMUSG00000034936 1.750411e-13 1.189876e-10         Arl4d
ENSMUSG00000086894 4.149679e-04 4.527248e-02       Gm15708
ENSMUSG00000081758 2.265848e-04 3.104388e-02          <NA>
ENSMUSG00000108761 2.937683e-05 6.037280e-03          <NA>
ENSMUSG00000023034 6.973217e-25 1.369385e-21         Nr4a1
ENSMUSG00000031103 7.314887e-05 1.292833e-02          Elf4
ENSMUSG00000067736 2.400219e-04 3.238281e-02       Gm10222
ENSMUSG00000021250 4.689102e-36 2.762506e-32           Fos
ENSMUSG00000037868 2.026068e-28 5.115531e-25          Egr2
ENSMUSG00000024206 6.522777e-08 2.352726e-05          Rfx2
ENSMUSG00000052837 1.684025e-32 5.952692e-29          Junb
ENSMUSG00000087006 2.249238e-13 1.472335e-10       Gm13889
ENSMUSG00000106375 3.979681e-16 3.349376e-13          <NA>
ENSMUSG00000020423 2.114640e-22 2.874935e-19          Btg2
ENSMUSG00000080904 1.835706e-04 2.637745e-02          <NA>
ENSMUSG00000032487 7.301724e-11 3.584741e-08         Ptgs2
ENSMUSG00000105434 3.071502e-10 1.391942e-07          <NA>
ENSMUSG00000097028 8.528950e-05 1.435625e-02       Ptgs2os
ENSMUSG00000121250 7.316418e-08 2.531286e-05          <NA>
ENSMUSG00000026360 5.908744e-22 7.459368e-19          Rgs2
ENSMUSG00000003545 1.370229e-19 1.424554e-16          Fosb
ENSMUSG00000104585 2.458049e-04 3.266433e-02 4921511C10Rik
ENSMUSG00000045991 2.565944e-04 3.307038e-02       Onecut2
ENSMUSG00000039910 1.142940e-26 2.525039e-23        Cited2
ENSMUSG00000024190 3.189697e-40 2.818735e-36         Dusp1
ENSMUSG00000043415 3.490272e-16 3.084353e-13         Otud1
ENSMUSG00000051499 4.624685e-07 1.385368e-04        Zfp786
ENSMUSG00000038418 1.083277e-11 6.176076e-09          Egr1
ENSMUSG00000019960 8.708189e-23 1.399169e-19         Dusp6
ENSMUSG00000028004 2.335827e-04 3.175646e-02         Npy2r
ENSMUSG00000053560 2.069417e-22 2.874935e-19          Ier2
ENSMUSG00000028702 3.798960e-04 4.304027e-02        Rad54l
ENSMUSG00000107128 3.438262e-04 4.028061e-02          <NA>
ENSMUSG00000046761 2.738294e-04 3.439741e-02        Fam83h
ENSMUSG00000035283 4.092845e-20 4.521059e-17         Adrb1
ENSMUSG00000051495 4.136113e-09 1.700039e-06       Irf2bp2
ENSMUSG00000105733 2.582161e-04 3.307038e-02          <NA>
ENSMUSG00000038550 4.358484e-09 1.750724e-06         Ciart
ENSMUSG00000113326 1.181115e-04 1.830591e-02          <NA>
ENSMUSG00000059991 2.287014e-14 1.684195e-11         Nptx2
ENSMUSG00000021765 2.099798e-05 4.581706e-03           Fst
ENSMUSG00000032501 2.454025e-16 2.282760e-13         Trib1
ENSMUSG00000005483 2.635307e-14 1.863057e-11        Dnajb1
ENSMUSG00000103906 9.939705e-07 2.744912e-04         Tigd5
ENSMUSG00000081227 7.628008e-05 1.321739e-02          <NA>
ENSMUSG00000034765 1.749273e-10 8.135962e-08         Dusp5
ENSMUSG00000004661 3.891662e-06 1.011489e-03        Arid3b
ENSMUSG00000002083 9.950301e-05 1.570193e-02          Bbc3
ENSMUSG00000029641 6.085816e-07 1.763290e-04       Rasl11a
ENSMUSG00000020491 1.270431e-05 3.034271e-03 2810021J22Rik
ENSMUSG00000052684 1.160319e-12 7.071545e-10           Jun
ENSMUSG00000025905 4.672824e-04 4.963948e-02         Oprk1
ENSMUSG00000056749 1.230543e-08 4.833027e-06         Nfil3
ENSMUSG00000044471 2.017147e-04 2.829449e-02       Lncpint
ENSMUSG00000116358 1.730859e-05 3.872304e-03          <NA>
ENSMUSG00000048482 2.092133e-11 1.155511e-08          Bdnf
ENSMUSG00000056724 2.236365e-04 3.087930e-02        Nbeal2
ENSMUSG00000044991 2.199160e-04 3.060469e-02         Shld1
ENSMUSG00000037169 8.007498e-05 1.374024e-02          Mycn
ENSMUSG00000022114 1.296269e-06 3.524655e-04         Spry2
ENSMUSG00000021647 2.732827e-04 3.439741e-02        Cartpt
ENSMUSG00000038530 1.129623e-21 1.330997e-18          Rgs4
ENSMUSG00000035828 8.302355e-11 3.965833e-08          Pim3
ENSMUSG00000029135 7.690338e-08 2.564510e-05         Fosl2
ENSMUSG00000026628 1.826241e-09 7.872435e-07          Atf3
ENSMUSG00000051243 1.136829e-04 1.778081e-02         Islr2
ENSMUSG00000037573 2.168554e-11 1.161425e-08          Tob1
ENSMUSG00000031530 4.369351e-04 4.711956e-02         Dusp4
ENSMUSG00000072620 1.191117e-04 1.830591e-02         Slfn2
ENSMUSG00000018648 5.413493e-18 5.315448e-15        Dusp14
ENSMUSG00000034640 1.723666e-07 5.440014e-05        Tiparp
ENSMUSG00000017144 6.429430e-06 1.623339e-03          Rnd3
ENSMUSG00000028341 8.648439e-05 1.442005e-02         Nr4a3
ENSMUSG00000019970 3.496455e-07 1.065454e-04          Sgk1
ENSMUSG00000075224 1.624619e-05 3.681220e-03        Lrrc55
ENSMUSG00000049649 9.920111e-05 1.570193e-02          Gpr3
ENSMUSG00000024112 2.744164e-04 3.439741e-02       Cacna1h
ENSMUSG00000051652 3.783018e-04 4.304027e-02         Lrrc3
ENSMUSG00000057706 2.574560e-04 3.307038e-02         Mex3b
ENSMUSG00000037211 1.972780e-06 5.282866e-04         Spry1
ENSMUSG00000050890 1.252082e-04 1.876437e-02        Pdik1l
ENSMUSG00000035329 5.680295e-11 2.952751e-08        Fbxo33
ENSMUSG00000003032 8.896410e-08 2.911762e-05          Klf4
ENSMUSG00000076431 4.947581e-12 2.914785e-09          Sox4
ENSMUSG00000024042 1.206238e-05 2.920417e-03          Sik1
ENSMUSG00000020893 2.613226e-07 8.102833e-05          Per1
ENSMUSG00000072893 4.822533e-06 1.235267e-03 4933439C10Rik
ENSMUSG00000038894 3.464102e-09 1.457727e-06          Irs2
ENSMUSG00000090877 2.924759e-05 6.037280e-03        Hspa1b
ENSMUSG00000026814 2.562945e-05 5.392559e-03           Eng
ENSMUSG00000051022 4.677537e-05 9.084702e-03        Hs3st1
```

```
write.csv(treatment4, file = "genelist.csv")
genes4 <- rownames(treatment4)

dlist3 <- bitr(genes4, fromType = "ENSEMBL", toType= 'ENTREZID', OrgDb = org.Mm.eg.db)
```

```
'select()' returned 1:1 mapping between keys and columns
```

```
Warning in bitr(genes4, fromType = "ENSEMBL", toType = "ENTREZID", OrgDb =
org.Mm.eg.db): 20.91% of input gene IDs are fail to map...
```

```
dlist4 <- dlist3$ENTREZID
dlist4
```

```
 [1] "225872"    "13656"     "387150"    "11838"     "23882"     "227885"   
 [7] "16007"     "80981"     "100504231" "15370"     "56501"     "100142658"
[13] "14281"     "13654"     "19725"     "16477"     "620695"    "12227"    
[19] "19225"     "320019"    "19735"     "14282"     "70924"     "225631"   
[25] "17684"     "19252"     "71198"     "330301"    "13653"     "67603"    
[31] "18167"     "15936"     "19366"     "105732"    "11554"     "270110"   
[37] "229599"    "53324"     "14313"     "211770"    "81489"     "105734"   
[43] "240672"    "56380"     "170770"    "68895"     "69944"     "16476"    
[49] "18387"     "18030"     "232685"    "12064"     "235627"    "73747"    
[55] "18109"     "24064"     "27220"     "19736"     "223775"    "14284"    
[61] "11910"     "320563"    "22057"     "319520"    "20556"     "56405"    
[67] "99929"     "74194"     "18124"     "20393"     "241528"    "14748"    
[73] "58226"     "237387"    "108797"    "24063"     "230809"    "70611"    
[79] "16600"     "20677"     "17691"     "18626"     "74476"     "384783"   
[85] "15511"     "13805"     "15476"
```

```
#BP pathway 
GO_results <- enrichGO(genes4, OrgDb= org.Mm.eg.db, keyType = 'ENSEMBL', ont = "BP")
#getting rid of redundant terms 
bp <- pairwise_termsim(GO_results)
bp4 <- simplify(bp, cutoff=0.7, by="p.adjust", select_fun=min)
bp5 <- as.data.frame(bp4)
bp5
```

```
                   ID
GO:0060537 GO:0060537
GO:0043409 GO:0043409
GO:0007611 GO:0007611
GO:0035914 GO:0035914
GO:0007623 GO:0007623
GO:0007178 GO:0007178
GO:0090257 GO:0090257
GO:0042326 GO:0042326
GO:0007622 GO:0007622
GO:0070371 GO:0070371
GO:0001933 GO:0001933
GO:1900744 GO:1900744
GO:0043618 GO:0043618
GO:0060420 GO:0060420
GO:0090287 GO:0090287
GO:1903524 GO:1903524
GO:0051147 GO:0051147
GO:0003012 GO:0003012
GO:0071559 GO:0071559
GO:0038066 GO:0038066
GO:0033605 GO:0033605
GO:0061469 GO:0061469
GO:0045444 GO:0045444
GO:0051348 GO:0051348
GO:0033002 GO:0033002
GO:2000630 GO:2000630
GO:0099565 GO:0099565
GO:0048660 GO:0048660
GO:0035051 GO:0035051
GO:0034405 GO:0034405
GO:0051952 GO:0051952
GO:0003007 GO:0003007
GO:0046697 GO:0046697
GO:0031644 GO:0031644
GO:0043281 GO:0043281
GO:0060078 GO:0060078
GO:0007631 GO:0007631
GO:0032922 GO:0032922
GO:0035265 GO:0035265
GO:0050805 GO:0050805
GO:0061614 GO:0061614
GO:0032891 GO:0032891
GO:0007492 GO:0007492
GO:0043154 GO:0043154
GO:0071277 GO:0071277
GO:0019233 GO:0019233
GO:1902074 GO:1902074
GO:0035335 GO:0035335
GO:0061050 GO:0061050
GO:0010959 GO:0010959
GO:0001558 GO:0001558
GO:1903320 GO:1903320
GO:0044342 GO:0044342
GO:0030099 GO:0030099
GO:0051346 GO:0051346
GO:0045926 GO:0045926
GO:0038179 GO:0038179
GO:0048520 GO:0048520
GO:0001570 GO:0001570
GO:0001706 GO:0001706
GO:0043434 GO:0043434
GO:0010586 GO:0010586
GO:0034764 GO:0034764
GO:1903532 GO:1903532
GO:0046877 GO:0046877
GO:0060136 GO:0060136
GO:1902746 GO:1902746
GO:0001659 GO:0001659
GO:0045923 GO:0045923
GO:1901214 GO:1901214
GO:0043523 GO:0043523
GO:1902075 GO:1902075
GO:0045639 GO:0045639
GO:0043405 GO:0043405
GO:0090084 GO:0090084
GO:2000322 GO:2000322
GO:0009749 GO:0009749
GO:0060213 GO:0060213
GO:0060840 GO:0060840
GO:0002042 GO:0002042
GO:0045777 GO:0045777
GO:0009314 GO:0009314
GO:0015816 GO:0015816
GO:0042752 GO:0042752
GO:0042692 GO:0042692
GO:0051402 GO:0051402
GO:0007406 GO:0007406
GO:0042596 GO:0042596
GO:0070997 GO:0070997
GO:0042303 GO:0042303
GO:0042633 GO:0042633
GO:0051592 GO:0051592
GO:0071498 GO:0071498
GO:0042180 GO:0042180
GO:0051385 GO:0051385
GO:0007389 GO:0007389
GO:0055117 GO:0055117
GO:2000112 GO:2000112
GO:0007193 GO:0007193
GO:0048148 GO:0048148
GO:0006700 GO:0006700
GO:0010958 GO:0010958
GO:0035994 GO:0035994
GO:1903789 GO:1903789
GO:0007422 GO:0007422
GO:0050432 GO:0050432
GO:0098773 GO:0098773
GO:0010565 GO:0010565
GO:0060602 GO:0060602
GO:0008544 GO:0008544
GO:0120255 GO:0120255
GO:0019216 GO:0019216
GO:0043270 GO:0043270
GO:0060562 GO:0060562
GO:0001829 GO:0001829
GO:0001678 GO:0001678
GO:0009648 GO:0009648
GO:0048588 GO:0048588
GO:0035296 GO:0035296
GO:0097746 GO:0097746
GO:0060716 GO:0060716
GO:0072567 GO:0072567
GO:2000341 GO:2000341
GO:0035150 GO:0035150
GO:0031214 GO:0031214
GO:0002088 GO:0002088
GO:0001503 GO:0001503
GO:0045187 GO:0045187
GO:0070841 GO:0070841
GO:0042886 GO:0042886
GO:1901379 GO:1901379
GO:0030534 GO:0030534
GO:0042634 GO:0042634
GO:0010717 GO:0010717
GO:0046888 GO:0046888
GO:0070482 GO:0070482
GO:0120162 GO:0120162
GO:0030336 GO:0030336
GO:2000178 GO:2000178
GO:0010721 GO:0010721
GO:0014910 GO:0014910
GO:0002028 GO:0002028
GO:0032651 GO:0032651
GO:0002053 GO:0002053
GO:0051085 GO:0051085
GO:2000108 GO:2000108
GO:0003156 GO:0003156
GO:0008340 GO:0008340
GO:0010631 GO:0010631
GO:0090132 GO:0090132
GO:0002790 GO:0002790
GO:0043588 GO:0043588
GO:0098739 GO:0098739
GO:0032611 GO:0032611
GO:0000132 GO:0000132
GO:0046686 GO:0046686
GO:0034248 GO:0034248
GO:0048562 GO:0048562
GO:0001822 GO:0001822
                                                                                         Description
GO:0060537                                                                 muscle tissue development
GO:0043409                                                       negative regulation of MAPK cascade
GO:0007611                                                                        learning or memory
GO:0035914                                                      skeletal muscle cell differentiation
GO:0007623                                                                          circadian rhythm
GO:0007178                  transmembrane receptor protein serine/threonine kinase signaling pathway
GO:0090257                                                       regulation of muscle system process
GO:0042326                                                    negative regulation of phosphorylation
GO:0007622                                                                         rhythmic behavior
GO:0070371                                                                     ERK1 and ERK2 cascade
GO:0001933                                            negative regulation of protein phosphorylation
GO:1900744                                                             regulation of p38MAPK cascade
GO:0043618         regulation of transcription from RNA polymerase II promoter in response to stress
GO:0060420                                                                regulation of heart growth
GO:0090287                                 regulation of cellular response to growth factor stimulus
GO:1903524                                                  positive regulation of blood circulation
GO:0051147                                                 regulation of muscle cell differentiation
GO:0003012                                                                     muscle system process
GO:0071559                                               response to transforming growth factor beta
GO:0038066                                                                           p38MAPK cascade
GO:0033605                                            positive regulation of catecholamine secretion
GO:0061469                                        regulation of type B pancreatic cell proliferation
GO:0045444                                                                  fat cell differentiation
GO:0051348                                               negative regulation of transferase activity
GO:0033002                                                                 muscle cell proliferation
GO:2000630                                            positive regulation of miRNA metabolic process
GO:0099565                                              chemical synaptic transmission, postsynaptic
GO:0048660                                            regulation of smooth muscle cell proliferation
GO:0035051                                                                cardiocyte differentiation
GO:0034405                                                            response to fluid shear stress
GO:0051952                                                             regulation of amine transport
GO:0003007                                                                       heart morphogenesis
GO:0046697                                                                           decidualization
GO:0031644                                                      regulation of nervous system process
GO:0043281          regulation of cysteine-type endopeptidase activity involved in apoptotic process
GO:0060078                                             regulation of postsynaptic membrane potential
GO:0007631                                                                          feeding behavior
GO:0032922                                                   circadian regulation of gene expression
GO:0035265                                                                              organ growth
GO:0050805                                              negative regulation of synaptic transmission
GO:0061614                                                                       miRNA transcription
GO:0032891                                             negative regulation of organic acid transport
GO:0007492                                                                      endoderm development
GO:0043154 negative regulation of cysteine-type endopeptidase activity involved in apoptotic process
GO:0071277                                                          cellular response to calcium ion
GO:0019233                                                                sensory perception of pain
GO:1902074                                                                          response to salt
GO:0035335                                                       peptidyl-tyrosine dephosphorylation
GO:0061050                     regulation of cell growth involved in cardiac muscle cell development
GO:0010959                                                         regulation of metal ion transport
GO:0001558                                                                 regulation of cell growth
GO:1903320                regulation of protein modification by small protein conjugation or removal
GO:0044342                                                      type B pancreatic cell proliferation
GO:0030099                                                              myeloid cell differentiation
GO:0051346                                                 negative regulation of hydrolase activity
GO:0045926                                                             negative regulation of growth
GO:0038179                                                            neurotrophin signaling pathway
GO:0048520                                                           positive regulation of behavior
GO:0001570                                                                            vasculogenesis
GO:0001706                                                                        endoderm formation
GO:0043434                                                               response to peptide hormone
GO:0010586                                                                   miRNA metabolic process
GO:0034764                                            positive regulation of transmembrane transport
GO:1903532                                                  positive regulation of secretion by cell
GO:0046877                                                            regulation of saliva secretion
GO:0060136                                            embryonic process involved in female pregnancy
GO:1902746                                             regulation of lens fiber cell differentiation
GO:0001659                                                                   temperature homeostasis
GO:0045923                                       positive regulation of fatty acid metabolic process
GO:1901214                                                                regulation of neuron death
GO:0043523                                                    regulation of neuron apoptotic process
GO:1902075                                                                 cellular response to salt
GO:0045639                                       positive regulation of myeloid cell differentiation
GO:0043405                                                         regulation of MAP kinase activity
GO:0090084                                            negative regulation of inclusion body assembly
GO:2000322                                   regulation of glucocorticoid receptor signaling pathway
GO:0009749                                                                       response to glucose
GO:0060213                   positive regulation of nuclear-transcribed mRNA poly(A) tail shortening
GO:0060840                                                                        artery development
GO:0002042                                         cell migration involved in sprouting angiogenesis
GO:0045777                                                     positive regulation of blood pressure
GO:0009314                                                                     response to radiation
GO:0015816                                                                         glycine transport
GO:0042752                                                            regulation of circadian rhythm
GO:0042692                                                               muscle cell differentiation
GO:0051402                                                                  neuron apoptotic process
GO:0007406                                           negative regulation of neuroblast proliferation
GO:0042596                                                                             fear response
GO:0070997                                                                              neuron death
GO:0042303                                                                             molting cycle
GO:0042633                                                                                hair cycle
GO:0051592                                                                   response to calcium ion
GO:0071498                                                   cellular response to fluid shear stress
GO:0042180                                                         cellular ketone metabolic process
GO:0051385                                                             response to mineralocorticoid
GO:0007389                                                             pattern specification process
GO:0055117                                                  regulation of cardiac muscle contraction
GO:2000112                                 regulation of cellular macromolecule biosynthetic process
GO:0007193                 adenylate cyclase-inhibiting G protein-coupled receptor signaling pathway
GO:0048148                                                            behavioral response to cocaine
GO:0006700                                                  C21-steroid hormone biosynthetic process
GO:0010958                                    regulation of amino acid import across plasma membrane
GO:0035994                                                                response to muscle stretch
GO:1903789                                          regulation of amino acid transmembrane transport
GO:0007422                                                     peripheral nervous system development
GO:0050432                                                                   catecholamine secretion
GO:0098773                                                                skin epidermis development
GO:0010565                                           regulation of cellular ketone metabolic process
GO:0060602                                                        branch elongation of an epithelium
GO:0008544                                                                     epidermis development
GO:0120255                                                    olefinic compound biosynthetic process
GO:0019216                                                     regulation of lipid metabolic process
GO:0043270                                           positive regulation of monoatomic ion transport
GO:0060562                                                             epithelial tube morphogenesis
GO:0001829                                                      trophectodermal cell differentiation
GO:0001678                                                         intracellular glucose homeostasis
GO:0009648                                                                            photoperiodism
GO:0048588                                                                 developmental cell growth
GO:0035296                                                               regulation of tube diameter
GO:0097746                                                         blood vessel diameter maintenance
GO:0060716                                               labyrinthine layer blood vessel development
GO:0072567                                               chemokine (C-X-C motif) ligand 2 production
GO:2000341                                 regulation of chemokine (C-X-C motif) ligand 2 production
GO:0035150                                                                   regulation of tube size
GO:0031214                                                             biomineral tissue development
GO:0002088                                                       lens development in camera-type eye
GO:0001503                                                                              ossification
GO:0045187                                           regulation of circadian sleep/wake cycle, sleep
GO:0070841                                                                   inclusion body assembly
GO:0042886                                                                           amide transport
GO:1901379                                       regulation of potassium ion transmembrane transport
GO:0030534                                                                            adult behavior
GO:0042634                                                                  regulation of hair cycle
GO:0010717                                        regulation of epithelial to mesenchymal transition
GO:0046888                                                  negative regulation of hormone secretion
GO:0070482                                                                 response to oxygen levels
GO:0120162                                         positive regulation of cold-induced thermogenesis
GO:0030336                                                     negative regulation of cell migration
GO:2000178                                negative regulation of neural precursor cell proliferation
GO:0010721                                                   negative regulation of cell development
GO:0014910                                                regulation of smooth muscle cell migration
GO:0002028                                                        regulation of sodium ion transport
GO:0032651                                               regulation of interleukin-1 beta production
GO:0002053                                     positive regulation of mesenchymal cell proliferation
GO:0051085                                            chaperone cofactor-dependent protein refolding
GO:2000108                                        positive regulation of leukocyte apoptotic process
GO:0003156                                                      regulation of animal organ formation
GO:0008340                                                           determination of adult lifespan
GO:0010631                                                                 epithelial cell migration
GO:0090132                                                                      epithelium migration
GO:0002790                                                                         peptide secretion
GO:0043588                                                                          skin development
GO:0098739                                                             import across plasma membrane
GO:0032611                                                             interleukin-1 beta production
GO:0000132                                              establishment of mitotic spindle orientation
GO:0046686                                                                   response to cadmium ion
GO:0034248                                                     regulation of amide metabolic process
GO:0048562                                                             embryonic organ morphogenesis
GO:0001822                                                                        kidney development
           GeneRatio   BgRatio       pvalue     p.adjust       qvalue
GO:0060537     15/84 500/22777 3.970263e-10 9.000586e-07 6.168535e-07
GO:0043409     10/84 185/22777 1.606907e-09 1.821429e-06 1.248313e-06
GO:0007611     11/84 324/22777 3.003551e-08 1.702262e-05 1.166642e-05
GO:0035914      7/84  94/22777 5.683234e-08 2.576778e-05 1.765990e-05
GO:0007623      9/84 226/22777 1.534847e-07 4.970713e-05 3.406669e-05
GO:0007178     11/84 412/22777 3.375510e-07 9.565352e-05 6.555596e-05
GO:0090257      9/84 261/22777 5.165828e-07 1.171093e-04 8.026065e-05
GO:0042326     10/84 405/22777 2.420213e-06 3.469006e-04 2.377477e-04
GO:0007622      5/84  69/22777 5.646906e-06 7.111965e-04 4.874172e-04
GO:0070371      9/84 352/22777 6.031881e-06 7.156320e-04 4.904571e-04
GO:0001933      9/84 354/22777 6.313472e-06 7.156320e-04 4.904571e-04
GO:1900744      4/84  41/22777 1.570759e-05 1.424364e-03 9.761851e-04
GO:0043618      4/84  42/22777 1.731237e-05 1.471364e-03 1.008396e-03
GO:0060420      5/84  91/22777 2.192970e-05 1.680965e-03 1.152046e-03
GO:0090287      8/84 320/22777 2.400396e-05 1.755386e-03 1.203051e-03
GO:1903524      4/84  46/22777 2.495844e-05 1.768149e-03 1.211798e-03
GO:0051147      6/84 161/22777 2.929698e-05 1.953419e-03 1.338772e-03
GO:0003012      9/84 435/22777 3.230624e-05 2.047113e-03 1.402985e-03
GO:0071559      7/84 247/22777 3.606725e-05 2.190793e-03 1.501456e-03
GO:0038066      4/84  51/22777 3.768899e-05 2.190793e-03 1.501456e-03
GO:0033605      3/84  19/22777 4.492852e-05 2.412848e-03 1.653641e-03
GO:0061469      3/84  20/22777 5.271641e-05 2.655736e-03 1.820104e-03
GO:0045444      7/84 268/22777 6.041952e-05 2.914278e-03 1.997295e-03
GO:0051348      7/84 268/22777 6.041952e-05 2.914278e-03 1.997295e-03
GO:0033002      7/84 272/22777 6.631223e-05 3.105264e-03 2.128187e-03
GO:2000630      4/84  59/22777 6.711864e-05 3.105264e-03 2.128187e-03
GO:0099565      5/84 116/22777 7.037672e-05 3.190881e-03 2.186864e-03
GO:0048660      6/84 191/22777 7.593180e-05 3.336387e-03 2.286586e-03
GO:0035051      6/84 195/22777 8.512601e-05 3.641145e-03 2.495452e-03
GO:0034405      3/84  24/22777 9.260273e-05 3.709558e-03 2.542338e-03
GO:0051952      5/84 124/22777 9.654340e-05 3.709558e-03 2.542338e-03
GO:0003007      7/84 291/22777 1.010923e-04 3.756989e-03 2.574845e-03
GO:0046697      3/84  25/22777 1.049505e-04 3.776553e-03 2.588253e-03
GO:0031644      6/84 204/22777 1.090699e-04 3.837620e-03 2.630106e-03
GO:0043281      6/84 210/22777 1.278053e-04 4.186462e-03 2.869184e-03
GO:0060078      5/84 132/22777 1.296162e-04 4.186462e-03 2.869184e-03
GO:0007631      5/84 133/22777 1.342924e-04 4.228346e-03 2.897889e-03
GO:0032922      4/84  74/22777 1.627074e-04 4.910721e-03 3.365554e-03
GO:0035265      6/84 220/22777 1.646294e-04 4.910721e-03 3.365554e-03
GO:0050805      4/84  75/22777 1.713938e-04 4.918353e-03 3.370784e-03
GO:0061614      4/84  75/22777 1.713938e-04 4.918353e-03 3.370784e-03
GO:0032891      3/84  30/22777 1.828112e-04 5.116457e-03 3.506554e-03
GO:0007492      4/84  78/22777 1.994631e-04 5.447987e-03 3.733767e-03
GO:0043154      4/84  81/22777 2.307043e-04 5.943256e-03 4.073200e-03
GO:0071277      4/84  81/22777 2.307043e-04 5.943256e-03 4.073200e-03
GO:0019233      5/84 152/22777 2.502961e-04 6.304680e-03 4.320900e-03
GO:1902074      7/84 340/22777 2.627823e-04 6.361732e-03 4.360001e-03
GO:0035335      3/84  34/22777 2.665922e-04 6.361732e-03 4.360001e-03
GO:0061050      3/84  34/22777 2.665922e-04 6.361732e-03 4.360001e-03
GO:0010959      8/84 460/22777 2.967375e-04 6.864326e-03 4.704453e-03
GO:0001558      8/84 464/22777 3.144345e-04 6.979025e-03 4.783062e-03
GO:1903320      6/84 250/22777 3.274599e-04 7.070015e-03 4.845422e-03
GO:0044342      3/84  38/22777 3.718555e-04 7.952796e-03 5.450434e-03
GO:0030099      8/84 477/22777 3.780130e-04 8.008930e-03 5.488905e-03
GO:0051346      7/84 368/22777 4.230425e-04 8.829091e-03 6.051001e-03
GO:0045926      6/84 267/22777 4.642127e-04 9.352034e-03 6.409399e-03
GO:0038179      3/84  41/22777 4.661579e-04 9.352034e-03 6.409399e-03
GO:0048520      3/84  41/22777 4.661579e-04 9.352034e-03 6.409399e-03
GO:0001570      4/84  99/22777 4.963894e-04 9.619081e-03 6.592419e-03
GO:0001706      3/84  42/22777 5.006844e-04 9.619081e-03 6.592419e-03
GO:0043434      7/84 385/22777 5.533743e-04 9.948832e-03 6.818413e-03
GO:0010586      4/84 102/22777 5.556974e-04 9.948832e-03 6.818413e-03
GO:0034764      6/84 277/22777 5.633126e-04 9.962585e-03 6.827839e-03
GO:1903532      7/84 387/22777 5.706038e-04 9.962585e-03 6.827839e-03
GO:0046877      2/84  10/22777 5.932726e-04 9.962585e-03 6.827839e-03
GO:0060136      2/84  10/22777 5.932726e-04 9.962585e-03 6.827839e-03
GO:1902746      2/84  10/22777 5.932726e-04 9.962585e-03 6.827839e-03
GO:0001659      5/84 192/22777 7.276654e-04 1.119551e-02 7.672821e-03
GO:0045923      3/84  48/22777 7.424204e-04 1.119551e-02 7.672821e-03
GO:1901214      7/84 405/22777 7.457088e-04 1.119551e-02 7.672821e-03
GO:0043523      6/84 294/22777 7.687792e-04 1.146594e-02 7.858158e-03
GO:1902075      5/84 197/22777 8.167968e-04 1.210247e-02 8.294407e-03
GO:0045639      4/84 114/22777 8.438577e-04 1.215620e-02 8.331231e-03
GO:0043405      5/84 199/22777 8.546405e-04 1.215620e-02 8.331231e-03
GO:0090084      2/84  12/22777 8.659705e-04 1.215620e-02 8.331231e-03
GO:2000322      2/84  12/22777 8.659705e-04 1.215620e-02 8.331231e-03
GO:0009749      5/84 200/22777 8.740455e-04 1.215620e-02 8.331231e-03
GO:0060213      2/84  13/22777 1.020970e-03 1.346485e-02 9.228113e-03
GO:0060840      4/84 120/22777 1.021595e-03 1.346485e-02 9.228113e-03
GO:0002042      3/84  55/22777 1.105407e-03 1.440205e-02 9.870421e-03
GO:0045777      3/84  57/22777 1.226347e-03 1.553144e-02 1.064444e-02
GO:0009314      7/84 444/22777 1.271134e-03 1.600923e-02 1.097189e-02
GO:0015816      2/84  15/22777 1.367813e-03 1.656639e-02 1.135375e-02
GO:0042752      4/84 130/22777 1.373834e-03 1.656639e-02 1.135375e-02
GO:0042692      7/84 458/22777 1.518138e-03 1.800086e-02 1.233686e-02
GO:0051402      6/84 337/22777 1.548669e-03 1.800086e-02 1.233686e-02
GO:0007406      2/84  16/22777 1.559476e-03 1.800086e-02 1.233686e-02
GO:0042596      3/84  62/22777 1.564257e-03 1.800086e-02 1.233686e-02
GO:0070997      7/84 461/22777 1.575641e-03 1.804029e-02 1.236388e-02
GO:0042303      4/84 137/22777 1.665812e-03 1.869502e-02 1.281260e-02
GO:0042633      4/84 137/22777 1.665812e-03 1.869502e-02 1.281260e-02
GO:0051592      4/84 137/22777 1.665812e-03 1.869502e-02 1.281260e-02
GO:0071498      2/84  17/22777 1.763179e-03 1.912501e-02 1.310729e-02
GO:0042180      5/84 244/22777 2.101870e-03 2.237061e-02 1.533165e-02
GO:0051385      2/84  19/22777 2.206353e-03 2.283928e-02 1.565286e-02
GO:0007389      7/84 492/22777 2.274955e-03 2.333631e-02 1.599349e-02
GO:0055117      3/84  71/22777 2.308328e-03 2.357198e-02 1.615501e-02
GO:2000112      7/84 494/22777 2.327171e-03 2.365783e-02 1.621385e-02
GO:0007193      3/84  72/22777 2.402329e-03 2.420480e-02 1.658872e-02
GO:0048148      2/84  20/22777 2.445646e-03 2.442414e-02 1.673904e-02
GO:0006700      2/84  21/22777 2.696626e-03 2.570337e-02 1.761576e-02
GO:0010958      2/84  21/22777 2.696626e-03 2.570337e-02 1.761576e-02
GO:0035994      2/84  21/22777 2.696626e-03 2.570337e-02 1.761576e-02
GO:1903789      2/84  21/22777 2.696626e-03 2.570337e-02 1.761576e-02
GO:0007422      3/84  75/22777 2.698457e-03 2.570337e-02 1.761576e-02
GO:0050432      3/84  76/22777 2.801938e-03 2.657738e-02 1.821476e-02
GO:0098773      4/84 159/22777 2.860587e-03 2.690850e-02 1.844169e-02
GO:0010565      4/84 160/22777 2.925802e-03 2.738170e-02 1.876600e-02
GO:0060602      2/84  22/22777 2.959205e-03 2.738170e-02 1.876600e-02
GO:0008544      6/84 388/22777 3.128503e-03 2.883056e-02 1.975897e-02
GO:0120255      2/84  23/22777 3.233297e-03 2.943728e-02 2.017479e-02
GO:0019216      6/84 393/22777 3.331277e-03 3.003411e-02 2.058382e-02
GO:0043270      5/84 274/22777 3.460768e-03 3.088804e-02 2.116906e-02
GO:0060562      6/84 397/22777 3.500443e-03 3.111962e-02 2.132777e-02
GO:0001829      2/84  25/22777 3.815672e-03 3.314225e-02 2.271398e-02
GO:0001678      4/84 176/22777 4.111918e-03 3.488291e-02 2.390693e-02
GO:0009648      2/84  26/22777 4.123784e-03 3.488291e-02 2.390693e-02
GO:0048588      5/84 287/22777 4.211660e-03 3.549380e-02 2.432561e-02
GO:0035296      4/84 179/22777 4.365632e-03 3.617855e-02 2.479490e-02
GO:0097746      4/84 179/22777 4.365632e-03 3.617855e-02 2.479490e-02
GO:0060716      2/84  27/22777 4.443066e-03 3.617855e-02 2.479490e-02
GO:0072567      2/84  27/22777 4.443066e-03 3.617855e-02 2.479490e-02
GO:2000341      2/84  27/22777 4.443066e-03 3.617855e-02 2.479490e-02
GO:0035150      4/84 180/22777 4.452499e-03 3.617855e-02 2.479490e-02
GO:0031214      4/84 181/22777 4.540525e-03 3.676203e-02 2.519479e-02
GO:0002088      3/84  91/22777 4.654065e-03 3.754721e-02 2.573291e-02
GO:0001503      6/84 425/22777 4.869897e-03 3.865760e-02 2.649391e-02
GO:0045187      2/84  29/22777 5.114801e-03 3.957424e-02 2.712213e-02
GO:0070841      2/84  29/22777 5.114801e-03 3.957424e-02 2.712213e-02
GO:0042886      6/84 431/22777 5.208465e-03 4.016187e-02 2.752487e-02
GO:1901379      3/84  96/22777 5.402056e-03 4.118216e-02 2.822412e-02
GO:0030534      4/84 191/22777 5.486110e-03 4.118216e-02 2.822412e-02
GO:0042634      2/84  31/22777 5.830205e-03 4.291258e-02 2.941005e-02
GO:0010717      3/84  99/22777 5.883534e-03 4.302572e-02 2.948760e-02
GO:0046888      3/84  99/22777 5.883534e-03 4.302572e-02 2.948760e-02
GO:0070482      5/84 312/22777 5.971939e-03 4.353179e-02 2.983443e-02
GO:0120162      3/84 100/22777 6.049552e-03 4.383088e-02 3.003941e-02
GO:0030336      5/84 313/22777 6.051639e-03 4.383088e-02 3.003941e-02
GO:2000178      2/84  32/22777 6.204075e-03 4.422842e-02 3.031187e-02
GO:0010721      5/84 316/22777 6.295241e-03 4.471306e-02 3.064401e-02
GO:0014910      3/84 102/22777 6.389948e-03 4.498762e-02 3.083218e-02
GO:0002028      3/84 103/22777 6.564351e-03 4.512504e-02 3.092636e-02
GO:0032651      3/84 103/22777 6.564351e-03 4.512504e-02 3.092636e-02
GO:0002053      2/84  33/22777 6.588614e-03 4.512504e-02 3.092636e-02
GO:0051085      2/84  33/22777 6.588614e-03 4.512504e-02 3.092636e-02
GO:2000108      2/84  33/22777 6.588614e-03 4.512504e-02 3.092636e-02
GO:0003156      2/84  34/22777 6.983740e-03 4.711946e-02 3.229324e-02
GO:0008340      2/84  34/22777 6.983740e-03 4.711946e-02 3.229324e-02
GO:0010631      5/84 326/22777 7.157113e-03 4.772110e-02 3.270557e-02
GO:0090132      5/84 328/22777 7.338910e-03 4.855567e-02 3.327754e-02
GO:0002790      5/84 329/22777 7.431009e-03 4.868814e-02 3.336833e-02
GO:0043588      5/84 331/22777 7.617628e-03 4.899160e-02 3.357630e-02
GO:0098739      4/84 210/22777 7.628600e-03 4.899160e-02 3.357630e-02
GO:0032611      3/84 109/22777 7.670330e-03 4.912045e-02 3.366461e-02
GO:0000132      2/84  36/22777 7.805429e-03 4.938849e-02 3.384831e-02
GO:0046686      2/84  36/22777 7.805429e-03 4.938849e-02 3.384831e-02
GO:0034248      6/84 470/22777 7.845827e-03 4.938849e-02 3.384831e-02
GO:0048562      5/84 334/22777 7.903659e-03 4.949612e-02 3.392207e-02
GO:0001822      5/84 335/22777 8.000643e-03 4.996545e-02 3.424373e-02
                                                                                                                                                                                                                                                                                                 geneID
GO:0060537 ENSMUSG00000023034/ENSMUSG00000021250/ENSMUSG00000037868/ENSMUSG00000020423/ENSMUSG00000026360/ENSMUSG00000039910/ENSMUSG00000038418/ENSMUSG00000035283/ENSMUSG00000044471/ENSMUSG00000048482/ENSMUSG00000038530/ENSMUSG00000026628/ENSMUSG00000034640/ENSMUSG00000024042/ENSMUSG00000026814
GO:0043409                                                                                                ENSMUSG00000026360/ENSMUSG00000024190/ENSMUSG00000019960/ENSMUSG00000034765/ENSMUSG00000022114/ENSMUSG00000026628/ENSMUSG00000031530/ENSMUSG00000037211/ENSMUSG00000003032/ENSMUSG00000020893
GO:0007611                                                                             ENSMUSG00000045903/ENSMUSG00000022602/ENSMUSG00000020423/ENSMUSG00000032487/ENSMUSG00000038418/ENSMUSG00000035283/ENSMUSG00000059991/ENSMUSG00000052684/ENSMUSG00000025905/ENSMUSG00000048482/ENSMUSG00000019970
GO:0035914                                                                                                                                                         ENSMUSG00000023034/ENSMUSG00000021250/ENSMUSG00000037868/ENSMUSG00000020423/ENSMUSG00000039910/ENSMUSG00000038418/ENSMUSG00000026628
GO:0007623                                                                                                                   ENSMUSG00000038418/ENSMUSG00000028004/ENSMUSG00000035283/ENSMUSG00000038550/ENSMUSG00000056749/ENSMUSG00000048482/ENSMUSG00000021647/ENSMUSG00000024042/ENSMUSG00000020893
GO:0007178                                                                             ENSMUSG00000028195/ENSMUSG00000021250/ENSMUSG00000045991/ENSMUSG00000039910/ENSMUSG00000038418/ENSMUSG00000021765/ENSMUSG00000052684/ENSMUSG00000022114/ENSMUSG00000037573/ENSMUSG00000037211/ENSMUSG00000026814
GO:0090257                                                                                                                   ENSMUSG00000023034/ENSMUSG00000032487/ENSMUSG00000026360/ENSMUSG00000028004/ENSMUSG00000035283/ENSMUSG00000038530/ENSMUSG00000028341/ENSMUSG00000024112/ENSMUSG00000003032
GO:0042326                                                                                                ENSMUSG00000021453/ENSMUSG00000026360/ENSMUSG00000024190/ENSMUSG00000019960/ENSMUSG00000032501/ENSMUSG00000052684/ENSMUSG00000022114/ENSMUSG00000037211/ENSMUSG00000038894/ENSMUSG00000026814
GO:0007622                                                                                                                                                                                               ENSMUSG00000037868/ENSMUSG00000038418/ENSMUSG00000028004/ENSMUSG00000035283/ENSMUSG00000038550
GO:0070371                                                                                                                   ENSMUSG00000028195/ENSMUSG00000024190/ENSMUSG00000019960/ENSMUSG00000052684/ENSMUSG00000022114/ENSMUSG00000026628/ENSMUSG00000031530/ENSMUSG00000037211/ENSMUSG00000003032
GO:0001933                                                                                                                   ENSMUSG00000021453/ENSMUSG00000026360/ENSMUSG00000024190/ENSMUSG00000019960/ENSMUSG00000032501/ENSMUSG00000052684/ENSMUSG00000022114/ENSMUSG00000037211/ENSMUSG00000026814
GO:1900744                                                                                                                                                                                                                  ENSMUSG00000021453/ENSMUSG00000024190/ENSMUSG00000025905/ENSMUSG00000020893
GO:0043618                                                                                                                                                                                                                  ENSMUSG00000039910/ENSMUSG00000038418/ENSMUSG00000052684/ENSMUSG00000026628
GO:0060420                                                                                                                                                                                               ENSMUSG00000026360/ENSMUSG00000039910/ENSMUSG00000019960/ENSMUSG00000035283/ENSMUSG00000038530
GO:0090287                                                                                                                                      ENSMUSG00000028195/ENSMUSG00000045991/ENSMUSG00000039910/ENSMUSG00000021765/ENSMUSG00000022114/ENSMUSG00000037573/ENSMUSG00000037211/ENSMUSG00000026814
GO:1903524                                                                                                                                                                                                                  ENSMUSG00000026360/ENSMUSG00000035283/ENSMUSG00000034765/ENSMUSG00000038530
GO:0051147                                                                                                                                                                            ENSMUSG00000026360/ENSMUSG00000035283/ENSMUSG00000048482/ENSMUSG00000038530/ENSMUSG00000024042/ENSMUSG00000026814
GO:0003012                                                                                                                   ENSMUSG00000023034/ENSMUSG00000032487/ENSMUSG00000026360/ENSMUSG00000028004/ENSMUSG00000035283/ENSMUSG00000038530/ENSMUSG00000028341/ENSMUSG00000024112/ENSMUSG00000003032
GO:0071559                                                                                                                                                         ENSMUSG00000021250/ENSMUSG00000045991/ENSMUSG00000039910/ENSMUSG00000052684/ENSMUSG00000022114/ENSMUSG00000037211/ENSMUSG00000026814
GO:0038066                                                                                                                                                                                                                  ENSMUSG00000021453/ENSMUSG00000024190/ENSMUSG00000025905/ENSMUSG00000020893
GO:0033605                                                                                                                                                                                                                                     ENSMUSG00000028004/ENSMUSG00000025905/ENSMUSG00000021647
GO:0061469                                                                                                                                                                                                                                     ENSMUSG00000023034/ENSMUSG00000028341/ENSMUSG00000038894
GO:0045444                                                                                                                                                         ENSMUSG00000023034/ENSMUSG00000037868/ENSMUSG00000032487/ENSMUSG00000026360/ENSMUSG00000035283/ENSMUSG00000028341/ENSMUSG00000003032
GO:0051348                                                                                                                                                         ENSMUSG00000021453/ENSMUSG00000026360/ENSMUSG00000024190/ENSMUSG00000032501/ENSMUSG00000022114/ENSMUSG00000037211/ENSMUSG00000038894
GO:0033002                                                                                                                                                         ENSMUSG00000032487/ENSMUSG00000039910/ENSMUSG00000038418/ENSMUSG00000032501/ENSMUSG00000052684/ENSMUSG00000028341/ENSMUSG00000003032
GO:2000630                                                                                                                                                                                                                  ENSMUSG00000021250/ENSMUSG00000038418/ENSMUSG00000052684/ENSMUSG00000003032
GO:0099565                                                                                                                                                                                               ENSMUSG00000045903/ENSMUSG00000028004/ENSMUSG00000035283/ENSMUSG00000048482/ENSMUSG00000038530
GO:0048660                                                                                                                                                                            ENSMUSG00000032487/ENSMUSG00000038418/ENSMUSG00000032501/ENSMUSG00000052684/ENSMUSG00000028341/ENSMUSG00000003032
GO:0035051                                                                                                                                                                            ENSMUSG00000026360/ENSMUSG00000039910/ENSMUSG00000035283/ENSMUSG00000038530/ENSMUSG00000037211/ENSMUSG00000024042
GO:0034405                                                                                                                                                                                                                                     ENSMUSG00000032487/ENSMUSG00000039910/ENSMUSG00000003032
GO:0051952                                                                                                                                                                                               ENSMUSG00000026360/ENSMUSG00000028004/ENSMUSG00000025905/ENSMUSG00000021647/ENSMUSG00000038530
GO:0003007                                                                                                                                                         ENSMUSG00000028195/ENSMUSG00000039910/ENSMUSG00000028004/ENSMUSG00000052684/ENSMUSG00000037211/ENSMUSG00000076431/ENSMUSG00000026814
GO:0046697                                                                                                                                                                                                                                     ENSMUSG00000052837/ENSMUSG00000032487/ENSMUSG00000039910
GO:0031644                                                                                                                                                                            ENSMUSG00000037868/ENSMUSG00000028004/ENSMUSG00000059991/ENSMUSG00000025905/ENSMUSG00000021647/ENSMUSG00000038530
GO:0043281                                                                                                                                                                            ENSMUSG00000028195/ENSMUSG00000023034/ENSMUSG00000032487/ENSMUSG00000002083/ENSMUSG00000003032/ENSMUSG00000090877
GO:0060078                                                                                                                                                                                               ENSMUSG00000045903/ENSMUSG00000028004/ENSMUSG00000035283/ENSMUSG00000048482/ENSMUSG00000038530
GO:0007631                                                                                                                                                                                               ENSMUSG00000028004/ENSMUSG00000025905/ENSMUSG00000048482/ENSMUSG00000021647/ENSMUSG00000028341
GO:0032922                                                                                                                                                                                                                  ENSMUSG00000038418/ENSMUSG00000038550/ENSMUSG00000021647/ENSMUSG00000020893
GO:0035265                                                                                                                                                                            ENSMUSG00000026360/ENSMUSG00000039910/ENSMUSG00000019960/ENSMUSG00000035283/ENSMUSG00000022114/ENSMUSG00000038530
GO:0050805                                                                                                                                                                                                                  ENSMUSG00000022602/ENSMUSG00000032487/ENSMUSG00000028004/ENSMUSG00000048482
GO:0061614                                                                                                                                                                                                                  ENSMUSG00000021250/ENSMUSG00000038418/ENSMUSG00000052684/ENSMUSG00000003032
GO:0032891                                                                                                                                                                                                                                     ENSMUSG00000026360/ENSMUSG00000038530/ENSMUSG00000038894
GO:0007492                                                                                                                                                                                                                  ENSMUSG00000022602/ENSMUSG00000024190/ENSMUSG00000034765/ENSMUSG00000031530
GO:0043154                                                                                                                                                                                                                  ENSMUSG00000023034/ENSMUSG00000032487/ENSMUSG00000003032/ENSMUSG00000090877
GO:0071277                                                                                                                                                                                                                  ENSMUSG00000021250/ENSMUSG00000052837/ENSMUSG00000003545/ENSMUSG00000052684
GO:0019233                                                                                                                                                                                               ENSMUSG00000032487/ENSMUSG00000028004/ENSMUSG00000035283/ENSMUSG00000025905/ENSMUSG00000024112
GO:1902074                                                                                                                                                         ENSMUSG00000021250/ENSMUSG00000052837/ENSMUSG00000003545/ENSMUSG00000038418/ENSMUSG00000052684/ENSMUSG00000025905/ENSMUSG00000048482
GO:0035335                                                                                                                                                                                                                                     ENSMUSG00000024190/ENSMUSG00000019960/ENSMUSG00000034765
GO:0061050                                                                                                                                                                                                                                     ENSMUSG00000026360/ENSMUSG00000035283/ENSMUSG00000038530
GO:0010959                                                                                                                                      ENSMUSG00000032487/ENSMUSG00000035283/ENSMUSG00000025905/ENSMUSG00000038530/ENSMUSG00000019970/ENSMUSG00000075224/ENSMUSG00000024042/ENSMUSG00000020893
GO:0001558                                                                                                                                      ENSMUSG00000026360/ENSMUSG00000035283/ENSMUSG00000002083/ENSMUSG00000048482/ENSMUSG00000038530/ENSMUSG00000051243/ENSMUSG00000019970/ENSMUSG00000090877
GO:1903320                                                                                                                                                                            ENSMUSG00000038418/ENSMUSG00000032501/ENSMUSG00000022114/ENSMUSG00000035329/ENSMUSG00000076431/ENSMUSG00000090877
GO:0044342                                                                                                                                                                                                                                     ENSMUSG00000023034/ENSMUSG00000028341/ENSMUSG00000038894
GO:0030099                                                                                                                                      ENSMUSG00000021250/ENSMUSG00000052837/ENSMUSG00000039910/ENSMUSG00000032501/ENSMUSG00000052684/ENSMUSG00000056724/ENSMUSG00000021647/ENSMUSG00000090877
GO:0051346                                                                                                                                                         ENSMUSG00000023034/ENSMUSG00000032487/ENSMUSG00000026360/ENSMUSG00000022114/ENSMUSG00000037211/ENSMUSG00000003032/ENSMUSG00000090877
GO:0045926                                                                                                                                                                            ENSMUSG00000026360/ENSMUSG00000039910/ENSMUSG00000035283/ENSMUSG00000002083/ENSMUSG00000038530/ENSMUSG00000090877
GO:0038179                                                                                                                                                                                                                                     ENSMUSG00000048482/ENSMUSG00000022114/ENSMUSG00000037211
GO:0048520                                                                                                                                                                                                                                     ENSMUSG00000028004/ENSMUSG00000025905/ENSMUSG00000028341
GO:0001570                                                                                                                                                                                                                  ENSMUSG00000052837/ENSMUSG00000039910/ENSMUSG00000034640/ENSMUSG00000026814
GO:0001706                                                                                                                                                                                                                                     ENSMUSG00000024190/ENSMUSG00000034765/ENSMUSG00000031530
GO:0043434                                                                                                                                                         ENSMUSG00000023034/ENSMUSG00000037868/ENSMUSG00000038418/ENSMUSG00000025905/ENSMUSG00000028341/ENSMUSG00000019970/ENSMUSG00000038894
GO:0010586                                                                                                                                                                                                                  ENSMUSG00000021250/ENSMUSG00000038418/ENSMUSG00000052684/ENSMUSG00000003032
GO:0034764                                                                                                                                                                            ENSMUSG00000022602/ENSMUSG00000035283/ENSMUSG00000025905/ENSMUSG00000028341/ENSMUSG00000075224/ENSMUSG00000038894
GO:1903532                                                                                                                                                         ENSMUSG00000028004/ENSMUSG00000035283/ENSMUSG00000025905/ENSMUSG00000021647/ENSMUSG00000024112/ENSMUSG00000076431/ENSMUSG00000038894
GO:0046877                                                                                                                                                                                                                                                        ENSMUSG00000035283/ENSMUSG00000025905
GO:0060136                                                                                                                                                                                                                                                        ENSMUSG00000052837/ENSMUSG00000039910
GO:1902746                                                                                                                                                                                                                                                        ENSMUSG00000022114/ENSMUSG00000037211
GO:0001659                                                                                                                                                                                               ENSMUSG00000021453/ENSMUSG00000032487/ENSMUSG00000038418/ENSMUSG00000035283/ENSMUSG00000049649
GO:0045923                                                                                                                                                                                                                                     ENSMUSG00000032487/ENSMUSG00000028341/ENSMUSG00000038894
GO:1901214                                                                                                                                                         ENSMUSG00000021250/ENSMUSG00000020423/ENSMUSG00000038418/ENSMUSG00000002083/ENSMUSG00000052684/ENSMUSG00000048482/ENSMUSG00000028341
GO:0043523                                                                                                                                                                            ENSMUSG00000020423/ENSMUSG00000038418/ENSMUSG00000002083/ENSMUSG00000052684/ENSMUSG00000048482/ENSMUSG00000028341
GO:1902075                                                                                                                                                                                               ENSMUSG00000021250/ENSMUSG00000052837/ENSMUSG00000003545/ENSMUSG00000038418/ENSMUSG00000052684
GO:0045639                                                                                                                                                                                                                  ENSMUSG00000021250/ENSMUSG00000032501/ENSMUSG00000052684/ENSMUSG00000090877
GO:0043405                                                                                                                                                                                               ENSMUSG00000026360/ENSMUSG00000024190/ENSMUSG00000032501/ENSMUSG00000022114/ENSMUSG00000037211
GO:0090084                                                                                                                                                                                                                                                        ENSMUSG00000005483/ENSMUSG00000090877
GO:2000322                                                                                                                                                                                                                                                        ENSMUSG00000048482/ENSMUSG00000020893
GO:0009749                                                                                                                                                                                               ENSMUSG00000038418/ENSMUSG00000025905/ENSMUSG00000035828/ENSMUSG00000076431/ENSMUSG00000038894
GO:0060213                                                                                                                                                                                                                                                        ENSMUSG00000020423/ENSMUSG00000037573
GO:0060840                                                                                                                                                                                                                  ENSMUSG00000037868/ENSMUSG00000039910/ENSMUSG00000076431/ENSMUSG00000026814
GO:0002042                                                                                                                                                                                                                                     ENSMUSG00000023034/ENSMUSG00000032487/ENSMUSG00000003032
GO:0045777                                                                                                                                                                                                                                     ENSMUSG00000035283/ENSMUSG00000021647/ENSMUSG00000026814
GO:0009314                                                                                                                                                         ENSMUSG00000038418/ENSMUSG00000028702/ENSMUSG00000002083/ENSMUSG00000052684/ENSMUSG00000019970/ENSMUSG00000024042/ENSMUSG00000020893
GO:0015816                                                                                                                                                                                                                                                        ENSMUSG00000026360/ENSMUSG00000038530
GO:0042752                                                                                                                                                                                                                  ENSMUSG00000028004/ENSMUSG00000035283/ENSMUSG00000024042/ENSMUSG00000020893
GO:0042692                                                                                                                                                         ENSMUSG00000026360/ENSMUSG00000035283/ENSMUSG00000044471/ENSMUSG00000048482/ENSMUSG00000038530/ENSMUSG00000024042/ENSMUSG00000026814
GO:0051402                                                                                                                                                                            ENSMUSG00000020423/ENSMUSG00000038418/ENSMUSG00000002083/ENSMUSG00000052684/ENSMUSG00000048482/ENSMUSG00000028341
GO:0007406                                                                                                                                                                                                                                                        ENSMUSG00000020423/ENSMUSG00000048482
GO:0042596                                                                                                                                                                                                                                     ENSMUSG00000028004/ENSMUSG00000035283/ENSMUSG00000048482
GO:0070997                                                                                                                                                         ENSMUSG00000021250/ENSMUSG00000020423/ENSMUSG00000038418/ENSMUSG00000002083/ENSMUSG00000052684/ENSMUSG00000048482/ENSMUSG00000028341
GO:0042303                                                                                                                                                                                                                  ENSMUSG00000032487/ENSMUSG00000021765/ENSMUSG00000044471/ENSMUSG00000020893
GO:0042633                                                                                                                                                                                                                  ENSMUSG00000032487/ENSMUSG00000021765/ENSMUSG00000044471/ENSMUSG00000020893
GO:0051592                                                                                                                                                                                                                  ENSMUSG00000021250/ENSMUSG00000052837/ENSMUSG00000003545/ENSMUSG00000052684
GO:0071498                                                                                                                                                                                                                                                        ENSMUSG00000032487/ENSMUSG00000003032
GO:0042180                                                                                                                                                                                               ENSMUSG00000032487/ENSMUSG00000038418/ENSMUSG00000028341/ENSMUSG00000024112/ENSMUSG00000038894
GO:0051385                                                                                                                                                                                                                                                        ENSMUSG00000045903/ENSMUSG00000019970
GO:0007389                                                                                                                                                         ENSMUSG00000022602/ENSMUSG00000037868/ENSMUSG00000020423/ENSMUSG00000039910/ENSMUSG00000021765/ENSMUSG00000037211/ENSMUSG00000026814
GO:0055117                                                                                                                                                                                                                                     ENSMUSG00000026360/ENSMUSG00000035283/ENSMUSG00000024112
GO:2000112                                                                                                                                                         ENSMUSG00000020423/ENSMUSG00000026360/ENSMUSG00000037573/ENSMUSG00000003032/ENSMUSG00000076431/ENSMUSG00000020893/ENSMUSG00000038894
GO:0007193                                                                                                                                                                                                                                     ENSMUSG00000026360/ENSMUSG00000028004/ENSMUSG00000025905
GO:0048148                                                                                                                                                                                                                                                        ENSMUSG00000025905/ENSMUSG00000048482
GO:0006700                                                                                                                                                                                                                                                        ENSMUSG00000038418/ENSMUSG00000024112
GO:0010958                                                                                                                                                                                                                                                        ENSMUSG00000026360/ENSMUSG00000038530
GO:0035994                                                                                                                                                                                                                                                        ENSMUSG00000021250/ENSMUSG00000052684
GO:1903789                                                                                                                                                                                                                                                        ENSMUSG00000026360/ENSMUSG00000038530
GO:0007422                                                                                                                                                                                                                                     ENSMUSG00000037868/ENSMUSG00000045991/ENSMUSG00000039910
GO:0050432                                                                                                                                                                                                                                     ENSMUSG00000028004/ENSMUSG00000025905/ENSMUSG00000021647
GO:0098773                                                                                                                                                                                                                  ENSMUSG00000032487/ENSMUSG00000021765/ENSMUSG00000044471/ENSMUSG00000003032
GO:0010565                                                                                                                                                                                                                  ENSMUSG00000032487/ENSMUSG00000038418/ENSMUSG00000028341/ENSMUSG00000038894
GO:0060602                                                                                                                                                                                                                                                        ENSMUSG00000022114/ENSMUSG00000037211
GO:0008544                                                                                                                                                                            ENSMUSG00000032487/ENSMUSG00000021765/ENSMUSG00000044471/ENSMUSG00000037169/ENSMUSG00000029135/ENSMUSG00000003032
GO:0120255                                                                                                                                                                                                                                                        ENSMUSG00000038418/ENSMUSG00000024112
GO:0019216                                                                                                                                                                            ENSMUSG00000028195/ENSMUSG00000032487/ENSMUSG00000038418/ENSMUSG00000028341/ENSMUSG00000024042/ENSMUSG00000038894
GO:0043270                                                                                                                                                                                               ENSMUSG00000022602/ENSMUSG00000035283/ENSMUSG00000025905/ENSMUSG00000019970/ENSMUSG00000075224
GO:0060562                                                                                                                                                                            ENSMUSG00000039910/ENSMUSG00000037169/ENSMUSG00000022114/ENSMUSG00000037211/ENSMUSG00000076431/ENSMUSG00000026814
GO:0001829                                                                                                                                                                                                                                                        ENSMUSG00000052837/ENSMUSG00000039910
GO:0001678                                                                                                                                                                                                                  ENSMUSG00000025905/ENSMUSG00000021647/ENSMUSG00000035828/ENSMUSG00000076431
GO:0009648                                                                                                                                                                                                                                                        ENSMUSG00000024042/ENSMUSG00000020893
GO:0048588                                                                                                                                                                                               ENSMUSG00000026360/ENSMUSG00000035283/ENSMUSG00000048482/ENSMUSG00000038530/ENSMUSG00000051243
GO:0035296                                                                                                                                                                                                                  ENSMUSG00000032487/ENSMUSG00000026360/ENSMUSG00000035283/ENSMUSG00000034765
GO:0097746                                                                                                                                                                                                                  ENSMUSG00000032487/ENSMUSG00000026360/ENSMUSG00000035283/ENSMUSG00000034765
GO:0060716                                                                                                                                                                                                                                                        ENSMUSG00000028195/ENSMUSG00000052837
GO:0072567                                                                                                                                                                                                                                                        ENSMUSG00000028195/ENSMUSG00000003032
GO:2000341                                                                                                                                                                                                                                                        ENSMUSG00000028195/ENSMUSG00000003032
GO:0035150                                                                                                                                                                                                                  ENSMUSG00000032487/ENSMUSG00000026360/ENSMUSG00000035283/ENSMUSG00000034765
GO:0031214                                                                                                                                                                                                                  ENSMUSG00000028195/ENSMUSG00000032487/ENSMUSG00000046761/ENSMUSG00000044471
GO:0002088                                                                                                                                                                                                                                     ENSMUSG00000039910/ENSMUSG00000022114/ENSMUSG00000037211
GO:0001503                                                                                                                                                                            ENSMUSG00000028195/ENSMUSG00000037868/ENSMUSG00000052837/ENSMUSG00000032487/ENSMUSG00000044471/ENSMUSG00000037573
GO:0045187                                                                                                                                                                                                                                                        ENSMUSG00000028004/ENSMUSG00000035283
GO:0070841                                                                                                                                                                                                                                                        ENSMUSG00000005483/ENSMUSG00000090877
GO:0042886                                                                                                                                                                            ENSMUSG00000028004/ENSMUSG00000048482/ENSMUSG00000021647/ENSMUSG00000035828/ENSMUSG00000076431/ENSMUSG00000038894
GO:1901379                                                                                                                                                                                                                                     ENSMUSG00000025905/ENSMUSG00000038530/ENSMUSG00000075224
GO:0030534                                                                                                                                                                                                                  ENSMUSG00000025905/ENSMUSG00000048482/ENSMUSG00000021647/ENSMUSG00000028341
GO:0042634                                                                                                                                                                                                                                                        ENSMUSG00000021765/ENSMUSG00000020893
GO:0010717                                                                                                                                                                                                                                     ENSMUSG00000022114/ENSMUSG00000037211/ENSMUSG00000026814
GO:0046888                                                                                                                                                                                                                                     ENSMUSG00000025905/ENSMUSG00000021647/ENSMUSG00000035828
GO:0070482                                                                                                                                                                                               ENSMUSG00000065537/ENSMUSG00000039910/ENSMUSG00000038418/ENSMUSG00000029135/ENSMUSG00000026814
GO:0120162                                                                                                                                                                                                                                     ENSMUSG00000021453/ENSMUSG00000035283/ENSMUSG00000049649
GO:0030336                                                                                                                                                                                               ENSMUSG00000039910/ENSMUSG00000024190/ENSMUSG00000032501/ENSMUSG00000003032/ENSMUSG00000026814
GO:2000178                                                                                                                                                                                                                                                        ENSMUSG00000020423/ENSMUSG00000048482
GO:0010721                                                                                                                                                                                               ENSMUSG00000020423/ENSMUSG00000032501/ENSMUSG00000048482/ENSMUSG00000037169/ENSMUSG00000021647
GO:0014910                                                                                                                                                                                                                                     ENSMUSG00000038418/ENSMUSG00000032501/ENSMUSG00000028341
GO:0002028                                                                                                                                                                                                                                     ENSMUSG00000019970/ENSMUSG00000024042/ENSMUSG00000020893
GO:0032651                                                                                                                                                                                                                                     ENSMUSG00000028195/ENSMUSG00000031103/ENSMUSG00000038418
GO:0002053                                                                                                                                                                                                                                                        ENSMUSG00000037169/ENSMUSG00000038894
GO:0051085                                                                                                                                                                                                                                                        ENSMUSG00000005483/ENSMUSG00000090877
GO:2000108                                                                                                                                                                                                                                                        ENSMUSG00000002083/ENSMUSG00000028341
GO:0003156                                                                                                                                                                                                                                                        ENSMUSG00000039910/ENSMUSG00000037211
GO:0008340                                                                                                                                                                                                                                                        ENSMUSG00000028702/ENSMUSG00000002083
GO:0010631                                                                                                                                                                                               ENSMUSG00000023034/ENSMUSG00000032487/ENSMUSG00000052684/ENSMUSG00000003032/ENSMUSG00000038894
GO:0090132                                                                                                                                                                                               ENSMUSG00000023034/ENSMUSG00000032487/ENSMUSG00000052684/ENSMUSG00000003032/ENSMUSG00000038894
GO:0002790                                                                                                                                                                                               ENSMUSG00000028004/ENSMUSG00000021647/ENSMUSG00000035828/ENSMUSG00000076431/ENSMUSG00000038894
GO:0043588                                                                                                                                                                                               ENSMUSG00000032487/ENSMUSG00000021765/ENSMUSG00000044471/ENSMUSG00000029135/ENSMUSG00000003032
GO:0098739                                                                                                                                                                                                                  ENSMUSG00000026360/ENSMUSG00000035283/ENSMUSG00000038530/ENSMUSG00000038894
GO:0032611                                                                                                                                                                                                                                     ENSMUSG00000028195/ENSMUSG00000031103/ENSMUSG00000038418
GO:0000132                                                                                                                                                                                                                                                        ENSMUSG00000022114/ENSMUSG00000037211
GO:0046686                                                                                                                                                                                                                                                        ENSMUSG00000021250/ENSMUSG00000052684
GO:0034248                                                                                                                                                                            ENSMUSG00000028195/ENSMUSG00000020423/ENSMUSG00000026360/ENSMUSG00000037573/ENSMUSG00000076431/ENSMUSG00000020893
GO:0048562                                                                                                                                                                                               ENSMUSG00000039910/ENSMUSG00000037169/ENSMUSG00000022114/ENSMUSG00000028341/ENSMUSG00000026814
GO:0001822                                                                                                                                                                                               ENSMUSG00000038418/ENSMUSG00000048482/ENSMUSG00000034640/ENSMUSG00000037211/ENSMUSG00000076431
           Count
GO:0060537    15
GO:0043409    10
GO:0007611    11
GO:0035914     7
GO:0007623     9
GO:0007178    11
GO:0090257     9
GO:0042326    10
GO:0007622     5
GO:0070371     9
GO:0001933     9
GO:1900744     4
GO:0043618     4
GO:0060420     5
GO:0090287     8
GO:1903524     4
GO:0051147     6
GO:0003012     9
GO:0071559     7
GO:0038066     4
GO:0033605     3
GO:0061469     3
GO:0045444     7
GO:0051348     7
GO:0033002     7
GO:2000630     4
GO:0099565     5
GO:0048660     6
GO:0035051     6
GO:0034405     3
GO:0051952     5
GO:0003007     7
GO:0046697     3
GO:0031644     6
GO:0043281     6
GO:0060078     5
GO:0007631     5
GO:0032922     4
GO:0035265     6
GO:0050805     4
GO:0061614     4
GO:0032891     3
GO:0007492     4
GO:0043154     4
GO:0071277     4
GO:0019233     5
GO:1902074     7
GO:0035335     3
GO:0061050     3
GO:0010959     8
GO:0001558     8
GO:1903320     6
GO:0044342     3
GO:0030099     8
GO:0051346     7
GO:0045926     6
GO:0038179     3
GO:0048520     3
GO:0001570     4
GO:0001706     3
GO:0043434     7
GO:0010586     4
GO:0034764     6
GO:1903532     7
GO:0046877     2
GO:0060136     2
GO:1902746     2
GO:0001659     5
GO:0045923     3
GO:1901214     7
GO:0043523     6
GO:1902075     5
GO:0045639     4
GO:0043405     5
GO:0090084     2
GO:2000322     2
GO:0009749     5
GO:0060213     2
GO:0060840     4
GO:0002042     3
GO:0045777     3
GO:0009314     7
GO:0015816     2
GO:0042752     4
GO:0042692     7
GO:0051402     6
GO:0007406     2
GO:0042596     3
GO:0070997     7
GO:0042303     4
GO:0042633     4
GO:0051592     4
GO:0071498     2
GO:0042180     5
GO:0051385     2
GO:0007389     7
GO:0055117     3
GO:2000112     7
GO:0007193     3
GO:0048148     2
GO:0006700     2
GO:0010958     2
GO:0035994     2
GO:1903789     2
GO:0007422     3
GO:0050432     3
GO:0098773     4
GO:0010565     4
GO:0060602     2
GO:0008544     6
GO:0120255     2
GO:0019216     6
GO:0043270     5
GO:0060562     6
GO:0001829     2
GO:0001678     4
GO:0009648     2
GO:0048588     5
GO:0035296     4
GO:0097746     4
GO:0060716     2
GO:0072567     2
GO:2000341     2
GO:0035150     4
GO:0031214     4
GO:0002088     3
GO:0001503     6
GO:0045187     2
GO:0070841     2
GO:0042886     6
GO:1901379     3
GO:0030534     4
GO:0042634     2
GO:0010717     3
GO:0046888     3
GO:0070482     5
GO:0120162     3
GO:0030336     5
GO:2000178     2
GO:0010721     5
GO:0014910     3
GO:0002028     3
GO:0032651     3
GO:0002053     2
GO:0051085     2
GO:2000108     2
GO:0003156     2
GO:0008340     2
GO:0010631     5
GO:0090132     5
GO:0002790     5
GO:0043588     5
GO:0098739     4
GO:0032611     3
GO:0000132     2
GO:0046686     2
GO:0034248     6
GO:0048562     5
GO:0001822     5
```

```
write.csv(GO_results, "bp_categories1.csv")
as.data.frame(GO_results)
```

```
                   ID
GO:0060537 GO:0060537
GO:0043409 GO:0043409
GO:0070373 GO:0070373
GO:0007611 GO:0007611
GO:0035914 GO:0035914
GO:0050890 GO:0050890
GO:0007623 GO:0007623
GO:0007178 GO:0007178
GO:0007612 GO:0007612
GO:0090257 GO:0090257
GO:0007519 GO:0007519
GO:0007517 GO:0007517
GO:0060538 GO:0060538
GO:0007613 GO:0007613
GO:0042326 GO:0042326
GO:0048511 GO:0048511
GO:0070372 GO:0070372
GO:0007622 GO:0007622
GO:0070371 GO:0070371
GO:0001933 GO:0001933
GO:0010563 GO:0010563
GO:0045936 GO:0045936
GO:0090092 GO:0090092
GO:0007179 GO:0007179
GO:1900744 GO:1900744
GO:0043618 GO:0043618
GO:0090101 GO:0090101
GO:0007616 GO:0007616
GO:0060420 GO:0060420
GO:0033673 GO:0033673
GO:0090287 GO:0090287
GO:1903524 GO:1903524
GO:0043620 GO:0043620
GO:0051147 GO:0051147
GO:0003012 GO:0003012
GO:0071560 GO:0071560
GO:0071559 GO:0071559
GO:0006937 GO:0006937
GO:0038066 GO:0038066
GO:0043502 GO:0043502
GO:1902895 GO:1902895
GO:0033605 GO:0033605
GO:0051153 GO:0051153
GO:0045986 GO:0045986
GO:0061469 GO:0061469
GO:0045444 GO:0045444
GO:0051348 GO:0051348
GO:0033002 GO:0033002
GO:2000630 GO:2000630
GO:0099565 GO:0099565
GO:0048660 GO:0048660
GO:0043407 GO:0043407
GO:0035051 GO:0035051
GO:0008306 GO:0008306
GO:0034405 GO:0034405
GO:0060080 GO:0060080
GO:0048659 GO:0048659
GO:0051952 GO:0051952
GO:0060419 GO:0060419
GO:0046620 GO:0046620
GO:0003007 GO:0003007
GO:0048512 GO:0048512
GO:0046697 GO:0046697
GO:0031644 GO:0031644
GO:0014706 GO:0014706
GO:0006469 GO:0006469
GO:0043500 GO:0043500
GO:0015837 GO:0015837
GO:0043281 GO:0043281
GO:0060078 GO:0060078
GO:0006940 GO:0006940
GO:0007631 GO:0007631
GO:0017015 GO:0017015
GO:1902893 GO:1902893
GO:0032922 GO:0032922
GO:0035265 GO:0035265
GO:1903844 GO:1903844
GO:0050805 GO:0050805
GO:0061614 GO:0061614
GO:0032891 GO:0032891
GO:0045932 GO:0045932
GO:0010611 GO:0010611
GO:0007492 GO:0007492
GO:0014743 GO:0014743
GO:0055022 GO:0055022
GO:0061117 GO:0061117
GO:0043154 GO:0043154
GO:0071277 GO:0071277
GO:0003206 GO:0003206
GO:0019233 GO:0019233
GO:1902074 GO:1902074
GO:0055021 GO:0055021
GO:0003231 GO:0003231
GO:0035335 GO:0035335
GO:0061050 GO:0061050
GO:0060411 GO:0060411
GO:0003209 GO:0003209
GO:0010959 GO:0010959
GO:0006942 GO:0006942
GO:0001558 GO:0001558
GO:0001893 GO:0001893
GO:0003151 GO:0003151
GO:2000628 GO:2000628
GO:1903320 GO:1903320
GO:2000116 GO:2000116
GO:0044342 GO:0044342
GO:0030099 GO:0030099
GO:0051346 GO:0051346
GO:0060079 GO:0060079
GO:0045926 GO:0045926
GO:0003230 GO:0003230
GO:0038179 GO:0038179
GO:0048520 GO:0048520
GO:1903522 GO:1903522
GO:0001570 GO:0001570
GO:0071774 GO:0071774
GO:0001706 GO:0001706
GO:0045823 GO:0045823
GO:0002573 GO:0002573
GO:2000117 GO:2000117
GO:0051154 GO:0051154
GO:0060259 GO:0060259
GO:0048738 GO:0048738
GO:0043434 GO:0043434
GO:0048167 GO:0048167
GO:0010586 GO:0010586
GO:0030509 GO:0030509
GO:0034764 GO:0034764
GO:1903532 GO:1903532
GO:0046621 GO:0046621
GO:0033603 GO:0033603
GO:0046877 GO:0046877
GO:0060136 GO:0060136
GO:0061418 GO:0061418
GO:1902746 GO:1902746
GO:0007369 GO:0007369
GO:0050795 GO:0050795
GO:0001704 GO:0001704
GO:0030510 GO:0030510
GO:0071901 GO:0071901
GO:0097201 GO:0097201
GO:0001659 GO:0001659
GO:0048638 GO:0048638
GO:0003300 GO:0003300
GO:0003298 GO:0003298
GO:0003301 GO:0003301
GO:0045923 GO:0045923
GO:0061049 GO:0061049
GO:0071772 GO:0071772
GO:0071773 GO:0071773
GO:1901214 GO:1901214
GO:0043523 GO:0043523
GO:1902075 GO:1902075
GO:0014897 GO:0014897
GO:0045639 GO:0045639
GO:0048661 GO:0048661
GO:0043405 GO:0043405
GO:0090084 GO:0090084
GO:2000253 GO:2000253
GO:2000322 GO:2000322
GO:0055017 GO:0055017
GO:0003205 GO:0003205
GO:0009749 GO:0009749
GO:0060135 GO:0060135
GO:0014896 GO:0014896
GO:0009746 GO:0009746
GO:0006939 GO:0006939
GO:0055006 GO:0055006
GO:0034284 GO:0034284
GO:0031953 GO:0031953
GO:0060213 GO:0060213
GO:0060840 GO:0060840
GO:0002042 GO:0002042
GO:0045933 GO:0045933
GO:0006936 GO:0006936
GO:0051954 GO:0051954
GO:0031396 GO:0031396
GO:0030857 GO:0030857
GO:0045777 GO:0045777
GO:0009314 GO:0009314
GO:1901652 GO:1901652
GO:0003279 GO:0003279
GO:0048640 GO:0048640
GO:0009743 GO:0009743
GO:0015816 GO:0015816
GO:0060211 GO:0060211
GO:0061052 GO:0061052
GO:0042752 GO:0042752
GO:0051047 GO:0051047
GO:1901216 GO:1901216
GO:0042692 GO:0042692
GO:0051402 GO:0051402
GO:0007406 GO:0007406
GO:0035970 GO:0035970
GO:0042921 GO:0042921
GO:0046541 GO:0046541
GO:0042596 GO:0042596
GO:0070997 GO:0070997
GO:0051146 GO:0051146
GO:0042303 GO:0042303
GO:0042633 GO:0042633
GO:0051592 GO:0051592
GO:0045637 GO:0045637
GO:0050873 GO:0050873
GO:0055001 GO:0055001
GO:0031958 GO:0031958
GO:0051386 GO:0051386
GO:0060413 GO:0060413
GO:0071498 GO:0071498
GO:0050679 GO:0050679
GO:0051403 GO:0051403
GO:0051148 GO:0051148
GO:0042180 GO:0042180
GO:0050433 GO:0050433
GO:0006470 GO:0006470
GO:0071241 GO:0071241
GO:0042391 GO:0042391
GO:0040037 GO:0040037
GO:0051385 GO:0051385
GO:0002761 GO:0002761
GO:0007389 GO:0007389
GO:0055117 GO:0055117
GO:2000112 GO:2000112
GO:0052548 GO:0052548
GO:0007193 GO:0007193
GO:0048148 GO:0048148
GO:0051956 GO:0051956
GO:0002763 GO:0002763
GO:0035904 GO:0035904
GO:0031098 GO:0031098
GO:0048168 GO:0048168
GO:0055007 GO:0055007
GO:0006700 GO:0006700
GO:0010958 GO:0010958
GO:0035994 GO:0035994
GO:0090083 GO:0090083
GO:1903789 GO:1903789
GO:0007422 GO:0007422
GO:0050432 GO:0050432
GO:0032868 GO:0032868
GO:0098773 GO:0098773
GO:0010565 GO:0010565
GO:0045475 GO:0045475
GO:0046321 GO:0046321
GO:0060602 GO:0060602
GO:0008544 GO:0008544
GO:0031649 GO:0031649
GO:0071276 GO:0071276
GO:0120255 GO:0120255
GO:0002791 GO:0002791
GO:0019216 GO:0019216
GO:0030856 GO:0030856
GO:0090087 GO:0090087
GO:0043270 GO:0043270
GO:0060562 GO:0060562
GO:0003283 GO:0003283
GO:0003208 GO:0003208
GO:0048844 GO:0048844
GO:0071383 GO:0071383
GO:0001829 GO:0001829
GO:0043153 GO:0043153
GO:0006941 GO:0006941
GO:0031397 GO:0031397
GO:0001678 GO:0001678
GO:0071248 GO:0071248
GO:0003084 GO:0003084
GO:0003181 GO:0003181
GO:0009648 GO:0009648
GO:0048588 GO:0048588
GO:0043525 GO:0043525
GO:0035296 GO:0035296
GO:0097746 GO:0097746
GO:0003281 GO:0003281
GO:0000289 GO:0000289
GO:0009649 GO:0009649
GO:0060716 GO:0060716
GO:0072567 GO:0072567
GO:2000341 GO:2000341
GO:0035150 GO:0035150
GO:0031214 GO:0031214
GO:0002088 GO:0002088
GO:1900745 GO:1900745
GO:0001890 GO:0001890
GO:0050678 GO:0050678
GO:0001503 GO:0001503
GO:0045834 GO:0045834
GO:0001654 GO:0001654
GO:0021675 GO:0021675
GO:0030512 GO:0030512
GO:0051937 GO:0051937
GO:0150063 GO:0150063
GO:0045187 GO:0045187
GO:0070841 GO:0070841
GO:0042886 GO:0042886
GO:0048880 GO:0048880
GO:0044344 GO:0044344
GO:1901379 GO:1901379
GO:0003171 GO:0003171
GO:0010460 GO:0010460
GO:0048745 GO:0048745
GO:0030534 GO:0030534
GO:0048754 GO:0048754
GO:0010951 GO:0010951
GO:1903321 GO:1903321
GO:1903706 GO:1903706
GO:0021602 GO:0021602
GO:0042634 GO:0042634
GO:0050802 GO:0050802
GO:0010717 GO:0010717
GO:0046888 GO:0046888
GO:0070482 GO:0070482
GO:0120162 GO:0120162
GO:0030336 GO:0030336
GO:0042749 GO:0042749
GO:0048011 GO:0048011
GO:0060317 GO:0060317
GO:0071875 GO:0071875
GO:2000178 GO:2000178
GO:0010721 GO:0010721
GO:0003002 GO:0003002
GO:0043524 GO:0043524
GO:0014910 GO:0014910
GO:0008016 GO:0008016
GO:0002028 GO:0002028
GO:0032651 GO:0032651
GO:0060395 GO:0060395
GO:0002053 GO:0002053
GO:0034260 GO:0034260
GO:0040036 GO:0040036
GO:0051085 GO:0051085
GO:2000108 GO:2000108
GO:0032872 GO:0032872
GO:0090288 GO:0090288
GO:0003156 GO:0003156
GO:0008340 GO:0008340
GO:0022410 GO:0022410
GO:0070302 GO:0070302
GO:0034767 GO:0034767
GO:0010631 GO:0010631
GO:2000146 GO:2000146
GO:0090132 GO:0090132
GO:0030308 GO:0030308
GO:0052547 GO:0052547
GO:0010614 GO:0010614
GO:0045987 GO:0045987
GO:0002790 GO:0002790
GO:0019217 GO:0019217
GO:0090130 GO:0090130
GO:0043588 GO:0043588
GO:0030307 GO:0030307
GO:0098739 GO:0098739
GO:1902107 GO:1902107
GO:1903708 GO:1903708
GO:0032611 GO:0032611
GO:0046890 GO:0046890
GO:0000132 GO:0000132
GO:0010719 GO:0010719
GO:0046686 GO:0046686
GO:0034248 GO:0034248
GO:0032890 GO:0032890
GO:0055013 GO:0055013
GO:0048562 GO:0048562
GO:0001822 GO:0001822
                                                                                               Description
GO:0060537                                                                       muscle tissue development
GO:0043409                                                             negative regulation of MAPK cascade
GO:0070373                                                    negative regulation of ERK1 and ERK2 cascade
GO:0007611                                                                              learning or memory
GO:0035914                                                            skeletal muscle cell differentiation
GO:0050890                                                                                       cognition
GO:0007623                                                                                circadian rhythm
GO:0007178                        transmembrane receptor protein serine/threonine kinase signaling pathway
GO:0007612                                                                                        learning
GO:0090257                                                             regulation of muscle system process
GO:0007519                                                              skeletal muscle tissue development
GO:0007517                                                                        muscle organ development
GO:0060538                                                               skeletal muscle organ development
GO:0007613                                                                                          memory
GO:0042326                                                          negative regulation of phosphorylation
GO:0048511                                                                                rhythmic process
GO:0070372                                                             regulation of ERK1 and ERK2 cascade
GO:0007622                                                                               rhythmic behavior
GO:0070371                                                                           ERK1 and ERK2 cascade
GO:0001933                                                  negative regulation of protein phosphorylation
GO:0010563                                             negative regulation of phosphorus metabolic process
GO:0045936                                              negative regulation of phosphate metabolic process
GO:0090092          regulation of transmembrane receptor protein serine/threonine kinase signaling pathway
GO:0007179                                      transforming growth factor beta receptor signaling pathway
GO:1900744                                                                   regulation of p38MAPK cascade
GO:0043618               regulation of transcription from RNA polymerase II promoter in response to stress
GO:0090101 negative regulation of transmembrane receptor protein serine/threonine kinase signaling pathway
GO:0007616                                                                                long-term memory
GO:0060420                                                                      regulation of heart growth
GO:0033673                                                          negative regulation of kinase activity
GO:0090287                                       regulation of cellular response to growth factor stimulus
GO:1903524                                                        positive regulation of blood circulation
GO:0043620                                 regulation of DNA-templated transcription in response to stress
GO:0051147                                                       regulation of muscle cell differentiation
GO:0003012                                                                           muscle system process
GO:0071560                                   cellular response to transforming growth factor beta stimulus
GO:0071559                                                     response to transforming growth factor beta
GO:0006937                                                                regulation of muscle contraction
GO:0038066                                                                                 p38MAPK cascade
GO:0043502                                                                 regulation of muscle adaptation
GO:1902895                                                      positive regulation of miRNA transcription
GO:0033605                                                  positive regulation of catecholamine secretion
GO:0051153                                              regulation of striated muscle cell differentiation
GO:0045986                                                negative regulation of smooth muscle contraction
GO:0061469                                              regulation of type B pancreatic cell proliferation
GO:0045444                                                                        fat cell differentiation
GO:0051348                                                     negative regulation of transferase activity
GO:0033002                                                                       muscle cell proliferation
GO:2000630                                                  positive regulation of miRNA metabolic process
GO:0099565                                                    chemical synaptic transmission, postsynaptic
GO:0048660                                                  regulation of smooth muscle cell proliferation
GO:0043407                                                      negative regulation of MAP kinase activity
GO:0035051                                                                      cardiocyte differentiation
GO:0008306                                                                            associative learning
GO:0034405                                                                  response to fluid shear stress
GO:0060080                                                               inhibitory postsynaptic potential
GO:0048659                                                                smooth muscle cell proliferation
GO:0051952                                                                   regulation of amine transport
GO:0060419                                                                                    heart growth
GO:0046620                                                                      regulation of organ growth
GO:0003007                                                                             heart morphogenesis
GO:0048512                                                                              circadian behavior
GO:0046697                                                                                 decidualization
GO:0031644                                                            regulation of nervous system process
GO:0014706                                                              striated muscle tissue development
GO:0006469                                                  negative regulation of protein kinase activity
GO:0043500                                                                               muscle adaptation
GO:0015837                                                                                 amine transport
GO:0043281                regulation of cysteine-type endopeptidase activity involved in apoptotic process
GO:0060078                                                   regulation of postsynaptic membrane potential
GO:0006940                                                         regulation of smooth muscle contraction
GO:0007631                                                                                feeding behavior
GO:0017015                        regulation of transforming growth factor beta receptor signaling pathway
GO:1902893                                                               regulation of miRNA transcription
GO:0032922                                                         circadian regulation of gene expression
GO:0035265                                                                                    organ growth
GO:1903844                     regulation of cellular response to transforming growth factor beta stimulus
GO:0050805                                                    negative regulation of synaptic transmission
GO:0061614                                                                             miRNA transcription
GO:0032891                                                   negative regulation of organic acid transport
GO:0045932                                                       negative regulation of muscle contraction
GO:0010611                                                        regulation of cardiac muscle hypertrophy
GO:0007492                                                                            endoderm development
GO:0014743                                                                regulation of muscle hypertrophy
GO:0055022                                             negative regulation of cardiac muscle tissue growth
GO:0061117                                                             negative regulation of heart growth
GO:0043154       negative regulation of cysteine-type endopeptidase activity involved in apoptotic process
GO:0071277                                                                cellular response to calcium ion
GO:0003206                                                                   cardiac chamber morphogenesis
GO:0019233                                                                      sensory perception of pain
GO:1902074                                                                                response to salt
GO:0055021                                                      regulation of cardiac muscle tissue growth
GO:0003231                                                                   cardiac ventricle development
GO:0035335                                                             peptidyl-tyrosine dephosphorylation
GO:0061050                           regulation of cell growth involved in cardiac muscle cell development
GO:0060411                                                                    cardiac septum morphogenesis
GO:0003209                                                                    cardiac atrium morphogenesis
GO:0010959                                                               regulation of metal ion transport
GO:0006942                                                       regulation of striated muscle contraction
GO:0001558                                                                       regulation of cell growth
GO:0001893                                                                   maternal placenta development
GO:0003151                                                                     outflow tract morphogenesis
GO:2000628                                                           regulation of miRNA metabolic process
GO:1903320                      regulation of protein modification by small protein conjugation or removal
GO:2000116                                              regulation of cysteine-type endopeptidase activity
GO:0044342                                                            type B pancreatic cell proliferation
GO:0030099                                                                    myeloid cell differentiation
GO:0051346                                                       negative regulation of hydrolase activity
GO:0060079                                                               excitatory postsynaptic potential
GO:0045926                                                                   negative regulation of growth
GO:0003230                                                                      cardiac atrium development
GO:0038179                                                                  neurotrophin signaling pathway
GO:0048520                                                                 positive regulation of behavior
GO:1903522                                                                 regulation of blood circulation
GO:0001570                                                                                  vasculogenesis
GO:0071774                                                            response to fibroblast growth factor
GO:0001706                                                                              endoderm formation
GO:0045823                                                        positive regulation of heart contraction
GO:0002573                                                               myeloid leukocyte differentiation
GO:2000117                                     negative regulation of cysteine-type endopeptidase activity
GO:0051154                                     negative regulation of striated muscle cell differentiation
GO:0060259                                                                  regulation of feeding behavior
GO:0048738                                                               cardiac muscle tissue development
GO:0043434                                                                     response to peptide hormone
GO:0048167                                                               regulation of synaptic plasticity
GO:0010586                                                                         miRNA metabolic process
GO:0030509                                                                           BMP signaling pathway
GO:0034764                                                  positive regulation of transmembrane transport
GO:1903532                                                        positive regulation of secretion by cell
GO:0046621                                                             negative regulation of organ growth
GO:0033603                                                       positive regulation of dopamine secretion
GO:0046877                                                                  regulation of saliva secretion
GO:0060136                                                  embryonic process involved in female pregnancy
GO:0061418              regulation of transcription from RNA polymerase II promoter in response to hypoxia
GO:1902746                                                   regulation of lens fiber cell differentiation
GO:0007369                                                                                    gastrulation
GO:0050795                                                                          regulation of behavior
GO:0001704                                                                 formation of primary germ layer
GO:0030510                                                             regulation of BMP signaling pathway
GO:0071901                                 negative regulation of protein serine/threonine kinase activity
GO:0097201      negative regulation of transcription from RNA polymerase II promoter in response to stress
GO:0001659                                                                         temperature homeostasis
GO:0048638                                                              regulation of developmental growth
GO:0003300                                                                      cardiac muscle hypertrophy
GO:0003298                                                                physiological muscle hypertrophy
GO:0003301                                                        physiological cardiac muscle hypertrophy
GO:0045923                                             positive regulation of fatty acid metabolic process
GO:0061049                                         cell growth involved in cardiac muscle cell development
GO:0071772                                                                                 response to BMP
GO:0071773                                                               cellular response to BMP stimulus
GO:1901214                                                                      regulation of neuron death
GO:0043523                                                          regulation of neuron apoptotic process
GO:1902075                                                                       cellular response to salt
GO:0014897                                                                     striated muscle hypertrophy
GO:0045639                                             positive regulation of myeloid cell differentiation
GO:0048661                                         positive regulation of smooth muscle cell proliferation
GO:0043405                                                               regulation of MAP kinase activity
GO:0090084                                                  negative regulation of inclusion body assembly
GO:2000253                                                         positive regulation of feeding behavior
GO:2000322                                         regulation of glucocorticoid receptor signaling pathway
GO:0055017                                                                    cardiac muscle tissue growth
GO:0003205                                                                     cardiac chamber development
GO:0009749                                                                             response to glucose
GO:0060135                                                   maternal process involved in female pregnancy
GO:0014896                                                                              muscle hypertrophy
GO:0009746                                                                              response to hexose
GO:0006939                                                                       smooth muscle contraction
GO:0055006                                                                        cardiac cell development
GO:0034284                                                                      response to monosaccharide
GO:0031953                                              negative regulation of protein autophosphorylation
GO:0060213                         positive regulation of nuclear-transcribed mRNA poly(A) tail shortening
GO:0060840                                                                              artery development
GO:0002042                                               cell migration involved in sprouting angiogenesis
GO:0045933                                                       positive regulation of muscle contraction
GO:0006936                                                                              muscle contraction
GO:0051954                                                          positive regulation of amine transport
GO:0031396                                                            regulation of protein ubiquitination
GO:0030857                                          negative regulation of epithelial cell differentiation
GO:0045777                                                           positive regulation of blood pressure
GO:0009314                                                                           response to radiation
GO:1901652                                                                             response to peptide
GO:0003279                                                                      cardiac septum development
GO:0048640                                                     negative regulation of developmental growth
GO:0009743                                                                        response to carbohydrate
GO:0015816                                                                               glycine transport
GO:0060211                                  regulation of nuclear-transcribed mRNA poly(A) tail shortening
GO:0061052                  negative regulation of cell growth involved in cardiac muscle cell development
GO:0042752                                                                  regulation of circadian rhythm
GO:0051047                                                                positive regulation of secretion
GO:1901216                                                             positive regulation of neuron death
GO:0042692                                                                     muscle cell differentiation
GO:0051402                                                                        neuron apoptotic process
GO:0007406                                                 negative regulation of neuroblast proliferation
GO:0035970                                                            peptidyl-threonine dephosphorylation
GO:0042921                                                       glucocorticoid receptor signaling pathway
GO:0046541                                                                                saliva secretion
GO:0042596                                                                                   fear response
GO:0070997                                                                                    neuron death
GO:0051146                                                            striated muscle cell differentiation
GO:0042303                                                                                   molting cycle
GO:0042633                                                                                      hair cycle
GO:0051592                                                                         response to calcium ion
GO:0045637                                                      regulation of myeloid cell differentiation
GO:0050873                                                                  brown fat cell differentiation
GO:0055001                                                                         muscle cell development
GO:0031958                                                       corticosteroid receptor signaling pathway
GO:0051386                                       regulation of neurotrophin TRK receptor signaling pathway
GO:0060413                                                                     atrial septum morphogenesis
GO:0071498                                                         cellular response to fluid shear stress
GO:0050679                                            positive regulation of epithelial cell proliferation
GO:0051403                                                                   stress-activated MAPK cascade
GO:0051148                                              negative regulation of muscle cell differentiation
GO:0042180                                                               cellular ketone metabolic process
GO:0050433                                                           regulation of catecholamine secretion
GO:0006470                                                                       protein dephosphorylation
GO:0071241                                                        cellular response to inorganic substance
GO:0042391                                                                regulation of membrane potential
GO:0040037                      negative regulation of fibroblast growth factor receptor signaling pathway
GO:0051385                                                                   response to mineralocorticoid
GO:0002761                                                 regulation of myeloid leukocyte differentiation
GO:0007389                                                                   pattern specification process
GO:0055117                                                        regulation of cardiac muscle contraction
GO:2000112                                       regulation of cellular macromolecule biosynthetic process
GO:0052548                                                            regulation of endopeptidase activity
GO:0007193                       adenylate cyclase-inhibiting G protein-coupled receptor signaling pathway
GO:0048148                                                                  behavioral response to cocaine
GO:0051956                                                     negative regulation of amino acid transport
GO:0002763                                        positive regulation of myeloid leukocyte differentiation
GO:0035904                                                                               aorta development
GO:0031098                                               stress-activated protein kinase signaling cascade
GO:0048168                                                      regulation of neuronal synaptic plasticity
GO:0055007                                                             cardiac muscle cell differentiation
GO:0006700                                                        C21-steroid hormone biosynthetic process
GO:0010958                                          regulation of amino acid import across plasma membrane
GO:0035994                                                                      response to muscle stretch
GO:0090083                                                           regulation of inclusion body assembly
GO:1903789                                                regulation of amino acid transmembrane transport
GO:0007422                                                           peripheral nervous system development
GO:0050432                                                                         catecholamine secretion
GO:0032868                                                                             response to insulin
GO:0098773                                                                      skin epidermis development
GO:0010565                                                 regulation of cellular ketone metabolic process
GO:0045475                                                                                locomotor rhythm
GO:0046321                                                     positive regulation of fatty acid oxidation
GO:0060602                                                              branch elongation of an epithelium
GO:0008544                                                                           epidermis development
GO:0031649                                                                                 heat generation
GO:0071276                                                                cellular response to cadmium ion
GO:0120255                                                          olefinic compound biosynthetic process
GO:0002791                                                                 regulation of peptide secretion
GO:0019216                                                           regulation of lipid metabolic process
GO:0030856                                                   regulation of epithelial cell differentiation
GO:0090087                                                                 regulation of peptide transport
GO:0043270                                                 positive regulation of monoatomic ion transport
GO:0060562                                                                   epithelial tube morphogenesis
GO:0003283                                                                       atrial septum development
GO:0003208                                                                 cardiac ventricle morphogenesis
GO:0048844                                                                            artery morphogenesis
GO:0071383                                                   cellular response to steroid hormone stimulus
GO:0001829                                                            trophectodermal cell differentiation
GO:0043153                                                   entrainment of circadian clock by photoperiod
GO:0006941                                                                     striated muscle contraction
GO:0031397                                                   negative regulation of protein ubiquitination
GO:0001678                                                               intracellular glucose homeostasis
GO:0071248                                                                  cellular response to metal ion
GO:0003084                                         positive regulation of systemic arterial blood pressure
GO:0003181                                                            atrioventricular valve morphogenesis
GO:0009648                                                                                  photoperiodism
GO:0048588                                                                       developmental cell growth
GO:0043525                                                 positive regulation of neuron apoptotic process
GO:0035296                                                                     regulation of tube diameter
GO:0097746                                                               blood vessel diameter maintenance
GO:0003281                                                                  ventricular septum development
GO:0000289                                                nuclear-transcribed mRNA poly(A) tail shortening
GO:0009649                                                                  entrainment of circadian clock
GO:0060716                                                     labyrinthine layer blood vessel development
GO:0072567                                                     chemokine (C-X-C motif) ligand 2 production
GO:2000341                                       regulation of chemokine (C-X-C motif) ligand 2 production
GO:0035150                                                                         regulation of tube size
GO:0031214                                                                   biomineral tissue development
GO:0002088                                                             lens development in camera-type eye
GO:1900745                                                          positive regulation of p38MAPK cascade
GO:0001890                                                                            placenta development
GO:0050678                                                     regulation of epithelial cell proliferation
GO:0001503                                                                                    ossification
GO:0045834                                                  positive regulation of lipid metabolic process
GO:0001654                                                                                 eye development
GO:0021675                                                                               nerve development
GO:0030512               negative regulation of transforming growth factor beta receptor signaling pathway
GO:0051937                                                                         catecholamine transport
GO:0150063                                                                       visual system development
GO:0045187                                                 regulation of circadian sleep/wake cycle, sleep
GO:0070841                                                                         inclusion body assembly
GO:0042886                                                                                 amide transport
GO:0048880                                                                      sensory system development
GO:0044344                                          cellular response to fibroblast growth factor stimulus
GO:1901379                                             regulation of potassium ion transmembrane transport
GO:0003171                                                              atrioventricular valve development
GO:0010460                                                               positive regulation of heart rate
GO:0048745                                                                smooth muscle tissue development
GO:0030534                                                                                  adult behavior
GO:0048754                                                   branching morphogenesis of an epithelial tube
GO:0010951                                                   negative regulation of endopeptidase activity
GO:1903321             negative regulation of protein modification by small protein conjugation or removal
GO:1903706                                                                       regulation of hemopoiesis
GO:0021602                                                                     cranial nerve morphogenesis
GO:0042634                                                                        regulation of hair cycle
GO:0050802                                                               circadian sleep/wake cycle, sleep
GO:0010717                                              regulation of epithelial to mesenchymal transition
GO:0046888                                                        negative regulation of hormone secretion
GO:0070482                                                                       response to oxygen levels
GO:0120162                                               positive regulation of cold-induced thermogenesis
GO:0030336                                                           negative regulation of cell migration
GO:0042749                                                        regulation of circadian sleep/wake cycle
GO:0048011                                                     neurotrophin TRK receptor signaling pathway
GO:0060317                                                    cardiac epithelial to mesenchymal transition
GO:0071875                                                           adrenergic receptor signaling pathway
GO:2000178                                      negative regulation of neural precursor cell proliferation
GO:0010721                                                         negative regulation of cell development
GO:0003002                                                                                 regionalization
GO:0043524                                                 negative regulation of neuron apoptotic process
GO:0014910                                                      regulation of smooth muscle cell migration
GO:0008016                                                                 regulation of heart contraction
GO:0002028                                                              regulation of sodium ion transport
GO:0032651                                                     regulation of interleukin-1 beta production
GO:0060395                                                                SMAD protein signal transduction
GO:0002053                                           positive regulation of mesenchymal cell proliferation
GO:0034260                                                          negative regulation of GTPase activity
GO:0040036                               regulation of fibroblast growth factor receptor signaling pathway
GO:0051085                                                  chaperone cofactor-dependent protein refolding
GO:2000108                                              positive regulation of leukocyte apoptotic process
GO:0032872                                                     regulation of stress-activated MAPK cascade
GO:0090288                              negative regulation of cellular response to growth factor stimulus
GO:0003156                                                            regulation of animal organ formation
GO:0008340                                                                 determination of adult lifespan
GO:0022410                                                              circadian sleep/wake cycle process
GO:0070302                                 regulation of stress-activated protein kinase signaling cascade
GO:0034767                                   positive regulation of monoatomic ion transmembrane transport
GO:0010631                                                                       epithelial cell migration
GO:2000146                                                            negative regulation of cell motility
GO:0090132                                                                            epithelium migration
GO:0030308                                                              negative regulation of cell growth
GO:0052547                                                                regulation of peptidase activity
GO:0010614                                               negative regulation of cardiac muscle hypertrophy
GO:0045987                                                positive regulation of smooth muscle contraction
GO:0002790                                                                               peptide secretion
GO:0019217                                                      regulation of fatty acid metabolic process
GO:0090130                                                                                tissue migration
GO:0043588                                                                                skin development
GO:0030307                                                              positive regulation of cell growth
GO:0098739                                                                   import across plasma membrane
GO:1902107                                                positive regulation of leukocyte differentiation
GO:1903708                                                              positive regulation of hemopoiesis
GO:0032611                                                                   interleukin-1 beta production
GO:0046890                                                        regulation of lipid biosynthetic process
GO:0000132                                                    establishment of mitotic spindle orientation
GO:0010719                                     negative regulation of epithelial to mesenchymal transition
GO:0046686                                                                         response to cadmium ion
GO:0034248                                                           regulation of amide metabolic process
GO:0032890                                                            regulation of organic acid transport
GO:0055013                                                                 cardiac muscle cell development
GO:0048562                                                                   embryonic organ morphogenesis
GO:0001822                                                                              kidney development
           GeneRatio   BgRatio       pvalue     p.adjust       qvalue
GO:0060537     15/84 500/22777 3.970263e-10 9.000586e-07 6.168535e-07
GO:0043409     10/84 185/22777 1.606907e-09 1.821429e-06 1.248313e-06
GO:0070373      7/84  83/22777 2.381582e-08 1.702262e-05 1.166642e-05
GO:0007611     11/84 324/22777 3.003551e-08 1.702262e-05 1.166642e-05
GO:0035914      7/84  94/22777 5.683234e-08 2.576778e-05 1.765990e-05
GO:0050890     11/84 361/22777 9.002152e-08 3.401313e-05 2.331083e-05
GO:0007623      9/84 226/22777 1.534847e-07 4.970713e-05 3.406669e-05
GO:0007178     11/84 412/22777 3.375510e-07 9.565352e-05 6.555596e-05
GO:0007612      8/84 190/22777 5.131969e-07 1.171093e-04 8.026065e-05
GO:0090257      9/84 261/22777 5.165828e-07 1.171093e-04 8.026065e-05
GO:0007519      8/84 209/22777 1.054083e-06 2.172370e-04 1.488830e-04
GO:0007517     10/84 375/22777 1.214609e-06 2.294599e-04 1.572599e-04
GO:0060538      8/84 222/22777 1.656690e-06 2.889013e-04 1.979979e-04
GO:0007613      7/84 162/22777 2.312985e-06 3.469006e-04 2.377477e-04
GO:0042326     10/84 405/22777 2.420213e-06 3.469006e-04 2.377477e-04
GO:0048511      9/84 315/22777 2.448350e-06 3.469006e-04 2.377477e-04
GO:0070372      9/84 328/22777 3.405224e-06 4.540967e-04 3.112143e-04
GO:0007622      5/84  69/22777 5.646906e-06 7.111965e-04 4.874172e-04
GO:0070371      9/84 352/22777 6.031881e-06 7.156320e-04 4.904571e-04
GO:0001933      9/84 354/22777 6.313472e-06 7.156320e-04 4.904571e-04
GO:0010563     10/84 461/22777 7.584172e-06 7.815145e-04 5.356095e-04
GO:0045936     10/84 461/22777 7.584172e-06 7.815145e-04 5.356095e-04
GO:0090092      8/84 275/22777 8.043965e-06 7.928552e-04 5.433818e-04
GO:0007179      7/84 203/22777 1.021003e-05 9.644224e-04 6.609651e-04
GO:1900744      4/84  41/22777 1.570759e-05 1.424364e-03 9.761851e-04
GO:0043618      4/84  42/22777 1.731237e-05 1.471364e-03 1.008396e-03
GO:0090101      6/84 147/22777 1.752396e-05 1.471364e-03 1.008396e-03
GO:0007616      4/84  43/22777 1.903453e-05 1.541117e-03 1.056202e-03
GO:0060420      5/84  91/22777 2.192970e-05 1.680965e-03 1.152046e-03
GO:0033673      7/84 229/22777 2.224479e-05 1.680965e-03 1.152046e-03
GO:0090287      8/84 320/22777 2.400396e-05 1.755386e-03 1.203051e-03
GO:1903524      4/84  46/22777 2.495844e-05 1.768149e-03 1.211798e-03
GO:0043620      4/84  47/22777 2.720375e-05 1.868815e-03 1.280789e-03
GO:0051147      6/84 161/22777 2.929698e-05 1.953419e-03 1.338772e-03
GO:0003012      9/84 435/22777 3.230624e-05 2.047113e-03 1.402985e-03
GO:0071560      7/84 243/22777 3.250819e-05 2.047113e-03 1.402985e-03
GO:0071559      7/84 247/22777 3.606725e-05 2.190793e-03 1.501456e-03
GO:0006937      6/84 168/22777 3.720028e-05 2.190793e-03 1.501456e-03
GO:0038066      4/84  51/22777 3.768899e-05 2.190793e-03 1.501456e-03
GO:0043502      5/84 103/22777 3.987760e-05 2.260063e-03 1.548930e-03
GO:1902895      4/84  53/22777 4.391588e-05 2.412848e-03 1.653641e-03
GO:0033605      3/84  19/22777 4.492852e-05 2.412848e-03 1.653641e-03
GO:0051153      5/84 106/22777 4.576642e-05 2.412848e-03 1.653641e-03
GO:0045986      3/84  20/22777 5.271641e-05 2.655736e-03 1.820104e-03
GO:0061469      3/84  20/22777 5.271641e-05 2.655736e-03 1.820104e-03
GO:0045444      7/84 268/22777 6.041952e-05 2.914278e-03 1.997295e-03
GO:0051348      7/84 268/22777 6.041952e-05 2.914278e-03 1.997295e-03
GO:0033002      7/84 272/22777 6.631223e-05 3.105264e-03 2.128187e-03
GO:2000630      4/84  59/22777 6.711864e-05 3.105264e-03 2.128187e-03
GO:0099565      5/84 116/22777 7.037672e-05 3.190881e-03 2.186864e-03
GO:0048660      6/84 191/22777 7.593180e-05 3.336387e-03 2.286586e-03
GO:0043407      4/84  61/22777 7.652938e-05 3.336387e-03 2.286586e-03
GO:0035051      6/84 195/22777 8.512601e-05 3.641145e-03 2.495452e-03
GO:0008306      5/84 122/22777 8.939743e-05 3.709558e-03 2.542338e-03
GO:0034405      3/84  24/22777 9.260273e-05 3.709558e-03 2.542338e-03
GO:0060080      3/84  24/22777 9.260273e-05 3.709558e-03 2.542338e-03
GO:0048659      6/84 199/22777 9.518662e-05 3.709558e-03 2.542338e-03
GO:0051952      5/84 124/22777 9.654340e-05 3.709558e-03 2.542338e-03
GO:0060419      5/84 124/22777 9.654340e-05 3.709558e-03 2.542338e-03
GO:0046620      5/84 125/22777 1.002770e-04 3.756989e-03 2.574845e-03
GO:0003007      7/84 291/22777 1.010923e-04 3.756989e-03 2.574845e-03
GO:0048512      4/84  66/22777 1.042231e-04 3.776553e-03 2.588253e-03
GO:0046697      3/84  25/22777 1.049505e-04 3.776553e-03 2.588253e-03
GO:0031644      6/84 204/22777 1.090699e-04 3.837620e-03 2.630106e-03
GO:0014706      7/84 295/22777 1.100332e-04 3.837620e-03 2.630106e-03
GO:0006469      6/84 205/22777 1.120303e-04 3.848071e-03 2.637268e-03
GO:0043500      5/84 129/22777 1.163333e-04 3.936231e-03 2.697689e-03
GO:0015837      5/84 130/22777 1.206389e-04 4.021890e-03 2.756395e-03
GO:0043281      6/84 210/22777 1.278053e-04 4.186462e-03 2.869184e-03
GO:0060078      5/84 132/22777 1.296162e-04 4.186462e-03 2.869184e-03
GO:0006940      4/84  70/22777 1.311155e-04 4.186462e-03 2.869184e-03
GO:0007631      5/84 133/22777 1.342924e-04 4.228346e-03 2.897889e-03
GO:0017015      5/84 137/22777 1.543029e-04 4.728339e-03 3.240558e-03
GO:1902893      4/84  73/22777 1.543437e-04 4.728339e-03 3.240558e-03
GO:0032922      4/84  74/22777 1.627074e-04 4.910721e-03 3.365554e-03
GO:0035265      6/84 220/22777 1.646294e-04 4.910721e-03 3.365554e-03
GO:1903844      5/84 140/22777 1.707491e-04 4.918353e-03 3.370784e-03
GO:0050805      4/84  75/22777 1.713938e-04 4.918353e-03 3.370784e-03
GO:0061614      4/84  75/22777 1.713938e-04 4.918353e-03 3.370784e-03
GO:0032891      3/84  30/22777 1.828112e-04 5.116457e-03 3.506554e-03
GO:0045932      3/84  30/22777 1.828112e-04 5.116457e-03 3.506554e-03
GO:0010611      4/84  77/22777 1.897643e-04 5.246288e-03 3.595533e-03
GO:0007492      4/84  78/22777 1.994631e-04 5.447987e-03 3.733767e-03
GO:0014743      4/84  80/22777 2.199256e-04 5.855995e-03 4.013395e-03
GO:0055022      3/84  32/22777 2.221507e-04 5.855995e-03 4.013395e-03
GO:0061117      3/84  32/22777 2.221507e-04 5.855995e-03 4.013395e-03
GO:0043154      4/84  81/22777 2.307043e-04 5.943256e-03 4.073200e-03
GO:0071277      4/84  81/22777 2.307043e-04 5.943256e-03 4.073200e-03
GO:0003206      5/84 152/22777 2.502961e-04 6.304680e-03 4.320900e-03
GO:0019233      5/84 152/22777 2.502961e-04 6.304680e-03 4.320900e-03
GO:1902074      7/84 340/22777 2.627823e-04 6.361732e-03 4.360001e-03
GO:0055021      4/84  84/22777 2.653216e-04 6.361732e-03 4.360001e-03
GO:0003231      5/84 154/22777 2.659022e-04 6.361732e-03 4.360001e-03
GO:0035335      3/84  34/22777 2.665922e-04 6.361732e-03 4.360001e-03
GO:0061050      3/84  34/22777 2.665922e-04 6.361732e-03 4.360001e-03
GO:0060411      4/84  85/22777 2.776467e-04 6.556512e-03 4.493493e-03
GO:0003209      3/84  35/22777 2.908109e-04 6.796581e-03 4.658024e-03
GO:0010959      8/84 460/22777 2.967375e-04 6.864326e-03 4.704453e-03
GO:0006942      4/84  87/22777 3.035224e-04 6.950356e-03 4.763413e-03
GO:0001558      8/84 464/22777 3.144345e-04 6.979025e-03 4.783062e-03
GO:0001893      3/84  36/22777 3.164059e-04 6.979025e-03 4.783062e-03
GO:0003151      4/84  88/22777 3.170885e-04 6.979025e-03 4.783062e-03
GO:2000628      4/84  88/22777 3.170885e-04 6.979025e-03 4.783062e-03
GO:1903320      6/84 250/22777 3.274599e-04 7.070015e-03 4.845422e-03
GO:2000116      6/84 250/22777 3.274599e-04 7.070015e-03 4.845422e-03
GO:0044342      3/84  38/22777 3.718555e-04 7.952796e-03 5.450434e-03
GO:0030099      8/84 477/22777 3.780130e-04 8.008930e-03 5.488905e-03
GO:0051346      7/84 368/22777 4.230425e-04 8.829091e-03 6.051001e-03
GO:0060079      4/84  95/22777 4.245130e-04 8.829091e-03 6.051001e-03
GO:0045926      6/84 267/22777 4.642127e-04 9.352034e-03 6.409399e-03
GO:0003230      3/84  41/22777 4.661579e-04 9.352034e-03 6.409399e-03
GO:0038179      3/84  41/22777 4.661579e-04 9.352034e-03 6.409399e-03
GO:0048520      3/84  41/22777 4.661579e-04 9.352034e-03 6.409399e-03
GO:1903522      6/84 270/22777 4.923851e-04 9.619081e-03 6.592419e-03
GO:0001570      4/84  99/22777 4.963894e-04 9.619081e-03 6.592419e-03
GO:0071774      4/84  99/22777 4.963894e-04 9.619081e-03 6.592419e-03
GO:0001706      3/84  42/22777 5.006844e-04 9.619081e-03 6.592419e-03
GO:0045823      3/84  42/22777 5.006844e-04 9.619081e-03 6.592419e-03
GO:0002573      6/84 274/22777 5.320032e-04 9.948832e-03 6.818413e-03
GO:2000117      4/84 101/22777 5.353976e-04 9.948832e-03 6.818413e-03
GO:0051154      3/84  43/22777 5.368082e-04 9.948832e-03 6.818413e-03
GO:0060259      3/84  43/22777 5.368082e-04 9.948832e-03 6.818413e-03
GO:0048738      6/84 275/22777 5.422850e-04 9.948832e-03 6.818413e-03
GO:0043434      7/84 385/22777 5.533743e-04 9.948832e-03 6.818413e-03
GO:0048167      7/84 385/22777 5.533743e-04 9.948832e-03 6.818413e-03
GO:0010586      4/84 102/22777 5.556974e-04 9.948832e-03 6.818413e-03
GO:0030509      5/84 181/22777 5.573452e-04 9.948832e-03 6.818413e-03
GO:0034764      6/84 277/22777 5.633126e-04 9.962585e-03 6.827839e-03
GO:1903532      7/84 387/22777 5.706038e-04 9.962585e-03 6.827839e-03
GO:0046621      3/84  44/22777 5.745591e-04 9.962585e-03 6.827839e-03
GO:0033603      2/84  10/22777 5.932726e-04 9.962585e-03 6.827839e-03
GO:0046877      2/84  10/22777 5.932726e-04 9.962585e-03 6.827839e-03
GO:0060136      2/84  10/22777 5.932726e-04 9.962585e-03 6.827839e-03
GO:0061418      2/84  10/22777 5.932726e-04 9.962585e-03 6.827839e-03
GO:1902746      2/84  10/22777 5.932726e-04 9.962585e-03 6.827839e-03
GO:0007369      5/84 185/22777 6.153661e-04 1.025761e-02 7.030034e-03
GO:0050795      4/84 106/22777 6.423918e-04 1.062994e-02 7.285212e-03
GO:0001704      4/84 107/22777 6.654800e-04 1.085355e-02 7.438459e-03
GO:0030510      4/84 107/22777 6.654800e-04 1.085355e-02 7.438459e-03
GO:0071901      4/84 109/22777 7.134110e-04 1.119551e-02 7.672821e-03
GO:0097201      2/84  11/22777 7.233742e-04 1.119551e-02 7.672821e-03
GO:0001659      5/84 192/22777 7.276654e-04 1.119551e-02 7.672821e-03
GO:0048638      7/84 404/22777 7.349819e-04 1.119551e-02 7.672821e-03
GO:0003300      4/84 110/22777 7.382703e-04 1.119551e-02 7.672821e-03
GO:0003298      3/84  48/22777 7.424204e-04 1.119551e-02 7.672821e-03
GO:0003301      3/84  48/22777 7.424204e-04 1.119551e-02 7.672821e-03
GO:0045923      3/84  48/22777 7.424204e-04 1.119551e-02 7.672821e-03
GO:0061049      3/84  48/22777 7.424204e-04 1.119551e-02 7.672821e-03
GO:0071772      5/84 193/22777 7.448792e-04 1.119551e-02 7.672821e-03
GO:0071773      5/84 193/22777 7.448792e-04 1.119551e-02 7.672821e-03
GO:1901214      7/84 405/22777 7.457088e-04 1.119551e-02 7.672821e-03
GO:0043523      6/84 294/22777 7.687792e-04 1.146594e-02 7.858158e-03
GO:1902075      5/84 197/22777 8.167968e-04 1.210247e-02 8.294407e-03
GO:0014897      4/84 114/22777 8.438577e-04 1.215620e-02 8.331231e-03
GO:0045639      4/84 114/22777 8.438577e-04 1.215620e-02 8.331231e-03
GO:0048661      4/84 114/22777 8.438577e-04 1.215620e-02 8.331231e-03
GO:0043405      5/84 199/22777 8.546405e-04 1.215620e-02 8.331231e-03
GO:0090084      2/84  12/22777 8.659705e-04 1.215620e-02 8.331231e-03
GO:2000253      2/84  12/22777 8.659705e-04 1.215620e-02 8.331231e-03
GO:2000322      2/84  12/22777 8.659705e-04 1.215620e-02 8.331231e-03
GO:0055017      4/84 115/22777 8.718332e-04 1.215620e-02 8.331231e-03
GO:0003205      5/84 200/22777 8.740455e-04 1.215620e-02 8.331231e-03
GO:0009749      5/84 200/22777 8.740455e-04 1.215620e-02 8.331231e-03
GO:0060135      3/84  51/22777 8.868140e-04 1.225858e-02 8.401396e-03
GO:0014896      4/84 116/22777 9.004566e-04 1.237173e-02 8.478941e-03
GO:0009746      5/84 203/22777 9.342349e-04 1.275850e-02 8.744012e-03
GO:0006939      4/84 119/22777 9.902970e-04 1.336311e-02 9.158386e-03
GO:0055006      4/84 119/22777 9.902970e-04 1.336311e-02 9.158386e-03
GO:0034284      5/84 206/22777 9.974587e-04 1.338011e-02 9.170034e-03
GO:0031953      2/84  13/22777 1.020970e-03 1.346485e-02 9.228113e-03
GO:0060213      2/84  13/22777 1.020970e-03 1.346485e-02 9.228113e-03
GO:0060840      4/84 120/22777 1.021595e-03 1.346485e-02 9.228113e-03
GO:0002042      3/84  55/22777 1.105407e-03 1.440205e-02 9.870421e-03
GO:0045933      3/84  55/22777 1.105407e-03 1.440205e-02 9.870421e-03
GO:0006936      6/84 318/22777 1.152390e-03 1.492839e-02 1.023115e-02
GO:0051954      3/84  56/22777 1.164885e-03 1.500452e-02 1.028332e-02
GO:0031396      5/84 215/22777 1.206287e-03 1.545001e-02 1.058864e-02
GO:0030857      3/84  57/22777 1.226347e-03 1.553144e-02 1.064444e-02
GO:0045777      3/84  57/22777 1.226347e-03 1.553144e-02 1.064444e-02
GO:0009314      7/84 444/22777 1.271134e-03 1.600923e-02 1.097189e-02
GO:1901652      7/84 446/22777 1.304341e-03 1.633669e-02 1.119632e-02
GO:0003279      4/84 129/22777 1.335274e-03 1.654135e-02 1.133658e-02
GO:0048640      4/84 129/22777 1.335274e-03 1.654135e-02 1.133658e-02
GO:0009743      5/84 221/22777 1.362433e-03 1.656639e-02 1.135375e-02
GO:0015816      2/84  15/22777 1.367813e-03 1.656639e-02 1.135375e-02
GO:0060211      2/84  15/22777 1.367813e-03 1.656639e-02 1.135375e-02
GO:0061052      2/84  15/22777 1.367813e-03 1.656639e-02 1.135375e-02
GO:0042752      4/84 130/22777 1.373834e-03 1.656639e-02 1.135375e-02
GO:0051047      7/84 451/22777 1.390342e-03 1.667675e-02 1.142938e-02
GO:1901216      4/84 133/22777 1.494175e-03 1.782787e-02 1.221830e-02
GO:0042692      7/84 458/22777 1.518138e-03 1.800086e-02 1.233686e-02
GO:0051402      6/84 337/22777 1.548669e-03 1.800086e-02 1.233686e-02
GO:0007406      2/84  16/22777 1.559476e-03 1.800086e-02 1.233686e-02
GO:0035970      2/84  16/22777 1.559476e-03 1.800086e-02 1.233686e-02
GO:0042921      2/84  16/22777 1.559476e-03 1.800086e-02 1.233686e-02
GO:0046541      2/84  16/22777 1.559476e-03 1.800086e-02 1.233686e-02
GO:0042596      3/84  62/22777 1.564257e-03 1.800086e-02 1.233686e-02
GO:0070997      7/84 461/22777 1.575641e-03 1.804029e-02 1.236388e-02
GO:0051146      6/84 340/22777 1.619695e-03 1.845150e-02 1.264570e-02
GO:0042303      4/84 137/22777 1.665812e-03 1.869502e-02 1.281260e-02
GO:0042633      4/84 137/22777 1.665812e-03 1.869502e-02 1.281260e-02
GO:0051592      4/84 137/22777 1.665812e-03 1.869502e-02 1.281260e-02
GO:0045637      5/84 232/22777 1.686814e-03 1.883748e-02 1.291023e-02
GO:0050873      3/84  64/22777 1.714146e-03 1.904887e-02 1.305510e-02
GO:0055001      5/84 234/22777 1.751395e-03 1.912501e-02 1.310729e-02
GO:0031958      2/84  17/22777 1.763179e-03 1.912501e-02 1.310729e-02
GO:0051386      2/84  17/22777 1.763179e-03 1.912501e-02 1.310729e-02
GO:0060413      2/84  17/22777 1.763179e-03 1.912501e-02 1.310729e-02
GO:0071498      2/84  17/22777 1.763179e-03 1.912501e-02 1.310729e-02
GO:0050679      5/84 235/22777 1.784358e-03 1.926257e-02 1.320156e-02
GO:0051403      5/84 242/22777 2.027999e-03 2.178897e-02 1.493303e-02
GO:0051148      3/84  68/22777 2.040168e-03 2.181632e-02 1.495177e-02
GO:0042180      5/84 244/22777 2.101870e-03 2.237061e-02 1.533165e-02
GO:0050433      3/84  69/22777 2.127268e-03 2.253512e-02 1.544440e-02
GO:0006470      5/84 245/22777 2.139534e-03 2.255965e-02 1.546121e-02
GO:0071241      5/84 246/22777 2.177690e-03 2.283928e-02 1.565286e-02
GO:0042391      7/84 489/22777 2.198338e-03 2.283928e-02 1.565286e-02
GO:0040037      2/84  19/22777 2.206353e-03 2.283928e-02 1.565286e-02
GO:0051385      2/84  19/22777 2.206353e-03 2.283928e-02 1.565286e-02
GO:0002761      4/84 149/22777 2.262333e-03 2.331232e-02 1.597705e-02
GO:0007389      7/84 492/22777 2.274955e-03 2.333631e-02 1.599349e-02
GO:0055117      3/84  71/22777 2.308328e-03 2.357198e-02 1.615501e-02
GO:2000112      7/84 494/22777 2.327171e-03 2.365783e-02 1.621385e-02
GO:0052548      6/84 367/22777 2.376150e-03 2.404791e-02 1.648119e-02
GO:0007193      3/84  72/22777 2.402329e-03 2.420480e-02 1.658872e-02
GO:0048148      2/84  20/22777 2.445646e-03 2.442414e-02 1.673904e-02
GO:0051956      2/84  20/22777 2.445646e-03 2.442414e-02 1.673904e-02
GO:0002763      3/84  73/22777 2.498672e-03 2.465174e-02 1.689502e-02
GO:0035904      3/84  73/22777 2.498672e-03 2.465174e-02 1.689502e-02
GO:0031098      5/84 254/22777 2.501059e-03 2.465174e-02 1.689502e-02
GO:0048168      3/84  74/22777 2.597374e-03 2.549025e-02 1.746970e-02
GO:0055007      4/84 156/22777 2.670929e-03 2.570337e-02 1.761576e-02
GO:0006700      2/84  21/22777 2.696626e-03 2.570337e-02 1.761576e-02
GO:0010958      2/84  21/22777 2.696626e-03 2.570337e-02 1.761576e-02
GO:0035994      2/84  21/22777 2.696626e-03 2.570337e-02 1.761576e-02
GO:0090083      2/84  21/22777 2.696626e-03 2.570337e-02 1.761576e-02
GO:1903789      2/84  21/22777 2.696626e-03 2.570337e-02 1.761576e-02
GO:0007422      3/84  75/22777 2.698457e-03 2.570337e-02 1.761576e-02
GO:0050432      3/84  76/22777 2.801938e-03 2.657738e-02 1.821476e-02
GO:0032868      5/84 262/22777 2.858032e-03 2.690850e-02 1.844169e-02
GO:0098773      4/84 159/22777 2.860587e-03 2.690850e-02 1.844169e-02
GO:0010565      4/84 160/22777 2.925802e-03 2.738170e-02 1.876600e-02
GO:0045475      2/84  22/22777 2.959205e-03 2.738170e-02 1.876600e-02
GO:0046321      2/84  22/22777 2.959205e-03 2.738170e-02 1.876600e-02
GO:0060602      2/84  22/22777 2.959205e-03 2.738170e-02 1.876600e-02
GO:0008544      6/84 388/22777 3.128503e-03 2.883056e-02 1.975897e-02
GO:0031649      2/84  23/22777 3.233297e-03 2.943728e-02 2.017479e-02
GO:0071276      2/84  23/22777 3.233297e-03 2.943728e-02 2.017479e-02
GO:0120255      2/84  23/22777 3.233297e-03 2.943728e-02 2.017479e-02
GO:0002791      5/84 270/22777 3.250560e-03 2.947608e-02 2.020137e-02
GO:0019216      6/84 393/22777 3.331277e-03 3.003411e-02 2.058382e-02
GO:0030856      4/84 166/22777 3.338595e-03 3.003411e-02 2.058382e-02
GO:0090087      5/84 272/22777 3.354477e-03 3.005770e-02 2.059999e-02
GO:0043270      5/84 274/22777 3.460768e-03 3.088804e-02 2.116906e-02
GO:0060562      6/84 397/22777 3.500443e-03 3.111962e-02 2.132777e-02
GO:0003283      2/84  24/22777 3.518814e-03 3.116075e-02 2.135596e-02
GO:0003208      3/84  83/22777 3.595012e-03 3.158873e-02 2.164928e-02
GO:0048844      3/84  83/22777 3.595012e-03 3.158873e-02 2.164928e-02
GO:0071383      4/84 170/22777 3.634825e-03 3.181525e-02 2.180452e-02
GO:0001829      2/84  25/22777 3.815672e-03 3.314225e-02 2.271398e-02
GO:0043153      2/84  25/22777 3.815672e-03 3.314225e-02 2.271398e-02
GO:0006941      4/84 173/22777 3.868382e-03 3.347184e-02 2.293986e-02
GO:0031397      3/84  87/22777 4.103705e-03 3.488291e-02 2.390693e-02
GO:0001678      4/84 176/22777 4.111918e-03 3.488291e-02 2.390693e-02
GO:0071248      4/84 176/22777 4.111918e-03 3.488291e-02 2.390693e-02
GO:0003084      2/84  26/22777 4.123784e-03 3.488291e-02 2.390693e-02
GO:0003181      2/84  26/22777 4.123784e-03 3.488291e-02 2.390693e-02
GO:0009648      2/84  26/22777 4.123784e-03 3.488291e-02 2.390693e-02
GO:0048588      5/84 287/22777 4.211660e-03 3.549380e-02 2.432561e-02
GO:0043525      3/84  88/22777 4.237351e-03 3.557805e-02 2.438335e-02
GO:0035296      4/84 179/22777 4.365632e-03 3.617855e-02 2.479490e-02
GO:0097746      4/84 179/22777 4.365632e-03 3.617855e-02 2.479490e-02
GO:0003281      3/84  89/22777 4.373616e-03 3.617855e-02 2.479490e-02
GO:0000289      2/84  27/22777 4.443066e-03 3.617855e-02 2.479490e-02
GO:0009649      2/84  27/22777 4.443066e-03 3.617855e-02 2.479490e-02
GO:0060716      2/84  27/22777 4.443066e-03 3.617855e-02 2.479490e-02
GO:0072567      2/84  27/22777 4.443066e-03 3.617855e-02 2.479490e-02
GO:2000341      2/84  27/22777 4.443066e-03 3.617855e-02 2.479490e-02
GO:0035150      4/84 180/22777 4.452499e-03 3.617855e-02 2.479490e-02
GO:0031214      4/84 181/22777 4.540525e-03 3.676203e-02 2.519479e-02
GO:0002088      3/84  91/22777 4.654065e-03 3.754721e-02 2.573291e-02
GO:1900745      2/84  28/22777 4.773433e-03 3.837366e-02 2.629932e-02
GO:0001890      4/84 184/22777 4.811631e-03 3.843594e-02 2.634200e-02
GO:0050678      6/84 424/22777 4.815089e-03 3.843594e-02 2.634200e-02
GO:0001503      6/84 425/22777 4.869897e-03 3.865760e-02 2.649391e-02
GO:0045834      4/84 185/22777 4.904366e-03 3.865760e-02 2.649391e-02
GO:0001654      6/84 426/22777 4.925165e-03 3.865760e-02 2.649391e-02
GO:0021675      3/84  93/22777 4.945171e-03 3.865760e-02 2.649391e-02
GO:0030512      3/84  93/22777 4.945171e-03 3.865760e-02 2.649391e-02
GO:0051937      3/84  93/22777 4.945171e-03 3.865760e-02 2.649391e-02
GO:0150063      6/84 429/22777 5.093743e-03 3.957424e-02 2.712213e-02
GO:0045187      2/84  29/22777 5.114801e-03 3.957424e-02 2.712213e-02
GO:0070841      2/84  29/22777 5.114801e-03 3.957424e-02 2.712213e-02
GO:0042886      6/84 431/22777 5.208465e-03 4.016187e-02 2.752487e-02
GO:0048880      6/84 433/22777 5.325078e-03 4.092187e-02 2.804573e-02
GO:0044344      3/84  96/22777 5.402056e-03 4.118216e-02 2.822412e-02
GO:1901379      3/84  96/22777 5.402056e-03 4.118216e-02 2.822412e-02
GO:0003171      2/84  30/22777 5.467086e-03 4.118216e-02 2.822412e-02
GO:0010460      2/84  30/22777 5.467086e-03 4.118216e-02 2.822412e-02
GO:0048745      2/84  30/22777 5.467086e-03 4.118216e-02 2.822412e-02
GO:0030534      4/84 191/22777 5.486110e-03 4.118216e-02 2.822412e-02
GO:0048754      4/84 191/22777 5.486110e-03 4.118216e-02 2.822412e-02
GO:0010951      4/84 192/22777 5.587354e-03 4.180373e-02 2.865011e-02
GO:1903321      3/84  98/22777 5.720288e-03 4.265754e-02 2.923526e-02
GO:1903706      6/84 441/22777 5.810797e-03 4.291258e-02 2.941005e-02
GO:0021602      2/84  31/22777 5.830205e-03 4.291258e-02 2.941005e-02
GO:0042634      2/84  31/22777 5.830205e-03 4.291258e-02 2.941005e-02
GO:0050802      2/84  31/22777 5.830205e-03 4.291258e-02 2.941005e-02
GO:0010717      3/84  99/22777 5.883534e-03 4.302572e-02 2.948760e-02
GO:0046888      3/84  99/22777 5.883534e-03 4.302572e-02 2.948760e-02
GO:0070482      5/84 312/22777 5.971939e-03 4.353179e-02 2.983443e-02
GO:0120162      3/84 100/22777 6.049552e-03 4.383088e-02 3.003941e-02
GO:0030336      5/84 313/22777 6.051639e-03 4.383088e-02 3.003941e-02
GO:0042749      2/84  32/22777 6.204075e-03 4.422842e-02 3.031187e-02
GO:0048011      2/84  32/22777 6.204075e-03 4.422842e-02 3.031187e-02
GO:0060317      2/84  32/22777 6.204075e-03 4.422842e-02 3.031187e-02
GO:0071875      2/84  32/22777 6.204075e-03 4.422842e-02 3.031187e-02
GO:2000178      2/84  32/22777 6.204075e-03 4.422842e-02 3.031187e-02
GO:0010721      5/84 316/22777 6.295241e-03 4.471306e-02 3.064401e-02
GO:0003002      6/84 449/22777 6.328221e-03 4.471306e-02 3.064401e-02
GO:0043524      4/84 199/22777 6.331228e-03 4.471306e-02 3.064401e-02
GO:0014910      3/84 102/22777 6.389948e-03 4.498762e-02 3.083218e-02
GO:0008016      4/84 200/22777 6.442599e-03 4.512504e-02 3.092636e-02
GO:0002028      3/84 103/22777 6.564351e-03 4.512504e-02 3.092636e-02
GO:0032651      3/84 103/22777 6.564351e-03 4.512504e-02 3.092636e-02
GO:0060395      3/84 103/22777 6.564351e-03 4.512504e-02 3.092636e-02
GO:0002053      2/84  33/22777 6.588614e-03 4.512504e-02 3.092636e-02
GO:0034260      2/84  33/22777 6.588614e-03 4.512504e-02 3.092636e-02
GO:0040036      2/84  33/22777 6.588614e-03 4.512504e-02 3.092636e-02
GO:0051085      2/84  33/22777 6.588614e-03 4.512504e-02 3.092636e-02
GO:2000108      2/84  33/22777 6.588614e-03 4.512504e-02 3.092636e-02
GO:0032872      4/84 202/22777 6.669233e-03 4.553961e-02 3.121049e-02
GO:0090288      3/84 105/22777 6.921621e-03 4.711946e-02 3.229324e-02
GO:0003156      2/84  34/22777 6.983740e-03 4.711946e-02 3.229324e-02
GO:0008340      2/84  34/22777 6.983740e-03 4.711946e-02 3.229324e-02
GO:0022410      2/84  34/22777 6.983740e-03 4.711946e-02 3.229324e-02
GO:0070302      4/84 205/22777 7.019012e-03 4.721692e-02 3.236002e-02
GO:0034767      4/84 206/22777 7.138251e-03 4.772110e-02 3.270557e-02
GO:0010631      5/84 326/22777 7.157113e-03 4.772110e-02 3.270557e-02
GO:2000146      5/84 326/22777 7.157113e-03 4.772110e-02 3.270557e-02
GO:0090132      5/84 328/22777 7.338910e-03 4.855567e-02 3.327754e-02
GO:0030308      4/84 208/22777 7.380734e-03 4.855567e-02 3.327754e-02
GO:0052547      6/84 464/22777 7.387893e-03 4.855567e-02 3.327754e-02
GO:0010614      2/84  35/22777 7.389373e-03 4.855567e-02 3.327754e-02
GO:0045987      2/84  35/22777 7.389373e-03 4.855567e-02 3.327754e-02
GO:0002790      5/84 329/22777 7.431009e-03 4.868814e-02 3.336833e-02
GO:0019217      3/84 108/22777 7.478855e-03 4.886042e-02 3.348640e-02
GO:0090130      5/84 330/22777 7.523914e-03 4.899160e-02 3.357630e-02
GO:0043588      5/84 331/22777 7.617628e-03 4.899160e-02 3.357630e-02
GO:0030307      4/84 210/22777 7.628600e-03 4.899160e-02 3.357630e-02
GO:0098739      4/84 210/22777 7.628600e-03 4.899160e-02 3.357630e-02
GO:1902107      4/84 210/22777 7.628600e-03 4.899160e-02 3.357630e-02
GO:1903708      4/84 210/22777 7.628600e-03 4.899160e-02 3.357630e-02
GO:0032611      3/84 109/22777 7.670330e-03 4.912045e-02 3.366461e-02
GO:0046890      4/84 211/22777 7.754566e-03 4.938849e-02 3.384831e-02
GO:0000132      2/84  36/22777 7.805429e-03 4.938849e-02 3.384831e-02
GO:0010719      2/84  36/22777 7.805429e-03 4.938849e-02 3.384831e-02
GO:0046686      2/84  36/22777 7.805429e-03 4.938849e-02 3.384831e-02
GO:0034248      6/84 470/22777 7.845827e-03 4.938849e-02 3.384831e-02
GO:0032890      3/84 110/22777 7.864687e-03 4.938849e-02 3.384831e-02
GO:0055013      3/84 110/22777 7.864687e-03 4.938849e-02 3.384831e-02
GO:0048562      5/84 334/22777 7.903659e-03 4.949612e-02 3.392207e-02
GO:0001822      5/84 335/22777 8.000643e-03 4.996545e-02 3.424373e-02
                                                                                                                                                                                                                                                                                                 geneID
GO:0060537 ENSMUSG00000023034/ENSMUSG00000021250/ENSMUSG00000037868/ENSMUSG00000020423/ENSMUSG00000026360/ENSMUSG00000039910/ENSMUSG00000038418/ENSMUSG00000035283/ENSMUSG00000044471/ENSMUSG00000048482/ENSMUSG00000038530/ENSMUSG00000026628/ENSMUSG00000034640/ENSMUSG00000024042/ENSMUSG00000026814
GO:0043409                                                                                                ENSMUSG00000026360/ENSMUSG00000024190/ENSMUSG00000019960/ENSMUSG00000034765/ENSMUSG00000022114/ENSMUSG00000026628/ENSMUSG00000031530/ENSMUSG00000037211/ENSMUSG00000003032/ENSMUSG00000020893
GO:0070373                                                                                                                                                         ENSMUSG00000024190/ENSMUSG00000019960/ENSMUSG00000022114/ENSMUSG00000026628/ENSMUSG00000031530/ENSMUSG00000037211/ENSMUSG00000003032
GO:0007611                                                                             ENSMUSG00000045903/ENSMUSG00000022602/ENSMUSG00000020423/ENSMUSG00000032487/ENSMUSG00000038418/ENSMUSG00000035283/ENSMUSG00000059991/ENSMUSG00000052684/ENSMUSG00000025905/ENSMUSG00000048482/ENSMUSG00000019970
GO:0035914                                                                                                                                                         ENSMUSG00000023034/ENSMUSG00000021250/ENSMUSG00000037868/ENSMUSG00000020423/ENSMUSG00000039910/ENSMUSG00000038418/ENSMUSG00000026628
GO:0050890                                                                             ENSMUSG00000045903/ENSMUSG00000022602/ENSMUSG00000020423/ENSMUSG00000032487/ENSMUSG00000038418/ENSMUSG00000035283/ENSMUSG00000059991/ENSMUSG00000052684/ENSMUSG00000025905/ENSMUSG00000048482/ENSMUSG00000019970
GO:0007623                                                                                                                   ENSMUSG00000038418/ENSMUSG00000028004/ENSMUSG00000035283/ENSMUSG00000038550/ENSMUSG00000056749/ENSMUSG00000048482/ENSMUSG00000021647/ENSMUSG00000024042/ENSMUSG00000020893
GO:0007178                                                                             ENSMUSG00000028195/ENSMUSG00000021250/ENSMUSG00000045991/ENSMUSG00000039910/ENSMUSG00000038418/ENSMUSG00000021765/ENSMUSG00000052684/ENSMUSG00000022114/ENSMUSG00000037573/ENSMUSG00000037211/ENSMUSG00000026814
GO:0007612                                                                                                                                      ENSMUSG00000045903/ENSMUSG00000020423/ENSMUSG00000032487/ENSMUSG00000059991/ENSMUSG00000052684/ENSMUSG00000025905/ENSMUSG00000048482/ENSMUSG00000019970
GO:0090257                                                                                                                   ENSMUSG00000023034/ENSMUSG00000032487/ENSMUSG00000026360/ENSMUSG00000028004/ENSMUSG00000035283/ENSMUSG00000038530/ENSMUSG00000028341/ENSMUSG00000024112/ENSMUSG00000003032
GO:0007519                                                                                                                                      ENSMUSG00000023034/ENSMUSG00000021250/ENSMUSG00000037868/ENSMUSG00000020423/ENSMUSG00000039910/ENSMUSG00000038418/ENSMUSG00000044471/ENSMUSG00000026628
GO:0007517                                                                                                ENSMUSG00000023034/ENSMUSG00000021250/ENSMUSG00000037868/ENSMUSG00000020423/ENSMUSG00000039910/ENSMUSG00000038418/ENSMUSG00000044471/ENSMUSG00000048482/ENSMUSG00000026628/ENSMUSG00000026814
GO:0060538                                                                                                                                      ENSMUSG00000023034/ENSMUSG00000021250/ENSMUSG00000037868/ENSMUSG00000020423/ENSMUSG00000039910/ENSMUSG00000038418/ENSMUSG00000044471/ENSMUSG00000026628
GO:0007613                                                                                                                                                         ENSMUSG00000045903/ENSMUSG00000022602/ENSMUSG00000032487/ENSMUSG00000038418/ENSMUSG00000035283/ENSMUSG00000048482/ENSMUSG00000019970
GO:0042326                                                                                                ENSMUSG00000021453/ENSMUSG00000026360/ENSMUSG00000024190/ENSMUSG00000019960/ENSMUSG00000032501/ENSMUSG00000052684/ENSMUSG00000022114/ENSMUSG00000037211/ENSMUSG00000038894/ENSMUSG00000026814
GO:0048511                                                                                                                   ENSMUSG00000038418/ENSMUSG00000028004/ENSMUSG00000035283/ENSMUSG00000038550/ENSMUSG00000056749/ENSMUSG00000048482/ENSMUSG00000021647/ENSMUSG00000024042/ENSMUSG00000020893
GO:0070372                                                                                                                   ENSMUSG00000028195/ENSMUSG00000024190/ENSMUSG00000019960/ENSMUSG00000052684/ENSMUSG00000022114/ENSMUSG00000026628/ENSMUSG00000031530/ENSMUSG00000037211/ENSMUSG00000003032
GO:0007622                                                                                                                                                                                               ENSMUSG00000037868/ENSMUSG00000038418/ENSMUSG00000028004/ENSMUSG00000035283/ENSMUSG00000038550
GO:0070371                                                                                                                   ENSMUSG00000028195/ENSMUSG00000024190/ENSMUSG00000019960/ENSMUSG00000052684/ENSMUSG00000022114/ENSMUSG00000026628/ENSMUSG00000031530/ENSMUSG00000037211/ENSMUSG00000003032
GO:0001933                                                                                                                   ENSMUSG00000021453/ENSMUSG00000026360/ENSMUSG00000024190/ENSMUSG00000019960/ENSMUSG00000032501/ENSMUSG00000052684/ENSMUSG00000022114/ENSMUSG00000037211/ENSMUSG00000026814
GO:0010563                                                                                                ENSMUSG00000021453/ENSMUSG00000026360/ENSMUSG00000024190/ENSMUSG00000019960/ENSMUSG00000032501/ENSMUSG00000052684/ENSMUSG00000022114/ENSMUSG00000037211/ENSMUSG00000038894/ENSMUSG00000026814
GO:0045936                                                                                                ENSMUSG00000021453/ENSMUSG00000026360/ENSMUSG00000024190/ENSMUSG00000019960/ENSMUSG00000032501/ENSMUSG00000052684/ENSMUSG00000022114/ENSMUSG00000037211/ENSMUSG00000038894/ENSMUSG00000026814
GO:0090092                                                                                                                                      ENSMUSG00000028195/ENSMUSG00000045991/ENSMUSG00000039910/ENSMUSG00000021765/ENSMUSG00000022114/ENSMUSG00000037573/ENSMUSG00000037211/ENSMUSG00000026814
GO:0007179                                                                                                                                                         ENSMUSG00000021250/ENSMUSG00000045991/ENSMUSG00000039910/ENSMUSG00000052684/ENSMUSG00000022114/ENSMUSG00000037211/ENSMUSG00000026814
GO:1900744                                                                                                                                                                                                                  ENSMUSG00000021453/ENSMUSG00000024190/ENSMUSG00000025905/ENSMUSG00000020893
GO:0043618                                                                                                                                                                                                                  ENSMUSG00000039910/ENSMUSG00000038418/ENSMUSG00000052684/ENSMUSG00000026628
GO:0090101                                                                                                                                                                            ENSMUSG00000045991/ENSMUSG00000021765/ENSMUSG00000022114/ENSMUSG00000037573/ENSMUSG00000037211/ENSMUSG00000026814
GO:0007616                                                                                                                                                                                                                  ENSMUSG00000045903/ENSMUSG00000022602/ENSMUSG00000038418/ENSMUSG00000019970
GO:0060420                                                                                                                                                                                               ENSMUSG00000026360/ENSMUSG00000039910/ENSMUSG00000019960/ENSMUSG00000035283/ENSMUSG00000038530
GO:0033673                                                                                                                                                         ENSMUSG00000021453/ENSMUSG00000026360/ENSMUSG00000024190/ENSMUSG00000032501/ENSMUSG00000022114/ENSMUSG00000037211/ENSMUSG00000038894
GO:0090287                                                                                                                                      ENSMUSG00000028195/ENSMUSG00000045991/ENSMUSG00000039910/ENSMUSG00000021765/ENSMUSG00000022114/ENSMUSG00000037573/ENSMUSG00000037211/ENSMUSG00000026814
GO:1903524                                                                                                                                                                                                                  ENSMUSG00000026360/ENSMUSG00000035283/ENSMUSG00000034765/ENSMUSG00000038530
GO:0043620                                                                                                                                                                                                                  ENSMUSG00000039910/ENSMUSG00000038418/ENSMUSG00000052684/ENSMUSG00000026628
GO:0051147                                                                                                                                                                            ENSMUSG00000026360/ENSMUSG00000035283/ENSMUSG00000048482/ENSMUSG00000038530/ENSMUSG00000024042/ENSMUSG00000026814
GO:0003012                                                                                                                   ENSMUSG00000023034/ENSMUSG00000032487/ENSMUSG00000026360/ENSMUSG00000028004/ENSMUSG00000035283/ENSMUSG00000038530/ENSMUSG00000028341/ENSMUSG00000024112/ENSMUSG00000003032
GO:0071560                                                                                                                                                         ENSMUSG00000021250/ENSMUSG00000045991/ENSMUSG00000039910/ENSMUSG00000052684/ENSMUSG00000022114/ENSMUSG00000037211/ENSMUSG00000026814
GO:0071559                                                                                                                                                         ENSMUSG00000021250/ENSMUSG00000045991/ENSMUSG00000039910/ENSMUSG00000052684/ENSMUSG00000022114/ENSMUSG00000037211/ENSMUSG00000026814
GO:0006937                                                                                                                                                                            ENSMUSG00000023034/ENSMUSG00000032487/ENSMUSG00000026360/ENSMUSG00000028004/ENSMUSG00000035283/ENSMUSG00000024112
GO:0038066                                                                                                                                                                                                                  ENSMUSG00000021453/ENSMUSG00000024190/ENSMUSG00000025905/ENSMUSG00000020893
GO:0043502                                                                                                                                                                                               ENSMUSG00000026360/ENSMUSG00000035283/ENSMUSG00000038530/ENSMUSG00000028341/ENSMUSG00000003032
GO:1902895                                                                                                                                                                                                                  ENSMUSG00000021250/ENSMUSG00000038418/ENSMUSG00000052684/ENSMUSG00000003032
GO:0033605                                                                                                                                                                                                                                     ENSMUSG00000028004/ENSMUSG00000025905/ENSMUSG00000021647
GO:0051153                                                                                                                                                                                               ENSMUSG00000026360/ENSMUSG00000035283/ENSMUSG00000048482/ENSMUSG00000038530/ENSMUSG00000024042
GO:0045986                                                                                                                                                                                                                                     ENSMUSG00000032487/ENSMUSG00000026360/ENSMUSG00000035283
GO:0061469                                                                                                                                                                                                                                     ENSMUSG00000023034/ENSMUSG00000028341/ENSMUSG00000038894
GO:0045444                                                                                                                                                         ENSMUSG00000023034/ENSMUSG00000037868/ENSMUSG00000032487/ENSMUSG00000026360/ENSMUSG00000035283/ENSMUSG00000028341/ENSMUSG00000003032
GO:0051348                                                                                                                                                         ENSMUSG00000021453/ENSMUSG00000026360/ENSMUSG00000024190/ENSMUSG00000032501/ENSMUSG00000022114/ENSMUSG00000037211/ENSMUSG00000038894
GO:0033002                                                                                                                                                         ENSMUSG00000032487/ENSMUSG00000039910/ENSMUSG00000038418/ENSMUSG00000032501/ENSMUSG00000052684/ENSMUSG00000028341/ENSMUSG00000003032
GO:2000630                                                                                                                                                                                                                  ENSMUSG00000021250/ENSMUSG00000038418/ENSMUSG00000052684/ENSMUSG00000003032
GO:0099565                                                                                                                                                                                               ENSMUSG00000045903/ENSMUSG00000028004/ENSMUSG00000035283/ENSMUSG00000048482/ENSMUSG00000038530
GO:0048660                                                                                                                                                                            ENSMUSG00000032487/ENSMUSG00000038418/ENSMUSG00000032501/ENSMUSG00000052684/ENSMUSG00000028341/ENSMUSG00000003032
GO:0043407                                                                                                                                                                                                                  ENSMUSG00000026360/ENSMUSG00000024190/ENSMUSG00000022114/ENSMUSG00000037211
GO:0035051                                                                                                                                                                            ENSMUSG00000026360/ENSMUSG00000039910/ENSMUSG00000035283/ENSMUSG00000038530/ENSMUSG00000037211/ENSMUSG00000024042
GO:0008306                                                                                                                                                                                               ENSMUSG00000020423/ENSMUSG00000059991/ENSMUSG00000025905/ENSMUSG00000048482/ENSMUSG00000019970
GO:0034405                                                                                                                                                                                                                                     ENSMUSG00000032487/ENSMUSG00000039910/ENSMUSG00000003032
GO:0060080                                                                                                                                                                                                                                     ENSMUSG00000045903/ENSMUSG00000035283/ENSMUSG00000048482
GO:0048659                                                                                                                                                                            ENSMUSG00000032487/ENSMUSG00000038418/ENSMUSG00000032501/ENSMUSG00000052684/ENSMUSG00000028341/ENSMUSG00000003032
GO:0051952                                                                                                                                                                                               ENSMUSG00000026360/ENSMUSG00000028004/ENSMUSG00000025905/ENSMUSG00000021647/ENSMUSG00000038530
GO:0060419                                                                                                                                                                                               ENSMUSG00000026360/ENSMUSG00000039910/ENSMUSG00000019960/ENSMUSG00000035283/ENSMUSG00000038530
GO:0046620                                                                                                                                                                                               ENSMUSG00000026360/ENSMUSG00000039910/ENSMUSG00000019960/ENSMUSG00000035283/ENSMUSG00000038530
GO:0003007                                                                                                                                                         ENSMUSG00000028195/ENSMUSG00000039910/ENSMUSG00000028004/ENSMUSG00000052684/ENSMUSG00000037211/ENSMUSG00000076431/ENSMUSG00000026814
GO:0048512                                                                                                                                                                                                                  ENSMUSG00000038418/ENSMUSG00000028004/ENSMUSG00000035283/ENSMUSG00000038550
GO:0046697                                                                                                                                                                                                                                     ENSMUSG00000052837/ENSMUSG00000032487/ENSMUSG00000039910
GO:0031644                                                                                                                                                                            ENSMUSG00000037868/ENSMUSG00000028004/ENSMUSG00000059991/ENSMUSG00000025905/ENSMUSG00000021647/ENSMUSG00000038530
GO:0014706                                                                                                                                                         ENSMUSG00000026360/ENSMUSG00000039910/ENSMUSG00000035283/ENSMUSG00000048482/ENSMUSG00000038530/ENSMUSG00000024042/ENSMUSG00000026814
GO:0006469                                                                                                                                                                            ENSMUSG00000021453/ENSMUSG00000026360/ENSMUSG00000024190/ENSMUSG00000032501/ENSMUSG00000022114/ENSMUSG00000037211
GO:0043500                                                                                                                                                                                               ENSMUSG00000026360/ENSMUSG00000035283/ENSMUSG00000038530/ENSMUSG00000028341/ENSMUSG00000003032
GO:0015837                                                                                                                                                                                               ENSMUSG00000026360/ENSMUSG00000028004/ENSMUSG00000025905/ENSMUSG00000021647/ENSMUSG00000038530
GO:0043281                                                                                                                                                                            ENSMUSG00000028195/ENSMUSG00000023034/ENSMUSG00000032487/ENSMUSG00000002083/ENSMUSG00000003032/ENSMUSG00000090877
GO:0060078                                                                                                                                                                                               ENSMUSG00000045903/ENSMUSG00000028004/ENSMUSG00000035283/ENSMUSG00000048482/ENSMUSG00000038530
GO:0006940                                                                                                                                                                                                                  ENSMUSG00000032487/ENSMUSG00000026360/ENSMUSG00000028004/ENSMUSG00000035283
GO:0007631                                                                                                                                                                                               ENSMUSG00000028004/ENSMUSG00000025905/ENSMUSG00000048482/ENSMUSG00000021647/ENSMUSG00000028341
GO:0017015                                                                                                                                                                                               ENSMUSG00000045991/ENSMUSG00000039910/ENSMUSG00000022114/ENSMUSG00000037211/ENSMUSG00000026814
GO:1902893                                                                                                                                                                                                                  ENSMUSG00000021250/ENSMUSG00000038418/ENSMUSG00000052684/ENSMUSG00000003032
GO:0032922                                                                                                                                                                                                                  ENSMUSG00000038418/ENSMUSG00000038550/ENSMUSG00000021647/ENSMUSG00000020893
GO:0035265                                                                                                                                                                            ENSMUSG00000026360/ENSMUSG00000039910/ENSMUSG00000019960/ENSMUSG00000035283/ENSMUSG00000022114/ENSMUSG00000038530
GO:1903844                                                                                                                                                                                               ENSMUSG00000045991/ENSMUSG00000039910/ENSMUSG00000022114/ENSMUSG00000037211/ENSMUSG00000026814
GO:0050805                                                                                                                                                                                                                  ENSMUSG00000022602/ENSMUSG00000032487/ENSMUSG00000028004/ENSMUSG00000048482
GO:0061614                                                                                                                                                                                                                  ENSMUSG00000021250/ENSMUSG00000038418/ENSMUSG00000052684/ENSMUSG00000003032
GO:0032891                                                                                                                                                                                                                                     ENSMUSG00000026360/ENSMUSG00000038530/ENSMUSG00000038894
GO:0045932                                                                                                                                                                                                                                     ENSMUSG00000032487/ENSMUSG00000026360/ENSMUSG00000035283
GO:0010611                                                                                                                                                                                                                  ENSMUSG00000026360/ENSMUSG00000035283/ENSMUSG00000038530/ENSMUSG00000028341
GO:0007492                                                                                                                                                                                                                  ENSMUSG00000022602/ENSMUSG00000024190/ENSMUSG00000034765/ENSMUSG00000031530
GO:0014743                                                                                                                                                                                                                  ENSMUSG00000026360/ENSMUSG00000035283/ENSMUSG00000038530/ENSMUSG00000028341
GO:0055022                                                                                                                                                                                                                                     ENSMUSG00000026360/ENSMUSG00000039910/ENSMUSG00000038530
GO:0061117                                                                                                                                                                                                                                     ENSMUSG00000026360/ENSMUSG00000039910/ENSMUSG00000038530
GO:0043154                                                                                                                                                                                                                  ENSMUSG00000023034/ENSMUSG00000032487/ENSMUSG00000003032/ENSMUSG00000090877
GO:0071277                                                                                                                                                                                                                  ENSMUSG00000021250/ENSMUSG00000052837/ENSMUSG00000003545/ENSMUSG00000052684
GO:0003206                                                                                                                                                                                               ENSMUSG00000028195/ENSMUSG00000039910/ENSMUSG00000028004/ENSMUSG00000076431/ENSMUSG00000026814
GO:0019233                                                                                                                                                                                               ENSMUSG00000032487/ENSMUSG00000028004/ENSMUSG00000035283/ENSMUSG00000025905/ENSMUSG00000024112
GO:1902074                                                                                                                                                         ENSMUSG00000021250/ENSMUSG00000052837/ENSMUSG00000003545/ENSMUSG00000038418/ENSMUSG00000052684/ENSMUSG00000025905/ENSMUSG00000048482
GO:0055021                                                                                                                                                                                                                  ENSMUSG00000026360/ENSMUSG00000039910/ENSMUSG00000035283/ENSMUSG00000038530
GO:0003231                                                                                                                                                                                               ENSMUSG00000028195/ENSMUSG00000039910/ENSMUSG00000028004/ENSMUSG00000076431/ENSMUSG00000026814
GO:0035335                                                                                                                                                                                                                                     ENSMUSG00000024190/ENSMUSG00000019960/ENSMUSG00000034765
GO:0061050                                                                                                                                                                                                                                     ENSMUSG00000026360/ENSMUSG00000035283/ENSMUSG00000038530
GO:0060411                                                                                                                                                                                                                  ENSMUSG00000028195/ENSMUSG00000039910/ENSMUSG00000076431/ENSMUSG00000026814
GO:0003209                                                                                                                                                                                                                                     ENSMUSG00000028195/ENSMUSG00000076431/ENSMUSG00000026814
GO:0010959                                                                                                                                      ENSMUSG00000032487/ENSMUSG00000035283/ENSMUSG00000025905/ENSMUSG00000038530/ENSMUSG00000019970/ENSMUSG00000075224/ENSMUSG00000024042/ENSMUSG00000020893
GO:0006942                                                                                                                                                                                                                  ENSMUSG00000023034/ENSMUSG00000026360/ENSMUSG00000035283/ENSMUSG00000024112
GO:0001558                                                                                                                                      ENSMUSG00000026360/ENSMUSG00000035283/ENSMUSG00000002083/ENSMUSG00000048482/ENSMUSG00000038530/ENSMUSG00000051243/ENSMUSG00000019970/ENSMUSG00000090877
GO:0001893                                                                                                                                                                                                                                     ENSMUSG00000052837/ENSMUSG00000032487/ENSMUSG00000039910
GO:0003151                                                                                                                                                                                                                  ENSMUSG00000039910/ENSMUSG00000028004/ENSMUSG00000052684/ENSMUSG00000026814
GO:2000628                                                                                                                                                                                                                  ENSMUSG00000021250/ENSMUSG00000038418/ENSMUSG00000052684/ENSMUSG00000003032
GO:1903320                                                                                                                                                                            ENSMUSG00000038418/ENSMUSG00000032501/ENSMUSG00000022114/ENSMUSG00000035329/ENSMUSG00000076431/ENSMUSG00000090877
GO:2000116                                                                                                                                                                            ENSMUSG00000028195/ENSMUSG00000023034/ENSMUSG00000032487/ENSMUSG00000002083/ENSMUSG00000003032/ENSMUSG00000090877
GO:0044342                                                                                                                                                                                                                                     ENSMUSG00000023034/ENSMUSG00000028341/ENSMUSG00000038894
GO:0030099                                                                                                                                      ENSMUSG00000021250/ENSMUSG00000052837/ENSMUSG00000039910/ENSMUSG00000032501/ENSMUSG00000052684/ENSMUSG00000056724/ENSMUSG00000021647/ENSMUSG00000090877
GO:0051346                                                                                                                                                         ENSMUSG00000023034/ENSMUSG00000032487/ENSMUSG00000026360/ENSMUSG00000022114/ENSMUSG00000037211/ENSMUSG00000003032/ENSMUSG00000090877
GO:0060079                                                                                                                                                                                                                  ENSMUSG00000045903/ENSMUSG00000028004/ENSMUSG00000048482/ENSMUSG00000038530
GO:0045926                                                                                                                                                                            ENSMUSG00000026360/ENSMUSG00000039910/ENSMUSG00000035283/ENSMUSG00000002083/ENSMUSG00000038530/ENSMUSG00000090877
GO:0003230                                                                                                                                                                                                                                     ENSMUSG00000028195/ENSMUSG00000076431/ENSMUSG00000026814
GO:0038179                                                                                                                                                                                                                                     ENSMUSG00000048482/ENSMUSG00000022114/ENSMUSG00000037211
GO:0048520                                                                                                                                                                                                                                     ENSMUSG00000028004/ENSMUSG00000025905/ENSMUSG00000028341
GO:1903522                                                                                                                                                                            ENSMUSG00000032487/ENSMUSG00000026360/ENSMUSG00000035283/ENSMUSG00000034765/ENSMUSG00000038530/ENSMUSG00000024112
GO:0001570                                                                                                                                                                                                                  ENSMUSG00000052837/ENSMUSG00000039910/ENSMUSG00000034640/ENSMUSG00000026814
GO:0071774                                                                                                                                                                                                                  ENSMUSG00000023034/ENSMUSG00000053560/ENSMUSG00000022114/ENSMUSG00000037211
GO:0001706                                                                                                                                                                                                                                     ENSMUSG00000024190/ENSMUSG00000034765/ENSMUSG00000031530
GO:0045823                                                                                                                                                                                                                                     ENSMUSG00000026360/ENSMUSG00000035283/ENSMUSG00000038530
GO:0002573                                                                                                                                                                            ENSMUSG00000021250/ENSMUSG00000052837/ENSMUSG00000039910/ENSMUSG00000032501/ENSMUSG00000052684/ENSMUSG00000021647
GO:2000117                                                                                                                                                                                                                  ENSMUSG00000023034/ENSMUSG00000032487/ENSMUSG00000003032/ENSMUSG00000090877
GO:0051154                                                                                                                                                                                                                                     ENSMUSG00000026360/ENSMUSG00000048482/ENSMUSG00000038530
GO:0060259                                                                                                                                                                                                                                     ENSMUSG00000028004/ENSMUSG00000025905/ENSMUSG00000028341
GO:0048738                                                                                                                                                                            ENSMUSG00000026360/ENSMUSG00000039910/ENSMUSG00000035283/ENSMUSG00000038530/ENSMUSG00000024042/ENSMUSG00000026814
GO:0043434                                                                                                                                                         ENSMUSG00000023034/ENSMUSG00000037868/ENSMUSG00000038418/ENSMUSG00000025905/ENSMUSG00000028341/ENSMUSG00000019970/ENSMUSG00000038894
GO:0048167                                                                                                                                                         ENSMUSG00000045903/ENSMUSG00000065537/ENSMUSG00000022602/ENSMUSG00000032487/ENSMUSG00000038418/ENSMUSG00000035283/ENSMUSG00000048482
GO:0010586                                                                                                                                                                                                                  ENSMUSG00000021250/ENSMUSG00000038418/ENSMUSG00000052684/ENSMUSG00000003032
GO:0030509                                                                                                                                                                                               ENSMUSG00000028195/ENSMUSG00000038418/ENSMUSG00000021765/ENSMUSG00000037573/ENSMUSG00000026814
GO:0034764                                                                                                                                                                            ENSMUSG00000022602/ENSMUSG00000035283/ENSMUSG00000025905/ENSMUSG00000028341/ENSMUSG00000075224/ENSMUSG00000038894
GO:1903532                                                                                                                                                         ENSMUSG00000028004/ENSMUSG00000035283/ENSMUSG00000025905/ENSMUSG00000021647/ENSMUSG00000024112/ENSMUSG00000076431/ENSMUSG00000038894
GO:0046621                                                                                                                                                                                                                                     ENSMUSG00000026360/ENSMUSG00000039910/ENSMUSG00000038530
GO:0033603                                                                                                                                                                                                                                                        ENSMUSG00000028004/ENSMUSG00000025905
GO:0046877                                                                                                                                                                                                                                                        ENSMUSG00000035283/ENSMUSG00000025905
GO:0060136                                                                                                                                                                                                                                                        ENSMUSG00000052837/ENSMUSG00000039910
GO:0061418                                                                                                                                                                                                                                                        ENSMUSG00000039910/ENSMUSG00000038418
GO:1902746                                                                                                                                                                                                                                                        ENSMUSG00000022114/ENSMUSG00000037211
GO:0007369                                                                                                                                                                                               ENSMUSG00000024190/ENSMUSG00000034765/ENSMUSG00000031530/ENSMUSG00000028341/ENSMUSG00000003032
GO:0050795                                                                                                                                                                                                                  ENSMUSG00000028004/ENSMUSG00000035283/ENSMUSG00000025905/ENSMUSG00000028341
GO:0001704                                                                                                                                                                                                                  ENSMUSG00000024190/ENSMUSG00000034765/ENSMUSG00000031530/ENSMUSG00000028341
GO:0030510                                                                                                                                                                                                                  ENSMUSG00000028195/ENSMUSG00000021765/ENSMUSG00000037573/ENSMUSG00000026814
GO:0071901                                                                                                                                                                                                                  ENSMUSG00000026360/ENSMUSG00000024190/ENSMUSG00000022114/ENSMUSG00000037211
GO:0097201                                                                                                                                                                                                                                                        ENSMUSG00000039910/ENSMUSG00000052684
GO:0001659                                                                                                                                                                                               ENSMUSG00000021453/ENSMUSG00000032487/ENSMUSG00000038418/ENSMUSG00000035283/ENSMUSG00000049649
GO:0048638                                                                                                                                                         ENSMUSG00000026360/ENSMUSG00000039910/ENSMUSG00000019960/ENSMUSG00000035283/ENSMUSG00000048482/ENSMUSG00000038530/ENSMUSG00000051243
GO:0003300                                                                                                                                                                                                                  ENSMUSG00000026360/ENSMUSG00000035283/ENSMUSG00000038530/ENSMUSG00000028341
GO:0003298                                                                                                                                                                                                                                     ENSMUSG00000026360/ENSMUSG00000035283/ENSMUSG00000038530
GO:0003301                                                                                                                                                                                                                                     ENSMUSG00000026360/ENSMUSG00000035283/ENSMUSG00000038530
GO:0045923                                                                                                                                                                                                                                     ENSMUSG00000032487/ENSMUSG00000028341/ENSMUSG00000038894
GO:0061049                                                                                                                                                                                                                                     ENSMUSG00000026360/ENSMUSG00000035283/ENSMUSG00000038530
GO:0071772                                                                                                                                                                                               ENSMUSG00000028195/ENSMUSG00000038418/ENSMUSG00000021765/ENSMUSG00000037573/ENSMUSG00000026814
GO:0071773                                                                                                                                                                                               ENSMUSG00000028195/ENSMUSG00000038418/ENSMUSG00000021765/ENSMUSG00000037573/ENSMUSG00000026814
GO:1901214                                                                                                                                                         ENSMUSG00000021250/ENSMUSG00000020423/ENSMUSG00000038418/ENSMUSG00000002083/ENSMUSG00000052684/ENSMUSG00000048482/ENSMUSG00000028341
GO:0043523                                                                                                                                                                            ENSMUSG00000020423/ENSMUSG00000038418/ENSMUSG00000002083/ENSMUSG00000052684/ENSMUSG00000048482/ENSMUSG00000028341
GO:1902075                                                                                                                                                                                               ENSMUSG00000021250/ENSMUSG00000052837/ENSMUSG00000003545/ENSMUSG00000038418/ENSMUSG00000052684
GO:0014897                                                                                                                                                                                                                  ENSMUSG00000026360/ENSMUSG00000035283/ENSMUSG00000038530/ENSMUSG00000028341
GO:0045639                                                                                                                                                                                                                  ENSMUSG00000021250/ENSMUSG00000032501/ENSMUSG00000052684/ENSMUSG00000090877
GO:0048661                                                                                                                                                                                                                  ENSMUSG00000032487/ENSMUSG00000038418/ENSMUSG00000052684/ENSMUSG00000028341
GO:0043405                                                                                                                                                                                               ENSMUSG00000026360/ENSMUSG00000024190/ENSMUSG00000032501/ENSMUSG00000022114/ENSMUSG00000037211
GO:0090084                                                                                                                                                                                                                                                        ENSMUSG00000005483/ENSMUSG00000090877
GO:2000253                                                                                                                                                                                                                                                        ENSMUSG00000025905/ENSMUSG00000028341
GO:2000322                                                                                                                                                                                                                                                        ENSMUSG00000048482/ENSMUSG00000020893
GO:0055017                                                                                                                                                                                                                  ENSMUSG00000026360/ENSMUSG00000039910/ENSMUSG00000035283/ENSMUSG00000038530
GO:0003205                                                                                                                                                                                               ENSMUSG00000028195/ENSMUSG00000039910/ENSMUSG00000028004/ENSMUSG00000076431/ENSMUSG00000026814
GO:0009749                                                                                                                                                                                               ENSMUSG00000038418/ENSMUSG00000025905/ENSMUSG00000035828/ENSMUSG00000076431/ENSMUSG00000038894
GO:0060135                                                                                                                                                                                                                                     ENSMUSG00000052837/ENSMUSG00000032487/ENSMUSG00000039910
GO:0014896                                                                                                                                                                                                                  ENSMUSG00000026360/ENSMUSG00000035283/ENSMUSG00000038530/ENSMUSG00000028341
GO:0009746                                                                                                                                                                                               ENSMUSG00000038418/ENSMUSG00000025905/ENSMUSG00000035828/ENSMUSG00000076431/ENSMUSG00000038894
GO:0006939                                                                                                                                                                                                                  ENSMUSG00000032487/ENSMUSG00000026360/ENSMUSG00000028004/ENSMUSG00000035283
GO:0055006                                                                                                                                                                                                                  ENSMUSG00000026360/ENSMUSG00000035283/ENSMUSG00000038530/ENSMUSG00000037211
GO:0034284                                                                                                                                                                                               ENSMUSG00000038418/ENSMUSG00000025905/ENSMUSG00000035828/ENSMUSG00000076431/ENSMUSG00000038894
GO:0031953                                                                                                                                                                                                                                                        ENSMUSG00000052684/ENSMUSG00000026814
GO:0060213                                                                                                                                                                                                                                                        ENSMUSG00000020423/ENSMUSG00000037573
GO:0060840                                                                                                                                                                                                                  ENSMUSG00000037868/ENSMUSG00000039910/ENSMUSG00000076431/ENSMUSG00000026814
GO:0002042                                                                                                                                                                                                                                     ENSMUSG00000023034/ENSMUSG00000032487/ENSMUSG00000003032
GO:0045933                                                                                                                                                                                                                                     ENSMUSG00000032487/ENSMUSG00000026360/ENSMUSG00000028004
GO:0006936                                                                                                                                                                            ENSMUSG00000023034/ENSMUSG00000032487/ENSMUSG00000026360/ENSMUSG00000028004/ENSMUSG00000035283/ENSMUSG00000024112
GO:0051954                                                                                                                                                                                                                                     ENSMUSG00000028004/ENSMUSG00000025905/ENSMUSG00000021647
GO:0031396                                                                                                                                                                                               ENSMUSG00000032501/ENSMUSG00000022114/ENSMUSG00000035329/ENSMUSG00000076431/ENSMUSG00000090877
GO:0030857                                                                                                                                                                                                                                     ENSMUSG00000021765/ENSMUSG00000022114/ENSMUSG00000037211
GO:0045777                                                                                                                                                                                                                                     ENSMUSG00000035283/ENSMUSG00000021647/ENSMUSG00000026814
GO:0009314                                                                                                                                                         ENSMUSG00000038418/ENSMUSG00000028702/ENSMUSG00000002083/ENSMUSG00000052684/ENSMUSG00000019970/ENSMUSG00000024042/ENSMUSG00000020893
GO:1901652                                                                                                                                                         ENSMUSG00000023034/ENSMUSG00000037868/ENSMUSG00000038418/ENSMUSG00000025905/ENSMUSG00000028341/ENSMUSG00000019970/ENSMUSG00000038894
GO:0003279                                                                                                                                                                                                                  ENSMUSG00000028195/ENSMUSG00000039910/ENSMUSG00000076431/ENSMUSG00000026814
GO:0048640                                                                                                                                                                                                                  ENSMUSG00000026360/ENSMUSG00000039910/ENSMUSG00000035283/ENSMUSG00000038530
GO:0009743                                                                                                                                                                                               ENSMUSG00000038418/ENSMUSG00000025905/ENSMUSG00000035828/ENSMUSG00000076431/ENSMUSG00000038894
GO:0015816                                                                                                                                                                                                                                                        ENSMUSG00000026360/ENSMUSG00000038530
GO:0060211                                                                                                                                                                                                                                                        ENSMUSG00000020423/ENSMUSG00000037573
GO:0061052                                                                                                                                                                                                                                                        ENSMUSG00000026360/ENSMUSG00000038530
GO:0042752                                                                                                                                                                                                                  ENSMUSG00000028004/ENSMUSG00000035283/ENSMUSG00000024042/ENSMUSG00000020893
GO:0051047                                                                                                                                                         ENSMUSG00000028004/ENSMUSG00000035283/ENSMUSG00000025905/ENSMUSG00000021647/ENSMUSG00000024112/ENSMUSG00000076431/ENSMUSG00000038894
GO:1901216                                                                                                                                                                                                                  ENSMUSG00000021250/ENSMUSG00000038418/ENSMUSG00000002083/ENSMUSG00000052684
GO:0042692                                                                                                                                                         ENSMUSG00000026360/ENSMUSG00000035283/ENSMUSG00000044471/ENSMUSG00000048482/ENSMUSG00000038530/ENSMUSG00000024042/ENSMUSG00000026814
GO:0051402                                                                                                                                                                            ENSMUSG00000020423/ENSMUSG00000038418/ENSMUSG00000002083/ENSMUSG00000052684/ENSMUSG00000048482/ENSMUSG00000028341
GO:0007406                                                                                                                                                                                                                                                        ENSMUSG00000020423/ENSMUSG00000048482
GO:0035970                                                                                                                                                                                                                                                        ENSMUSG00000024190/ENSMUSG00000034765
GO:0042921                                                                                                                                                                                                                                                        ENSMUSG00000048482/ENSMUSG00000020893
GO:0046541                                                                                                                                                                                                                                                        ENSMUSG00000035283/ENSMUSG00000025905
GO:0042596                                                                                                                                                                                                                                     ENSMUSG00000028004/ENSMUSG00000035283/ENSMUSG00000048482
GO:0070997                                                                                                                                                         ENSMUSG00000021250/ENSMUSG00000020423/ENSMUSG00000038418/ENSMUSG00000002083/ENSMUSG00000052684/ENSMUSG00000048482/ENSMUSG00000028341
GO:0051146                                                                                                                                                                            ENSMUSG00000026360/ENSMUSG00000035283/ENSMUSG00000044471/ENSMUSG00000048482/ENSMUSG00000038530/ENSMUSG00000024042
GO:0042303                                                                                                                                                                                                                  ENSMUSG00000032487/ENSMUSG00000021765/ENSMUSG00000044471/ENSMUSG00000020893
GO:0042633                                                                                                                                                                                                                  ENSMUSG00000032487/ENSMUSG00000021765/ENSMUSG00000044471/ENSMUSG00000020893
GO:0051592                                                                                                                                                                                                                  ENSMUSG00000021250/ENSMUSG00000052837/ENSMUSG00000003545/ENSMUSG00000052684
GO:0045637                                                                                                                                                                                               ENSMUSG00000021250/ENSMUSG00000032501/ENSMUSG00000052684/ENSMUSG00000021647/ENSMUSG00000090877
GO:0050873                                                                                                                                                                                                                                     ENSMUSG00000032487/ENSMUSG00000026360/ENSMUSG00000035283
GO:0055001                                                                                                                                                                                               ENSMUSG00000026360/ENSMUSG00000035283/ENSMUSG00000044471/ENSMUSG00000038530/ENSMUSG00000026814
GO:0031958                                                                                                                                                                                                                                                        ENSMUSG00000048482/ENSMUSG00000020893
GO:0051386                                                                                                                                                                                                                                                        ENSMUSG00000022114/ENSMUSG00000037211
GO:0060413                                                                                                                                                                                                                                                        ENSMUSG00000028195/ENSMUSG00000076431
GO:0071498                                                                                                                                                                                                                                                        ENSMUSG00000032487/ENSMUSG00000003032
GO:0050679                                                                                                                                                                                               ENSMUSG00000023034/ENSMUSG00000052684/ENSMUSG00000037169/ENSMUSG00000028341/ENSMUSG00000038894
GO:0051403                                                                                                                                                                                               ENSMUSG00000021453/ENSMUSG00000024190/ENSMUSG00000032501/ENSMUSG00000025905/ENSMUSG00000020893
GO:0051148                                                                                                                                                                                                                                     ENSMUSG00000026360/ENSMUSG00000048482/ENSMUSG00000038530
GO:0042180                                                                                                                                                                                               ENSMUSG00000032487/ENSMUSG00000038418/ENSMUSG00000028341/ENSMUSG00000024112/ENSMUSG00000038894
GO:0050433                                                                                                                                                                                                                                     ENSMUSG00000028004/ENSMUSG00000025905/ENSMUSG00000021647
GO:0006470                                                                                                                                                                                               ENSMUSG00000024190/ENSMUSG00000019960/ENSMUSG00000034765/ENSMUSG00000031530/ENSMUSG00000018648
GO:0071241                                                                                                                                                                                               ENSMUSG00000065537/ENSMUSG00000021250/ENSMUSG00000052837/ENSMUSG00000003545/ENSMUSG00000052684
GO:0042391                                                                                                                                                         ENSMUSG00000045903/ENSMUSG00000028004/ENSMUSG00000035283/ENSMUSG00000052684/ENSMUSG00000048482/ENSMUSG00000038530/ENSMUSG00000024112
GO:0040037                                                                                                                                                                                                                                                        ENSMUSG00000022114/ENSMUSG00000037211
GO:0051385                                                                                                                                                                                                                                                        ENSMUSG00000045903/ENSMUSG00000019970
GO:0002761                                                                                                                                                                                                                  ENSMUSG00000021250/ENSMUSG00000032501/ENSMUSG00000052684/ENSMUSG00000021647
GO:0007389                                                                                                                                                         ENSMUSG00000022602/ENSMUSG00000037868/ENSMUSG00000020423/ENSMUSG00000039910/ENSMUSG00000021765/ENSMUSG00000037211/ENSMUSG00000026814
GO:0055117                                                                                                                                                                                                                                     ENSMUSG00000026360/ENSMUSG00000035283/ENSMUSG00000024112
GO:2000112                                                                                                                                                         ENSMUSG00000020423/ENSMUSG00000026360/ENSMUSG00000037573/ENSMUSG00000003032/ENSMUSG00000076431/ENSMUSG00000020893/ENSMUSG00000038894
GO:0052548                                                                                                                                                                            ENSMUSG00000028195/ENSMUSG00000023034/ENSMUSG00000032487/ENSMUSG00000002083/ENSMUSG00000003032/ENSMUSG00000090877
GO:0007193                                                                                                                                                                                                                                     ENSMUSG00000026360/ENSMUSG00000028004/ENSMUSG00000025905
GO:0048148                                                                                                                                                                                                                                                        ENSMUSG00000025905/ENSMUSG00000048482
GO:0051956                                                                                                                                                                                                                                                        ENSMUSG00000026360/ENSMUSG00000038530
GO:0002763                                                                                                                                                                                                                                     ENSMUSG00000021250/ENSMUSG00000032501/ENSMUSG00000052684
GO:0035904                                                                                                                                                                                                                                     ENSMUSG00000037868/ENSMUSG00000076431/ENSMUSG00000026814
GO:0031098                                                                                                                                                                                               ENSMUSG00000021453/ENSMUSG00000024190/ENSMUSG00000032501/ENSMUSG00000025905/ENSMUSG00000020893
GO:0048168                                                                                                                                                                                                                                     ENSMUSG00000022602/ENSMUSG00000038418/ENSMUSG00000048482
GO:0055007                                                                                                                                                                                                                  ENSMUSG00000026360/ENSMUSG00000035283/ENSMUSG00000038530/ENSMUSG00000024042
GO:0006700                                                                                                                                                                                                                                                        ENSMUSG00000038418/ENSMUSG00000024112
GO:0010958                                                                                                                                                                                                                                                        ENSMUSG00000026360/ENSMUSG00000038530
GO:0035994                                                                                                                                                                                                                                                        ENSMUSG00000021250/ENSMUSG00000052684
GO:0090083                                                                                                                                                                                                                                                        ENSMUSG00000005483/ENSMUSG00000090877
GO:1903789                                                                                                                                                                                                                                                        ENSMUSG00000026360/ENSMUSG00000038530
GO:0007422                                                                                                                                                                                                                                     ENSMUSG00000037868/ENSMUSG00000045991/ENSMUSG00000039910
GO:0050432                                                                                                                                                                                                                                     ENSMUSG00000028004/ENSMUSG00000025905/ENSMUSG00000021647
GO:0032868                                                                                                                                                                                               ENSMUSG00000037868/ENSMUSG00000038418/ENSMUSG00000025905/ENSMUSG00000019970/ENSMUSG00000038894
GO:0098773                                                                                                                                                                                                                  ENSMUSG00000032487/ENSMUSG00000021765/ENSMUSG00000044471/ENSMUSG00000003032
GO:0010565                                                                                                                                                                                                                  ENSMUSG00000032487/ENSMUSG00000038418/ENSMUSG00000028341/ENSMUSG00000038894
GO:0045475                                                                                                                                                                                                                                                        ENSMUSG00000038418/ENSMUSG00000038550
GO:0046321                                                                                                                                                                                                                                                        ENSMUSG00000028341/ENSMUSG00000038894
GO:0060602                                                                                                                                                                                                                                                        ENSMUSG00000022114/ENSMUSG00000037211
GO:0008544                                                                                                                                                                            ENSMUSG00000032487/ENSMUSG00000021765/ENSMUSG00000044471/ENSMUSG00000037169/ENSMUSG00000029135/ENSMUSG00000003032
GO:0031649                                                                                                                                                                                                                                                        ENSMUSG00000032487/ENSMUSG00000035283
GO:0071276                                                                                                                                                                                                                                                        ENSMUSG00000021250/ENSMUSG00000052684
GO:0120255                                                                                                                                                                                                                                                        ENSMUSG00000038418/ENSMUSG00000024112
GO:0002791                                                                                                                                                                                               ENSMUSG00000028004/ENSMUSG00000021647/ENSMUSG00000035828/ENSMUSG00000076431/ENSMUSG00000038894
GO:0019216                                                                                                                                                                            ENSMUSG00000028195/ENSMUSG00000032487/ENSMUSG00000038418/ENSMUSG00000028341/ENSMUSG00000024042/ENSMUSG00000038894
GO:0030856                                                                                                                                                                                                                  ENSMUSG00000021765/ENSMUSG00000037169/ENSMUSG00000022114/ENSMUSG00000037211
GO:0090087                                                                                                                                                                                               ENSMUSG00000028004/ENSMUSG00000021647/ENSMUSG00000035828/ENSMUSG00000076431/ENSMUSG00000038894
GO:0043270                                                                                                                                                                                               ENSMUSG00000022602/ENSMUSG00000035283/ENSMUSG00000025905/ENSMUSG00000019970/ENSMUSG00000075224
GO:0060562                                                                                                                                                                            ENSMUSG00000039910/ENSMUSG00000037169/ENSMUSG00000022114/ENSMUSG00000037211/ENSMUSG00000076431/ENSMUSG00000026814
GO:0003283                                                                                                                                                                                                                                                        ENSMUSG00000028195/ENSMUSG00000076431
GO:0003208                                                                                                                                                                                                                                     ENSMUSG00000028004/ENSMUSG00000076431/ENSMUSG00000026814
GO:0048844                                                                                                                                                                                                                                     ENSMUSG00000039910/ENSMUSG00000076431/ENSMUSG00000026814
GO:0071383                                                                                                                                                                                                                  ENSMUSG00000045903/ENSMUSG00000048482/ENSMUSG00000019970/ENSMUSG00000020893
GO:0001829                                                                                                                                                                                                                                                        ENSMUSG00000052837/ENSMUSG00000039910
GO:0043153                                                                                                                                                                                                                                                        ENSMUSG00000024042/ENSMUSG00000020893
GO:0006941                                                                                                                                                                                                                  ENSMUSG00000023034/ENSMUSG00000026360/ENSMUSG00000035283/ENSMUSG00000024112
GO:0031397                                                                                                                                                                                                                                     ENSMUSG00000022114/ENSMUSG00000076431/ENSMUSG00000090877
GO:0001678                                                                                                                                                                                                                  ENSMUSG00000025905/ENSMUSG00000021647/ENSMUSG00000035828/ENSMUSG00000076431
GO:0071248                                                                                                                                                                                                                  ENSMUSG00000021250/ENSMUSG00000052837/ENSMUSG00000003545/ENSMUSG00000052684
GO:0003084                                                                                                                                                                                                                                                        ENSMUSG00000035283/ENSMUSG00000026814
GO:0003181                                                                                                                                                                                                                                                        ENSMUSG00000028195/ENSMUSG00000076431
GO:0009648                                                                                                                                                                                                                                                        ENSMUSG00000024042/ENSMUSG00000020893
GO:0048588                                                                                                                                                                                               ENSMUSG00000026360/ENSMUSG00000035283/ENSMUSG00000048482/ENSMUSG00000038530/ENSMUSG00000051243
GO:0043525                                                                                                                                                                                                                                     ENSMUSG00000038418/ENSMUSG00000002083/ENSMUSG00000052684
GO:0035296                                                                                                                                                                                                                  ENSMUSG00000032487/ENSMUSG00000026360/ENSMUSG00000035283/ENSMUSG00000034765
GO:0097746                                                                                                                                                                                                                  ENSMUSG00000032487/ENSMUSG00000026360/ENSMUSG00000035283/ENSMUSG00000034765
GO:0003281                                                                                                                                                                                                                                     ENSMUSG00000028195/ENSMUSG00000039910/ENSMUSG00000076431
GO:0000289                                                                                                                                                                                                                                                        ENSMUSG00000020423/ENSMUSG00000037573
GO:0009649                                                                                                                                                                                                                                                        ENSMUSG00000024042/ENSMUSG00000020893
GO:0060716                                                                                                                                                                                                                                                        ENSMUSG00000028195/ENSMUSG00000052837
GO:0072567                                                                                                                                                                                                                                                        ENSMUSG00000028195/ENSMUSG00000003032
GO:2000341                                                                                                                                                                                                                                                        ENSMUSG00000028195/ENSMUSG00000003032
GO:0035150                                                                                                                                                                                                                  ENSMUSG00000032487/ENSMUSG00000026360/ENSMUSG00000035283/ENSMUSG00000034765
GO:0031214                                                                                                                                                                                                                  ENSMUSG00000028195/ENSMUSG00000032487/ENSMUSG00000046761/ENSMUSG00000044471
GO:0002088                                                                                                                                                                                                                                     ENSMUSG00000039910/ENSMUSG00000022114/ENSMUSG00000037211
GO:1900745                                                                                                                                                                                                                                                        ENSMUSG00000021453/ENSMUSG00000025905
GO:0001890                                                                                                                                                                                                                  ENSMUSG00000028195/ENSMUSG00000052837/ENSMUSG00000032487/ENSMUSG00000039910
GO:0050678                                                                                                                                                                            ENSMUSG00000023034/ENSMUSG00000052684/ENSMUSG00000037169/ENSMUSG00000028341/ENSMUSG00000038894/ENSMUSG00000026814
GO:0001503                                                                                                                                                                            ENSMUSG00000028195/ENSMUSG00000037868/ENSMUSG00000052837/ENSMUSG00000032487/ENSMUSG00000044471/ENSMUSG00000037573
GO:0045834                                                                                                                                                                                                                  ENSMUSG00000028195/ENSMUSG00000032487/ENSMUSG00000028341/ENSMUSG00000038894
GO:0001654                                                                                                                                                                            ENSMUSG00000039910/ENSMUSG00000052684/ENSMUSG00000048482/ENSMUSG00000022114/ENSMUSG00000037211/ENSMUSG00000003032
GO:0021675                                                                                                                                                                                                                                     ENSMUSG00000037868/ENSMUSG00000039910/ENSMUSG00000048482
GO:0030512                                                                                                                                                                                                                                     ENSMUSG00000045991/ENSMUSG00000022114/ENSMUSG00000037211
GO:0051937                                                                                                                                                                                                                                     ENSMUSG00000028004/ENSMUSG00000025905/ENSMUSG00000021647
GO:0150063                                                                                                                                                                            ENSMUSG00000039910/ENSMUSG00000052684/ENSMUSG00000048482/ENSMUSG00000022114/ENSMUSG00000037211/ENSMUSG00000003032
GO:0045187                                                                                                                                                                                                                                                        ENSMUSG00000028004/ENSMUSG00000035283
GO:0070841                                                                                                                                                                                                                                                        ENSMUSG00000005483/ENSMUSG00000090877
GO:0042886                                                                                                                                                                            ENSMUSG00000028004/ENSMUSG00000048482/ENSMUSG00000021647/ENSMUSG00000035828/ENSMUSG00000076431/ENSMUSG00000038894
GO:0048880                                                                                                                                                                            ENSMUSG00000039910/ENSMUSG00000052684/ENSMUSG00000048482/ENSMUSG00000022114/ENSMUSG00000037211/ENSMUSG00000003032
GO:0044344                                                                                                                                                                                                                                     ENSMUSG00000023034/ENSMUSG00000022114/ENSMUSG00000037211
GO:1901379                                                                                                                                                                                                                                     ENSMUSG00000025905/ENSMUSG00000038530/ENSMUSG00000075224
GO:0003171                                                                                                                                                                                                                                                        ENSMUSG00000028195/ENSMUSG00000076431
GO:0010460                                                                                                                                                                                                                                                        ENSMUSG00000035283/ENSMUSG00000038530
GO:0048745                                                                                                                                                                                                                                                        ENSMUSG00000034640/ENSMUSG00000026814
GO:0030534                                                                                                                                                                                                                  ENSMUSG00000025905/ENSMUSG00000048482/ENSMUSG00000021647/ENSMUSG00000028341
GO:0048754                                                                                                                                                                                                                  ENSMUSG00000037169/ENSMUSG00000022114/ENSMUSG00000037211/ENSMUSG00000026814
GO:0010951                                                                                                                                                                                                                  ENSMUSG00000023034/ENSMUSG00000032487/ENSMUSG00000003032/ENSMUSG00000090877
GO:1903321                                                                                                                                                                                                                                     ENSMUSG00000022114/ENSMUSG00000076431/ENSMUSG00000090877
GO:1903706                                                                                                                                                                            ENSMUSG00000021250/ENSMUSG00000032501/ENSMUSG00000052684/ENSMUSG00000021647/ENSMUSG00000076431/ENSMUSG00000090877
GO:0021602                                                                                                                                                                                                                                                        ENSMUSG00000037868/ENSMUSG00000039910
GO:0042634                                                                                                                                                                                                                                                        ENSMUSG00000021765/ENSMUSG00000020893
GO:0050802                                                                                                                                                                                                                                                        ENSMUSG00000028004/ENSMUSG00000035283
GO:0010717                                                                                                                                                                                                                                     ENSMUSG00000022114/ENSMUSG00000037211/ENSMUSG00000026814
GO:0046888                                                                                                                                                                                                                                     ENSMUSG00000025905/ENSMUSG00000021647/ENSMUSG00000035828
GO:0070482                                                                                                                                                                                               ENSMUSG00000065537/ENSMUSG00000039910/ENSMUSG00000038418/ENSMUSG00000029135/ENSMUSG00000026814
GO:0120162                                                                                                                                                                                                                                     ENSMUSG00000021453/ENSMUSG00000035283/ENSMUSG00000049649
GO:0030336                                                                                                                                                                                               ENSMUSG00000039910/ENSMUSG00000024190/ENSMUSG00000032501/ENSMUSG00000003032/ENSMUSG00000026814
GO:0042749                                                                                                                                                                                                                                                        ENSMUSG00000028004/ENSMUSG00000035283
GO:0048011                                                                                                                                                                                                                                                        ENSMUSG00000022114/ENSMUSG00000037211
GO:0060317                                                                                                                                                                                                                                                        ENSMUSG00000037211/ENSMUSG00000026814
GO:0071875                                                                                                                                                                                                                                                        ENSMUSG00000026360/ENSMUSG00000035283
GO:2000178                                                                                                                                                                                                                                                        ENSMUSG00000020423/ENSMUSG00000048482
GO:0010721                                                                                                                                                                                               ENSMUSG00000020423/ENSMUSG00000032501/ENSMUSG00000048482/ENSMUSG00000037169/ENSMUSG00000021647
GO:0003002                                                                                                                                                                            ENSMUSG00000022602/ENSMUSG00000037868/ENSMUSG00000020423/ENSMUSG00000039910/ENSMUSG00000037211/ENSMUSG00000026814
GO:0043524                                                                                                                                                                                                                  ENSMUSG00000020423/ENSMUSG00000052684/ENSMUSG00000048482/ENSMUSG00000028341
GO:0014910                                                                                                                                                                                                                                     ENSMUSG00000038418/ENSMUSG00000032501/ENSMUSG00000028341
GO:0008016                                                                                                                                                                                                                  ENSMUSG00000026360/ENSMUSG00000035283/ENSMUSG00000038530/ENSMUSG00000024112
GO:0002028                                                                                                                                                                                                                                     ENSMUSG00000019970/ENSMUSG00000024042/ENSMUSG00000020893
GO:0032651                                                                                                                                                                                                                                     ENSMUSG00000028195/ENSMUSG00000031103/ENSMUSG00000038418
GO:0060395                                                                                                                                                                                                                                     ENSMUSG00000021250/ENSMUSG00000052684/ENSMUSG00000037573
GO:0002053                                                                                                                                                                                                                                                        ENSMUSG00000037169/ENSMUSG00000038894
GO:0034260                                                                                                                                                                                                                                                        ENSMUSG00000022114/ENSMUSG00000037211
GO:0040036                                                                                                                                                                                                                                                        ENSMUSG00000022114/ENSMUSG00000037211
GO:0051085                                                                                                                                                                                                                                                        ENSMUSG00000005483/ENSMUSG00000090877
GO:2000108                                                                                                                                                                                                                                                        ENSMUSG00000002083/ENSMUSG00000028341
GO:0032872                                                                                                                                                                                                                  ENSMUSG00000021453/ENSMUSG00000024190/ENSMUSG00000025905/ENSMUSG00000020893
GO:0090288                                                                                                                                                                                                                                     ENSMUSG00000022114/ENSMUSG00000037573/ENSMUSG00000037211
GO:0003156                                                                                                                                                                                                                                                        ENSMUSG00000039910/ENSMUSG00000037211
GO:0008340                                                                                                                                                                                                                                                        ENSMUSG00000028702/ENSMUSG00000002083
GO:0022410                                                                                                                                                                                                                                                        ENSMUSG00000028004/ENSMUSG00000035283
GO:0070302                                                                                                                                                                                                                  ENSMUSG00000021453/ENSMUSG00000024190/ENSMUSG00000025905/ENSMUSG00000020893
GO:0034767                                                                                                                                                                                                                  ENSMUSG00000022602/ENSMUSG00000035283/ENSMUSG00000025905/ENSMUSG00000075224
GO:0010631                                                                                                                                                                                               ENSMUSG00000023034/ENSMUSG00000032487/ENSMUSG00000052684/ENSMUSG00000003032/ENSMUSG00000038894
GO:2000146                                                                                                                                                                                               ENSMUSG00000039910/ENSMUSG00000024190/ENSMUSG00000032501/ENSMUSG00000003032/ENSMUSG00000026814
GO:0090132                                                                                                                                                                                               ENSMUSG00000023034/ENSMUSG00000032487/ENSMUSG00000052684/ENSMUSG00000003032/ENSMUSG00000038894
GO:0030308                                                                                                                                                                                                                  ENSMUSG00000026360/ENSMUSG00000002083/ENSMUSG00000038530/ENSMUSG00000090877
GO:0052547                                                                                                                                                                            ENSMUSG00000028195/ENSMUSG00000023034/ENSMUSG00000032487/ENSMUSG00000002083/ENSMUSG00000003032/ENSMUSG00000090877
GO:0010614                                                                                                                                                                                                                                                        ENSMUSG00000026360/ENSMUSG00000038530
GO:0045987                                                                                                                                                                                                                                                        ENSMUSG00000032487/ENSMUSG00000028004
GO:0002790                                                                                                                                                                                               ENSMUSG00000028004/ENSMUSG00000021647/ENSMUSG00000035828/ENSMUSG00000076431/ENSMUSG00000038894
GO:0019217                                                                                                                                                                                                                                     ENSMUSG00000032487/ENSMUSG00000028341/ENSMUSG00000038894
GO:0090130                                                                                                                                                                                               ENSMUSG00000023034/ENSMUSG00000032487/ENSMUSG00000052684/ENSMUSG00000003032/ENSMUSG00000038894
GO:0043588                                                                                                                                                                                               ENSMUSG00000032487/ENSMUSG00000021765/ENSMUSG00000044471/ENSMUSG00000029135/ENSMUSG00000003032
GO:0030307                                                                                                                                                                                                                  ENSMUSG00000035283/ENSMUSG00000048482/ENSMUSG00000051243/ENSMUSG00000019970
GO:0098739                                                                                                                                                                                                                  ENSMUSG00000026360/ENSMUSG00000035283/ENSMUSG00000038530/ENSMUSG00000038894
GO:1902107                                                                                                                                                                                                                  ENSMUSG00000021250/ENSMUSG00000032501/ENSMUSG00000052684/ENSMUSG00000076431
GO:1903708                                                                                                                                                                                                                  ENSMUSG00000021250/ENSMUSG00000032501/ENSMUSG00000052684/ENSMUSG00000076431
GO:0032611                                                                                                                                                                                                                                     ENSMUSG00000028195/ENSMUSG00000031103/ENSMUSG00000038418
GO:0046890                                                                                                                                                                                                                  ENSMUSG00000028195/ENSMUSG00000032487/ENSMUSG00000038418/ENSMUSG00000024042
GO:0000132                                                                                                                                                                                                                                                        ENSMUSG00000022114/ENSMUSG00000037211
GO:0010719                                                                                                                                                                                                                                                        ENSMUSG00000022114/ENSMUSG00000037211
GO:0046686                                                                                                                                                                                                                                                        ENSMUSG00000021250/ENSMUSG00000052684
GO:0034248                                                                                                                                                                            ENSMUSG00000028195/ENSMUSG00000020423/ENSMUSG00000026360/ENSMUSG00000037573/ENSMUSG00000076431/ENSMUSG00000020893
GO:0032890                                                                                                                                                                                                                                     ENSMUSG00000026360/ENSMUSG00000038530/ENSMUSG00000038894
GO:0055013                                                                                                                                                                                                                                     ENSMUSG00000026360/ENSMUSG00000035283/ENSMUSG00000038530
GO:0048562                                                                                                                                                                                               ENSMUSG00000039910/ENSMUSG00000037169/ENSMUSG00000022114/ENSMUSG00000028341/ENSMUSG00000026814
GO:0001822                                                                                                                                                                                               ENSMUSG00000038418/ENSMUSG00000048482/ENSMUSG00000034640/ENSMUSG00000037211/ENSMUSG00000076431
           Count
GO:0060537    15
GO:0043409    10
GO:0070373     7
GO:0007611    11
GO:0035914     7
GO:0050890    11
GO:0007623     9
GO:0007178    11
GO:0007612     8
GO:0090257     9
GO:0007519     8
GO:0007517    10
GO:0060538     8
GO:0007613     7
GO:0042326    10
GO:0048511     9
GO:0070372     9
GO:0007622     5
GO:0070371     9
GO:0001933     9
GO:0010563    10
GO:0045936    10
GO:0090092     8
GO:0007179     7
GO:1900744     4
GO:0043618     4
GO:0090101     6
GO:0007616     4
GO:0060420     5
GO:0033673     7
GO:0090287     8
GO:1903524     4
GO:0043620     4
GO:0051147     6
GO:0003012     9
GO:0071560     7
GO:0071559     7
GO:0006937     6
GO:0038066     4
GO:0043502     5
GO:1902895     4
GO:0033605     3
GO:0051153     5
GO:0045986     3
GO:0061469     3
GO:0045444     7
GO:0051348     7
GO:0033002     7
GO:2000630     4
GO:0099565     5
GO:0048660     6
GO:0043407     4
GO:0035051     6
GO:0008306     5
GO:0034405     3
GO:0060080     3
GO:0048659     6
GO:0051952     5
GO:0060419     5
GO:0046620     5
GO:0003007     7
GO:0048512     4
GO:0046697     3
GO:0031644     6
GO:0014706     7
GO:0006469     6
GO:0043500     5
GO:0015837     5
GO:0043281     6
GO:0060078     5
GO:0006940     4
GO:0007631     5
GO:0017015     5
GO:1902893     4
GO:0032922     4
GO:0035265     6
GO:1903844     5
GO:0050805     4
GO:0061614     4
GO:0032891     3
GO:0045932     3
GO:0010611     4
GO:0007492     4
GO:0014743     4
GO:0055022     3
GO:0061117     3
GO:0043154     4
GO:0071277     4
GO:0003206     5
GO:0019233     5
GO:1902074     7
GO:0055021     4
GO:0003231     5
GO:0035335     3
GO:0061050     3
GO:0060411     4
GO:0003209     3
GO:0010959     8
GO:0006942     4
GO:0001558     8
GO:0001893     3
GO:0003151     4
GO:2000628     4
GO:1903320     6
GO:2000116     6
GO:0044342     3
GO:0030099     8
GO:0051346     7
GO:0060079     4
GO:0045926     6
GO:0003230     3
GO:0038179     3
GO:0048520     3
GO:1903522     6
GO:0001570     4
GO:0071774     4
GO:0001706     3
GO:0045823     3
GO:0002573     6
GO:2000117     4
GO:0051154     3
GO:0060259     3
GO:0048738     6
GO:0043434     7
GO:0048167     7
GO:0010586     4
GO:0030509     5
GO:0034764     6
GO:1903532     7
GO:0046621     3
GO:0033603     2
GO:0046877     2
GO:0060136     2
GO:0061418     2
GO:1902746     2
GO:0007369     5
GO:0050795     4
GO:0001704     4
GO:0030510     4
GO:0071901     4
GO:0097201     2
GO:0001659     5
GO:0048638     7
GO:0003300     4
GO:0003298     3
GO:0003301     3
GO:0045923     3
GO:0061049     3
GO:0071772     5
GO:0071773     5
GO:1901214     7
GO:0043523     6
GO:1902075     5
GO:0014897     4
GO:0045639     4
GO:0048661     4
GO:0043405     5
GO:0090084     2
GO:2000253     2
GO:2000322     2
GO:0055017     4
GO:0003205     5
GO:0009749     5
GO:0060135     3
GO:0014896     4
GO:0009746     5
GO:0006939     4
GO:0055006     4
GO:0034284     5
GO:0031953     2
GO:0060213     2
GO:0060840     4
GO:0002042     3
GO:0045933     3
GO:0006936     6
GO:0051954     3
GO:0031396     5
GO:0030857     3
GO:0045777     3
GO:0009314     7
GO:1901652     7
GO:0003279     4
GO:0048640     4
GO:0009743     5
GO:0015816     2
GO:0060211     2
GO:0061052     2
GO:0042752     4
GO:0051047     7
GO:1901216     4
GO:0042692     7
GO:0051402     6
GO:0007406     2
GO:0035970     2
GO:0042921     2
GO:0046541     2
GO:0042596     3
GO:0070997     7
GO:0051146     6
GO:0042303     4
GO:0042633     4
GO:0051592     4
GO:0045637     5
GO:0050873     3
GO:0055001     5
GO:0031958     2
GO:0051386     2
GO:0060413     2
GO:0071498     2
GO:0050679     5
GO:0051403     5
GO:0051148     3
GO:0042180     5
GO:0050433     3
GO:0006470     5
GO:0071241     5
GO:0042391     7
GO:0040037     2
GO:0051385     2
GO:0002761     4
GO:0007389     7
GO:0055117     3
GO:2000112     7
GO:0052548     6
GO:0007193     3
GO:0048148     2
GO:0051956     2
GO:0002763     3
GO:0035904     3
GO:0031098     5
GO:0048168     3
GO:0055007     4
GO:0006700     2
GO:0010958     2
GO:0035994     2
GO:0090083     2
GO:1903789     2
GO:0007422     3
GO:0050432     3
GO:0032868     5
GO:0098773     4
GO:0010565     4
GO:0045475     2
GO:0046321     2
GO:0060602     2
GO:0008544     6
GO:0031649     2
GO:0071276     2
GO:0120255     2
GO:0002791     5
GO:0019216     6
GO:0030856     4
GO:0090087     5
GO:0043270     5
GO:0060562     6
GO:0003283     2
GO:0003208     3
GO:0048844     3
GO:0071383     4
GO:0001829     2
GO:0043153     2
GO:0006941     4
GO:0031397     3
GO:0001678     4
GO:0071248     4
GO:0003084     2
GO:0003181     2
GO:0009648     2
GO:0048588     5
GO:0043525     3
GO:0035296     4
GO:0097746     4
GO:0003281     3
GO:0000289     2
GO:0009649     2
GO:0060716     2
GO:0072567     2
GO:2000341     2
GO:0035150     4
GO:0031214     4
GO:0002088     3
GO:1900745     2
GO:0001890     4
GO:0050678     6
GO:0001503     6
GO:0045834     4
GO:0001654     6
GO:0021675     3
GO:0030512     3
GO:0051937     3
GO:0150063     6
GO:0045187     2
GO:0070841     2
GO:0042886     6
GO:0048880     6
GO:0044344     3
GO:1901379     3
GO:0003171     2
GO:0010460     2
GO:0048745     2
GO:0030534     4
GO:0048754     4
GO:0010951     4
GO:1903321     3
GO:1903706     6
GO:0021602     2
GO:0042634     2
GO:0050802     2
GO:0010717     3
GO:0046888     3
GO:0070482     5
GO:0120162     3
GO:0030336     5
GO:0042749     2
GO:0048011     2
GO:0060317     2
GO:0071875     2
GO:2000178     2
GO:0010721     5
GO:0003002     6
GO:0043524     4
GO:0014910     3
GO:0008016     4
GO:0002028     3
GO:0032651     3
GO:0060395     3
GO:0002053     2
GO:0034260     2
GO:0040036     2
GO:0051085     2
GO:2000108     2
GO:0032872     4
GO:0090288     3
GO:0003156     2
GO:0008340     2
GO:0022410     2
GO:0070302     4
GO:0034767     4
GO:0010631     5
GO:2000146     5
GO:0090132     5
GO:0030308     4
GO:0052547     6
GO:0010614     2
GO:0045987     2
GO:0002790     5
GO:0019217     3
GO:0090130     5
GO:0043588     5
GO:0030307     4
GO:0098739     4
GO:1902107     4
GO:1903708     4
GO:0032611     3
GO:0046890     4
GO:0000132     2
GO:0010719     2
GO:0046686     2
GO:0034248     6
GO:0032890     3
GO:0055013     3
GO:0048562     5
GO:0001822     5
```

```
#Selecting relevant terms 
relevant1 <- c("negative regulation of MAPK cascade", 'learning or memory', 'transmembrane receptor protein serine/threonine kinase signaling pathway', 'ERK1 and ERK2 cascade', 'regulation of transcription from RNA polymerase II promoter in response to stress', 'regulation of cellular response to growth factor stimulus', 'response to transforming growth factor beta', 'p38MAPK cascade', 'positive regulation of catecholamine secretion', 'positive regulation of miRNA metabolic process', 'chemical synaptic transmission, postsynaptic', 'regulation of nervous system process', 'regulation of postsynaptic membrane potential', 'negative regulation of synaptic transmission', 'cellular response to calcium ion', ' myeloid cell differentiation', 'neurotrophin signaling pathway', 'vasculogenesis', 'response to peptide hormone', 'neuron apoptotic process', 'regulation of glucocorticoid receptor signaling pathway', 'cell migration involved in sprouting angiogenesis','neuron death')


#Plot
bpplot <- barplot(bp4, showCategory = relevant1, title= "GO: Biological Process")
bpplot
```

```
png("bp.treatment5.png", res= 250, width= 2000, height= 2750)
print(bpplot)


#KEGG PATHWAYS
KEGG2 <- enrichKEGG(dlist4, organism= 'mmu')
```

```
Reading KEGG annotation online: "https://rest.kegg.jp/link/mmu/pathway"...
```

```
Reading KEGG annotation online: "https://rest.kegg.jp/list/pathway/mmu"...
```

```
kegg2 <- as.data.frame(KEGG2)
k <- mutate(KEGG2, Description = gsub(" - Mus musculus [(]house mouse)", "", Description)) %>% 
barplot(showCategory = 20, title= "KEGG Pathways", xlab= "Gene Counts" )
k
png("kegg.treatment5.png", res= 250, height= 2000, width= 2000)
print(k)
kegg2
```

```
               ID
mmu04010 mmu04010
mmu05031 mmu05031
mmu04380 mmu04380
mmu05210 mmu05210
mmu04657 mmu04657
mmu05030 mmu05030
mmu04668 mmu04668
mmu04923 mmu04923
mmu04210 mmu04210
mmu05140 mmu05140
mmu05162 mmu05162
                                                                Description
mmu04010                MAPK signaling pathway - Mus musculus (house mouse)
mmu05031                 Amphetamine addiction - Mus musculus (house mouse)
mmu04380            Osteoclast differentiation - Mus musculus (house mouse)
mmu05210                     Colorectal cancer - Mus musculus (house mouse)
mmu04657               IL-17 signaling pathway - Mus musculus (house mouse)
mmu05030                     Cocaine addiction - Mus musculus (house mouse)
mmu04668                 TNF signaling pathway - Mus musculus (house mouse)
mmu04923 Regulation of lipolysis in adipocytes - Mus musculus (house mouse)
mmu04210                             Apoptosis - Mus musculus (house mouse)
mmu05140                         Leishmaniasis - Mus musculus (house mouse)
mmu05162                               Measles - Mus musculus (house mouse)
         GeneRatio  BgRatio       pvalue     p.adjust       qvalue
mmu04010     11/42 301/9334 5.734180e-08 7.741143e-06 6.156698e-06
mmu05031      4/42  69/9334 2.477134e-04 1.463554e-02 1.163996e-02
mmu04380      5/42 135/9334 3.252343e-04 1.463554e-02 1.163996e-02
mmu05210      4/42  88/9334 6.281542e-04 2.089349e-02 1.661704e-02
mmu04657      4/42  93/9334 7.738329e-04 2.089349e-02 1.661704e-02
mmu05030      3/42  48/9334 1.272882e-03 2.863985e-02 2.277789e-02
mmu04668      4/42 115/9334 1.705995e-03 3.290133e-02 2.616714e-02
mmu04923      3/42  57/9334 2.093742e-03 3.533189e-02 2.810022e-02
mmu04210      4/42 136/9334 3.142999e-03 4.714498e-02 3.749542e-02
mmu05140      3/42  70/9334 3.761550e-03 4.854601e-02 3.860970e-02
mmu05162      4/42 145/9334 3.955601e-03 4.854601e-02 3.860970e-02
                                                                      geneID
mmu04010 23882/15370/14281/19252/67603/240672/16476/12064/319520/58226/15511
mmu05031                                             11838/14281/14282/16476
mmu04380                                       14281/16477/14282/16476/14284
mmu05210                                            23882/14281/170770/16476
mmu04657                                             14281/19225/14282/16476
mmu05030                                                   14282/16476/12064
mmu04668                                             14281/16477/19225/16476
mmu04923                                                  19225/11554/384783
mmu04210                                            23882/14281/170770/16476
mmu05140                                                   14281/19225/16476
mmu05162                                            14281/170770/16476/15511
         Count
mmu04010    11
mmu05031     4
mmu04380     5
mmu05210     4
mmu04657     4
mmu05030     3
mmu04668     4
mmu04923     3
mmu04210     4
mmu05140     3
mmu05162     4
```

```
write.csv(kegg2, "kegg.csv")
```

```
#Disease ontology
library(clusterProfiler)
library(AnnotationDbi)
library(org.Mm.eg.db)
library(enrichplot)
library(MeSHDbi)
```

```
MeSH-related packages (MeSH.XXX.eg.db, MeSH.db, MeSH.AOR.db, and MeSH.PCR.db) are deprecated since Bioconductor 3.14. Use AnnotationHub instead. For details, check the vignette of MeSHDbi
```

```
library(AnnotationHub)
```

```
Loading required package: BiocFileCache
```

```
Loading required package: dbplyr
```

```
Attaching package: 'AnnotationHub'
```

```
The following object is masked from 'package:Biobase':

    cache
```

```
library(ggplot2)

#Loading database
ah <- AnnotationHub(localHub=FALSE)
ah.mmu <- query(ah, c("MeSHDb", "Mus musculus"))
file_mmu <- ah.mmu[["AH107113"]]
```

```
loading from cache
```

```
db <- MeSHDbi::MeSHDb(file_mmu)

#CONVERTING GENE ID's 
dlist3 <- bitr(genes4, fromType = "ENSEMBL", toType= 'ENTREZID', OrgDb = org.Mm.eg.db)
```

```
'select()' returned 1:1 mapping between keys and columns
```

```
Warning in bitr(genes4, fromType = "ENSEMBL", toType = "ENTREZID", OrgDb =
org.Mm.eg.db): 20.91% of input gene IDs are fail to map...
```

```
dlist4 <- dlist3$ENTREZID
dlist4
```

```
 [1] "225872"    "13656"     "387150"    "11838"     "23882"     "227885"   
 [7] "16007"     "80981"     "100504231" "15370"     "56501"     "100142658"
[13] "14281"     "13654"     "19725"     "16477"     "620695"    "12227"    
[19] "19225"     "320019"    "19735"     "14282"     "70924"     "225631"   
[25] "17684"     "19252"     "71198"     "330301"    "13653"     "67603"    
[31] "18167"     "15936"     "19366"     "105732"    "11554"     "270110"   
[37] "229599"    "53324"     "14313"     "211770"    "81489"     "105734"   
[43] "240672"    "56380"     "170770"    "68895"     "69944"     "16476"    
[49] "18387"     "18030"     "232685"    "12064"     "235627"    "73747"    
[55] "18109"     "24064"     "27220"     "19736"     "223775"    "14284"    
[61] "11910"     "320563"    "22057"     "319520"    "20556"     "56405"    
[67] "99929"     "74194"     "18124"     "20393"     "241528"    "14748"    
[73] "58226"     "237387"    "108797"    "24063"     "230809"    "70611"    
[79] "16600"     "20677"     "17691"     "18626"     "74476"     "384783"   
[85] "15511"     "13805"     "15476"
```

```
#DISEASE RESULTS
library(meshes)
```

```
meshes v1.26.0  

If you use meshes in published research, please cite the most appropriate paper(s):

Guangchuang Yu. Using meshes for MeSH term enrichment and semantic analyses. Bioinformatics 2018, 34(21):3766-3767, doi:10.1093/bioinformatics/bty410
```

```
ora.mesh.results2 <- enrichMeSH( # 226 obs
  dlist4, # vector of DEGs
  MeSHDb = db,
  database="gene2pubmed",
  pAdjustMethod = "fdr",
  pvalueCutoff = 0.05,
  qvalueCutoff = 0.05,
  category = "C")
```

```
loading from cache
```

```
ddf2 <- as.data.frame(ora.mesh.results2)
disease_list2= ddf2$Description
disease_list2
```

```
  [1] "Pregnancy Complications"                                 
  [2] "Hypertension, Pulmonary"                                 
  [3] "Pulmonary Atelectasis"                                   
  [4] "Carotid Artery Injuries"                                 
  [5] "Cholestasis, Intrahepatic"                               
  [6] "Status Epilepticus"                                      
  [7] "Endotoxemia"                                             
  [8] "Diabetic Neuropathies"                                   
  [9] "Hyperoxia"                                               
 [10] "Peripheral Nervous System Diseases"                      
 [11] "Pruritus"                                                
 [12] "Weight Loss"                                             
 [13] "Chronic Pain"                                            
 [14] "Respiratory Distress Syndrome, Newborn"                  
 [15] "Eye Injuries"                                            
 [16] "Abnormalities, Drug-Induced"                             
 [17] "Idiopathic Pulmonary Fibrosis"                           
 [18] "Respiratory Insufficiency"                               
 [19] "Cocaine-Related Disorders"                               
 [20] "Hyperphagia"                                             
 [21] "Scleroderma, Systemic"                                   
 [22] "Diabetic Angiopathies"                                   
 [23] "Prenatal Exposure Delayed Effects"                       
 [24] "Renal Insufficiency, Chronic"                            
 [25] "Hyperlipidemias"                                         
 [26] "Teratoma"                                                
 [27] "Rett Syndrome"                                           
 [28] "Carotid Artery Diseases"                                 
 [29] "Sleep Deprivation"                                       
 [30] "Constriction, Pathologic"                                
 [31] "Neointima"                                               
 [32] "Bronchial Hyperreactivity"                               
 [33] "Ischemic Attack, Transient"                              
 [34] "Intracranial Hemorrhages"                                
 [35] "Fever"                                                   
 [36] "Pulmonary Disease, Chronic Obstructive"                  
 [37] "Ventilator-Induced Lung Injury"                          
 [38] "Muscular Dystrophy, Animal"                              
 [39] "Radiation Injuries, Experimental"                        
 [40] "Dyskinesia, Drug-Induced"                                
 [41] "Diabetic Retinopathy"                                    
 [42] "Gonadal Dysgenesis, 46,XY"                               
 [43] "Neuralgia"                                               
 [44] "Thymus Neoplasms"                                        
 [45] "Corneal Diseases"                                        
 [46] "Crohn Disease"                                           
 [47] "Osteosclerosis"                                          
 [48] "Shock, Septic"                                           
 [49] "Osteosarcoma"                                            
 [50] "Substance Withdrawal Syndrome"                           
 [51] "Prostatic Intraepithelial Neoplasia"                     
 [52] "Escherichia coli Infections"                             
 [53] "Cocarcinogenesis"                                        
 [54] "Gastrointestinal Diseases"                               
 [55] "Cardiovascular Diseases"                                 
 [56] "Epilepsy, Absence"                                       
 [57] "Dehydration"                                             
 [58] "Hypercholesterolemia"                                    
 [59] "Bronchopulmonary Dysplasia"                              
 [60] "Leukocyte Disorders"                                     
 [61] "Skin Diseases"                                           
 [62] "Craniosynostoses"                                        
 [63] "Coronary Occlusion"                                      
 [64] "Aortic Aneurysm, Abdominal"                              
 [65] "Brain Infarction"                                        
 [66] "Morphine Dependence"                                     
 [67] "Albuminuria"                                             
 [68] "Splenic Diseases"                                        
 [69] "Uterine Neoplasms"                                       
 [70] "Metabolic Syndrome"                                      
 [71] "Anovulation"                                             
 [72] "Hand Deformities"                                        
 [73] "Gram-Positive Bacterial Infections"                      
 [74] "Colitis, Ulcerative"                                     
 [75] "Dysbiosis"                                               
 [76] "Choriocarcinoma"                                         
 [77] "Endocardial Cushion Defects"                             
 [78] "Liver Failure, Acute"                                    
 [79] "Leukemia, B-Cell"                                        
 [80] "Chlamydophila Infections"                                
 [81] "Chemical and Drug Induced Liver Injury, Chronic"         
 [82] "Optic Nerve Injuries"                                    
 [83] "Somatosensory Disorders"                                 
 [84] "Liver Cirrhosis, Alcoholic"                              
 [85] "Airway Remodeling"                                       
 [86] "Precancerous Conditions"                                 
 [87] "Hearing Loss, Sensorineural"                             
 [88] "Heart Diseases"                                          
 [89] "Hepatitis"                                               
 [90] "MPTP Poisoning"                                          
 [91] "Pseudomonas Infections"                                  
 [92] "Cell Transformation, Viral"                              
 [93] "Atrial Fibrillation"                                     
 [94] "Staphylococcal Infections"                               
 [95] "Nephrosis"                                               
 [96] "Cerebellar Neoplasms"                                    
 [97] "Bone Diseases, Metabolic"                                
 [98] "Irritable Bowel Syndrome"                                
 [99] "Medulloblastoma"                                         
[100] "Toxoplasmosis"                                           
[101] "Malaria, Cerebral"                                       
[102] "Heroin Dependence"                                       
[103] "Bone Neoplasms"                                          
[104] "Neoplasm, Residual"                                      
[105] "Leukemia, Myelogenous, Chronic, BCR-ABL Positive"        
[106] "Encephalitis"                                            
[107] "Carcinoma, Embryonal"                                    
[108] "Lymphocytic Choriomeningitis"                            
[109] "Sciatic Neuropathy"                                      
[110] "Hypothermia"                                             
[111] "Heart Septal Defects, Atrial"                            
[112] "Fetal Diseases"                                          
[113] "Parkinson Disease, Secondary"                            
[114] "Binge Drinking"                                          
[115] "Familial Primary Pulmonary Hypertension"                 
[116] "Renal Artery Obstruction"                                
[117] "Lipodystrophy"                                           
[118] "Primary Myelofibrosis"                                   
[119] "Vascular Diseases"                                       
[120] "Lymphatic Metastasis"                                    
[121] "Keratoacanthoma"                                         
[122] "Vascular System Injuries"                                
[123] "Pheochromocytoma"                                        
[124] "Polydactyly"                                             
[125] "Hypovolemia"                                             
[126] "Carotid Artery Thrombosis"                               
[127] "Myelodysplastic Syndromes"                               
[128] "Pneumonia, Bacterial"                                    
[129] "Peripheral Nerve Injuries"                               
[130] "Intestinal Neoplasms"                                    
[131] "Ureteral Obstruction"                                    
[132] "Parkinsonian Disorders"                                  
[133] "Head and Neck Neoplasms"                                 
[134] "Adrenal Gland Neoplasms"                                 
[135] "Plaque, Amyloid"                                         
[136] "Wounds and Injuries"                                     
[137] "Pneumonia, Mycoplasma"                                   
[138] "Carcinoma, Ductal"                                       
[139] "Ovotesticular Disorders of Sex Development"              
[140] "Aortic Diseases"                                         
[141] "Colorectal Neoplasms, Hereditary Nonpolyposis"           
[142] "Helicobacter Infections"                                 
[143] "Neuroendocrine Tumors"                                   
[144] "Hemorrhage"                                              
[145] "Precursor T-Cell Lymphoblastic Leukemia-Lymphoma"        
[146] "Tachycardia, Ventricular"                                
[147] "Facial Pain"                                             
[148] "Hyperinsulinism"                                         
[149] "Hypogonadism"                                            
[150] "Learning Disabilities"                                   
[151] "Retinal Neovascularization"                              
[152] "Adenomatous Polyposis Coli"                              
[153] "Alcoholism"                                              
[154] "Immune System Diseases"                                  
[155] "Arthritis, Psoriatic"                                    
[156] "Multiple Organ Failure"                                  
[157] "Abortion, Habitual"                                      
[158] "Liposarcoma"                                             
[159] "Cerebellar Ataxia"                                       
[160] "DiGeorge Syndrome"                                       
[161] "Intestinal Polyps"                                       
[162] "Keratitis, Herpetic"                                     
[163] "Aneurysm"                                                
[164] "Amnesia"                                                 
[165] "Epilepsy, Tonic-Clonic"                                  
[166] "Lymphoma, Large-Cell, Anaplastic"                        
[167] "Urogenital Abnormalities"                                
[168] "Tongue Neoplasms"                                        
[169] "Hypertrophy, Left Ventricular"                           
[170] "Hand Deformities, Congenital"                            
[171] "Neoplasms, Germ Cell and Embryonal"                      
[172] "Sleep Apnea Syndromes"                                   
[173] "Carcinoma, Non-Small-Cell Lung"                          
[174] "Postoperative Complications"                             
[175] "Lentivirus Infections"                                   
[176] "Hypoglycemia"                                            
[177] "Hypoparathyroidism"                                      
[178] "Heat Stress Disorders"                                   
[179] "Anemia, Sickle Cell"                                     
[180] "Vesico-Ureteral Reflux"                                  
[181] "Fatigue"                                                 
[182] "Pain, Postoperative"                                     
[183] "Anorexia"                                                
[184] "Death"                                                   
[185] "Thromboembolism"                                         
[186] "Cholestasis"                                             
[187] "Endomyocardial Fibrosis"                                 
[188] "Edema"                                                   
[189] "Papilloma"                                               
[190] "Photosensitivity Disorders"                              
[191] "Vitreoretinopathy, Proliferative"                        
[192] "Coronary Restenosis"                                     
[193] "Keratosis"                                               
[194] "Retinal Diseases"                                        
[195] "Drug Eruptions"                                          
[196] "Hypertension, Portal"                                    
[197] "Rhabdoviridae Infections"                                
[198] "Calcinosis"                                              
[199] "Infertility"                                             
[200] "Hypotension"                                             
[201] "Herpes Simplex"                                          
[202] "Radiation Injuries"                                      
[203] "Scrapie"                                                 
[204] "Carcinoma, Pancreatic Ductal"                            
[205] "Hypertrophy, Right Ventricular"                          
[206] "Ellis-Van Creveld Syndrome"                              
[207] "Encephalitis, Herpes Simplex"                            
[208] "Epilepsy, Temporal Lobe"                                 
[209] "Hydrocephalus"                                           
[210] "Postmortem Changes"                                      
[211] "Anhedonia"                                               
[212] "Vascular Calcification"                                  
[213] "Diastema"                                                
[214] "Polycythemia Vera"                                       
[215] "Mouth Neoplasms"                                         
[216] "Stomach Ulcer"                                           
[217] "Primary Ovarian Insufficiency"                           
[218] "Leukemia, Lymphoid"                                      
[219] "Tremor"                                                  
[220] "Periodontitis"                                           
[221] "Neuromuscular Diseases"                                  
[222] "Dilatation, Pathologic"                                  
[223] "Carcinoma, Small Cell"                                   
[224] "Facial Nerve Injuries"                                   
[225] "Cryptorchidism"                                          
[226] "Prostatic Neoplasms, Castration-Resistant"               
[227] "Leukemia, Lymphocytic, Chronic, B-Cell"                  
[228] "Movement Disorders"                                      
[229] "Cerebral Infarction"                                     
[230] "Thyroid Neoplasms"                                       
[231] "Influenza, Human"                                        
[232] "Tooth, Supernumerary"                                    
[233] "Fractures, Bone"                                         
[234] "Jaw Abnormalities"                                       
[235] "Peripheral Arterial Disease"                             
[236] "Autonomic Nervous System Diseases"                       
[237] "Hyperaldosteronism"                                      
[238] "Synostosis"                                              
[239] "Wounds, Penetrating"                                     
[240] "Leukemia, Erythroblastic, Acute"                         
[241] "Blast Crisis"                                            
[242] "Tibial Fractures"                                        
[243] "Lipid Metabolism, Inborn Errors"                         
[244] "Obstetric Labor, Premature"                              
[245] "Arrhythmias, Cardiac"                                    
[246] "Arteriovenous Malformations"                             
[247] "Haemophilus Infections"                                  
[248] "Kidney Neoplasms"                                        
[249] "Hereditary Central Nervous System Demyelinating Diseases"
```

```
write.csv(ddf2, "diseaselist.csv")

#23 

relevant <- c("Rett Syndrome", "Intracranial Hemorrhages", 'Cocaine-Related Disorders', 'MPTP Poisoning (related to parkinsons)', 'Ischemic Attack, Transient', 'Status Epilepticus', 'Somatosensory Disorders', 'Substance Withdrawal Syndrome', 'Encephalitis', 'Parkinson Disease, Secondary', 'Parkinsonian Disorders', 'Brain Infarction', 'Epilepsy, Absence', 'Movement Disorders', 'Morphine Dependence', 'Aneurysm', 'Amnesia', 'Epilepsy, Tonic-Clonic', 'Cerebral Hemorrhage', 'Learning Disabilities', 'Cerebral Infarction', 'Alcoholism', 'Hypoxia-Ischemia, Brain') 

#DOT PLOT DISEASES

diseaseplot <- barplot(ora.mesh.results2, showCategory = relevant, title= "Brain-related Diseases Enrichment")

png("disease2.png", res= 250, height= 3000, width= 2000)

print(diseaseplot)
```

#### Error
